# GENECraft: A Function- and Prior-Constrained Discrete Dynamical System for Controllable Regulatory DNA Design

**DOI:** 10.64898/2026.11.16.767190

**Authors:** Jue Wang, Tongyue Xu, Cheng Tan, Wenqi Zhong, Guangyu Guo, Ling Zhang, Zhongwei Qiu, Stan Z. Li

## Abstract

Designing regulatory DNA requires reconciling two complementary paradigms: genomic language models (gLMs) capture evolutionary syntax without guaranteeing targeted activity, whereas sequence-to-function predictors provide explicit functional supervision without enforcing native-like sequence plausibility. We present **GENECraft**, a training-free Function- and Prior-Constrained Discrete Dynamical System (FP-DDS) that unifies a gLM prior (Carbon) and a functional oracle (AlphaGenome) for controllable inference-time sequence synthesis. Rather than relying on unconditional sampling or post-hoc filtering, GENECraft evolves continuous latent nucleotide logits while evaluating strictly decoded discrete DNA sequences within a time-dependent composite vector field. To bridge heterogeneous tokenizations without trainable adapters, we introduce a parameter-free nucleotide-to-6-mer relaxation and harmonize functional gradients, bidirectional contextual prior guidance, and differentiable compositional penalties via root-mean-square (RMS) normalized logit pullbacks. The non-monotonic discrete trajectory concludes with a feasibility-first readout that prioritizes constraint-satisfying states before maximizing functional utility. Extensive evaluations across 2,185 human loci spanning promoters, multi-constraint enhancers, and 18 transcription factors demonstrate that GENECraft achieves 62.70%, 65.23%, and 94.56% full-joint success rates, outperforming seven state-of-the-art baselines by large margins while significantly improving native compositional similarity and perturbation stability.

## 1. Introduction

*De novo* design of regulatory DNA elements, such as promoters, enhancers, and transcription factor (TF) binding sites, is foundational to synthetic biology and precision gene therapy [2, 18, 36, 40]. Achieving precise transcriptional control requires synthesizing sequences that not only drive strong cell-type-specific activity within long-range genomic contexts, but also obey strict biochemical composition rules and avoid off-target chromatin activation. Navigating this high-dimensional, discrete nucleotide space under tightly coupled functional and structural constraints remains a major computational challenge.

Recent advances in genomic foundation models offer two complementary yet largely disconnected paradigms to tackle this problem (Figure 1). On the generative side, large-scale genomic language models (gLMs) pretrained on massive sequence corpora effectively distill evolutionary syntax and local *k*-mer dependencies [1, 10, 12, 24, 33]. However, because unconditional gLM priors are agnostic to downstream assay readouts, they lack explicit control over cell-type-specific regulatory potency [17, 25]. Consequently, relying on generative sampling with post-hoc filtering becomes highly sample-inefficient when multiple functional and compositional constraints must be satisfied simultaneously.

**Figure 1.**
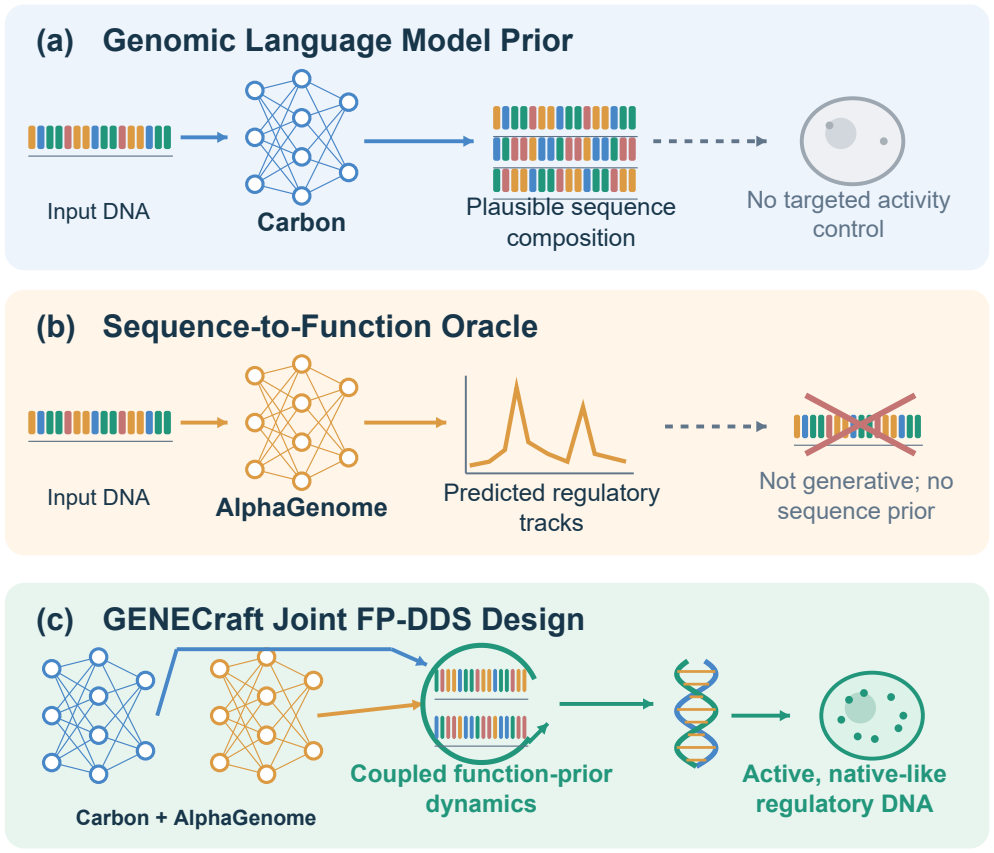
Conceptual comparison of regulatory DNA design paradigms. **(a)** Genomic language models (e.g., Carbon [1]) capture evolutionary syntax but lack explicit functional control. **(b)** Sequence-to-function oracles (e.g., AlphaGenome [7]) provide functional gradients but fail to enforce sequence plausibility. **(c)** GENECraft unifies functional guidance and prior regularization inside an inference-time discrete dynamical system.

Conversely, discriminative sequence-to-function predictors (oracles) directly map DNA sequences to multi-track epigenetic and transcriptional profiles, providing explicit gradients for sequence optimization [5–7, 31, 43]. Yet, discriminative oracles are not standalone generators and lack an intrinsic notion of evolutionary sequence plausibility. Optimizing oracle predictions in isolation, whether via gradient ascent, evolutionary search, or reward finetuning, readily exploits out-of-distribution predictor pathologies [11]. This adversarial over-optimization frequently yields unnatural GC extremes, homopolymer artifacts, and fragile motifs that fail to generalize across independent evaluators or mutational perturbations [3, 15, 22].

To unify these complementary strengths without task-specific retraining, we introduce **GENECraft**, a *Function- and Prior-Constrained Discrete Dynamical System* (FP-DDS; Figure 1c). Rather than treating generation and evaluation as separate stages, GENECraft evolves continuous latent nucleotide logits that deterministically decode discrete DNA sequences via a fixed-noise straight-through interface within native flanking context. To reconcile heterogeneous model representations at each step, we design a parameter-free nucleotide-to-6-mer relaxation bridge that extracts exact-evaluated bidirectional contextual gradients from a frozen gLM (Carbon) and aligns them with base-resolution functional gradients from a frozen oracle (AlphaGenome), along-side differentiable sequence-quality penalties. Crucially, we harmonize these competing forces in logit space using root-mean-square (RMS) normalized softmax Jacobian pullbacks, evolving the latent state under a linear step-size ramp that transitions from broad early exploration to late-stage motif consolidation. Because discrete trajectories over rugged multi-constraint landscapes are inherently non-monotonic, FP-DDS concludes with a feasibility-first readout over all visited states, prioritizing feasible candidates before maximizing functional utility.

Our primary contributions are three-fold:

- **FP-DDS Framework:** We cast inference-time regulatory DNA synthesis as a training-free discrete dynamical system that couples functional oracles and genomic priors via a parameter-free token bridge, RMS-normalized logit pullbacks, and feasibility-first trajectory readout.
- **High-Plausibility Functional Synthesis:** GENECraft synthesizes regulatory DNA uniting high target activity with strict compositional compliance and enhanced stability under mutational and contextual perturbations.
- **State-of-the-Art Performance:** Across 2,185 human regulatory loci and seven baselines, GENECraft achieves 62.70% promoter, 65.23% enhancer, and 94.56% TF macro full-joint success under independent Enformer and Borzoi cross-oracle validation.

## 2. Related Work

### Genomic Foundation Models

Genomic language models have progressed from representation learners such as DNABERT and Nucleotide Transformer [9, 12, 24] to long-context generative priors including Evo, Evo2, GENERator, Carbon, and regLM [1, 10, 28, 33, 42]. Whereas prior work-flows typically use these models for unconditional sampling or post-hoc filtering, GENECraft turns a gLM into an active, step-wise guidance signal during optimization.

### Regulatory Predictors and Readouts

Sequence-to-function models span local chromatin predictors [6, 43] to long-range transcriptional models such as Basenji, Enformer, Borzoi, and unified multi-assay AlphaGenome [5, 7, 26, 31], alongside quantitative reporter assays [3, 13, 22]. We leverage this ecosystem both to formulate multi-constraint objectives and to evaluate cross-oracle generalization with independent Enformer and Borzoi readouts.

### Functional Sequence Design

Existing design approaches encompass predictor-guided discrete search [38], straight-through optimization in Fast SeqProp [30], prior-regularized search [11], and reward-guided generative or diffusion frame-works [14, 18, 41, 44]. GENECraft advances this landscape through an inference-time FP-DDS controller featuring a parameter-free nucleotide/6-mer gradient bridge, synchronized bidirectional gLM and functional guidance, RMS-normalized pullbacks, and feasibility-first readout.

## 3. Method

### 3.1. Task Definition and Overview

Given fixed flanking genomic context *c*, a central editable region of length *L*, and a target regulatory specification *t* with explicit sequence-quality constraints, our goal is to synthesize a discrete DNA sequence *x*^*\**^ ∈ *{*A, C, G, T*}*^*L*^ that achieves strong regulatory activity while preserving native-like composition (Figure 2a). We formulate inference-time design as a Function- and Prior-Constrained Discrete Dynamical System (FP-DDS) evolving over discrete steps *s* ∈ *{*1, …, *S}* (Figure 2b) followed by a feasibility-first trajectory readout (Figure 2c). Both AlphaGenome and Carbon operate with frozen parameters: AlphaGenome defines the functional utility *U*_AG_(*x*; *c, t*) and binary generation-stage feasibility indicator *F* (*x*; *c, t*), whereas Carbon provides the contextual log-likelihood *L*_*C*_(*x*; *c*).

**Figure 2.**
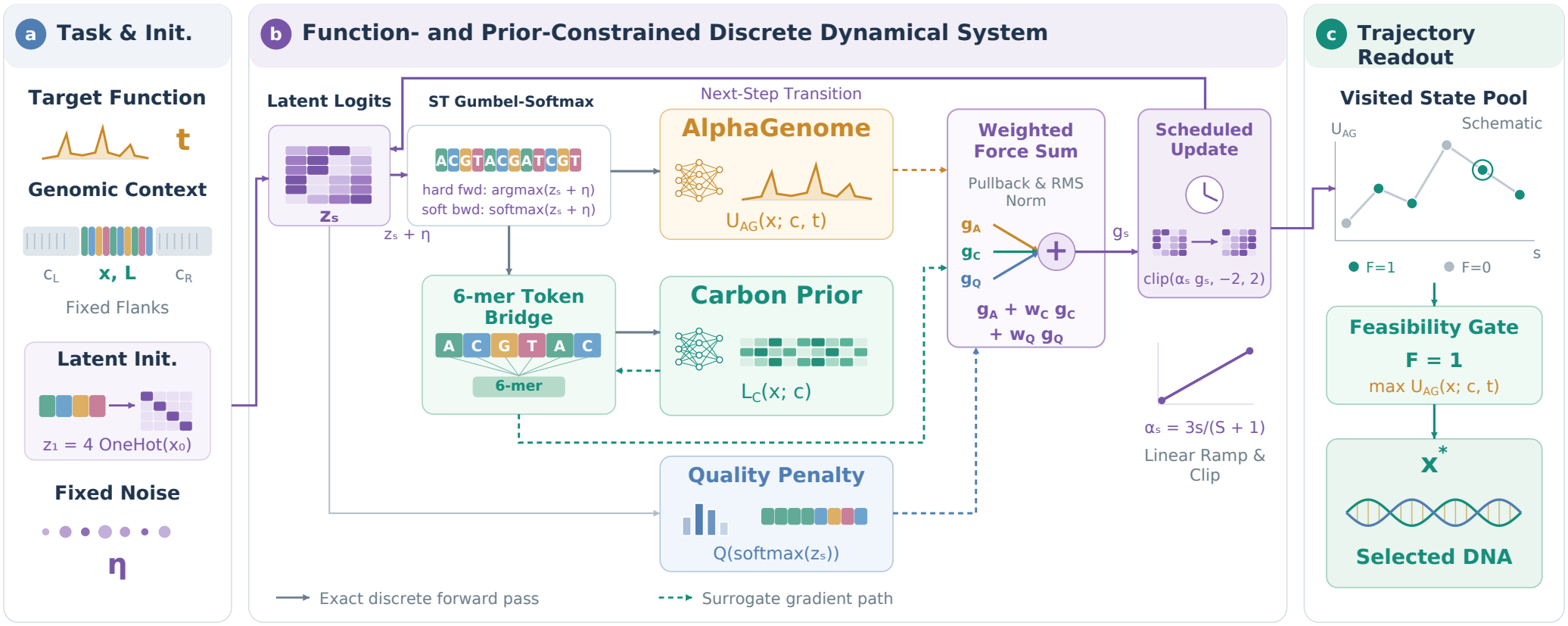
GENECraft architecture. **(a)** Latent logits initialize with fixed Gumbel noise *η*. **(b)** Fixed-noise ST Gumbel-Softmax couples hard forward arg max decoding on a shared discrete DNA sequence with soft backward softmax pullbacks across the 6-mer bridge to harmonize RMS-normalized AlphaGenome and Carbon gradients. **(c)** Feasibility-first readout selects the highest-utility constraint-compliant state.

Each optimization restart initializes a random discrete sequence *x*_0_ (see Appendix B for native initialization), continuous latent logits *z*_1_ = 4 OneHot(*x*_0_) ∈ ℝ^*L×*4^, and a fixed Gumbel perturbation *η* ∈ ℝ^*L×*4^. Both *x* and *z*_*s*_ span only the editable target *L*; model-specific adapters concatenate *x* with fixed context *c* for full-context forward evaluation while restricting updates to the target logits.

The utility *U*_AG_(*x*; *c, t*) encodes class-specific objectives: maximizing transcription start site (TSS) CAGE signal for promoters, and target ChIP-seq binding tracks for transcription factor (TF) elements. For enhancers, it jointly rewards chromatin accessibility (DNase) and enhancer histone marks (H3K27ac, H3K4me1) while suppressing off-target transcription initiation (CAGE) and promoter H3K4me3 marks.^1^

### 3.2. Sequence Representation and Gradient Interface

At each step *s*, latent logits *z*_*s*_ and fixed Gumbel noise *η* form a unit-temperature straight-through (ST) Gumbel-Softmax interface [8, 23, 32] coupling discrete evaluation with joint functional and prior optimization. The forward pass deterministically decodes a hard sequence via *x*_*i*_ = arg max_*b*_(*z*_*s,ib*_ + *η*_*ib*_) without differentiating through arg max, while the backward pass routes gradients through the soft surrogate *p* = softmax(*z*_*s*_ + *η*) via *P*_ST_ = sg(OneHot(*x*) *− p*) + *p*, with stop-gradient operator sg(*·*). Holding *η* fixed across steps ensures all models evaluate the exact same discrete sequence *x* and transitions reflect directed logit updates.

Although AlphaGenome and Carbon consume distinct input representations, their nucleotide-level gradients *G* ∈ ℝ^*L×*4^ are mapped onto the shared logit space via the softmax Jacobian pullback:

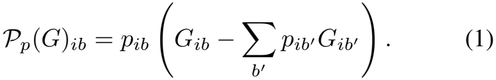

Whereas AlphaGenome operates on base-resolution one-hot encodings to yield *G*_*A*_, Carbon tokenizes DNA into 4096 6-mer IDs. To bridge these representations without trainable adapters, we introduce a parameter-free nucleotide-to-6-mer relaxation. Given nucleotide probabilities *P* ∈ [0, 1]^*L×*4^, each 6-mer block *j* induces a factorized distribution 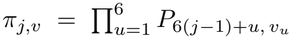 over the DNA token subset, forming expected embeddings *e*_*j*_ = ∑_*v*_ *π*_*j,v*_ *E*_*v*_ from Carbon’s embedding rows *E*_*v*_, with trailing bases (*L* mod 6 *>* 0) scored by prefix marginalization. Evaluated at *P* = OneHot(*x*), this forward relaxation matches the exact discrete Carbon log-likelihood; differentiation with respect to *P* yields a surrogate gradient *G*_*C*_, averaging forward and reverse-complement passes to enforce strand symmetry (Appendix A.1).

### 3.3. Joint Dynamics and Scheduled Updates

Latent state evolution is governed by a composite vector field that balances functional utility, evolutionary syntax, and compositional quality via root-mean-square (RMS) normalization, *N* (*g*) = *g/* max(RMS(*g*), 10^*−*12^). Denoting the normalized logit-space forces as *g*_*A*_ = *N* (*P*_*p*_(*G*_*A*_)), *g*_*C*_ = *N* (*P*_*p*_(*G*_*C*_)), and *g*_*Q*_ = *N* (*∇*_*z*_*Q*), the joint update direction at step *s* is:

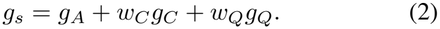

Applying RMS normalization after each pullback equalizes force scales across heterogeneous objectives prior to coordinate clipping, rendering *w*_*C*_ and *w*_*Q*_ interpretable relative weights.

Within this controller, bidirectional Carbon guidance (*w*_*C*_ = 0.1) acts as an in-loop prior regularizer that restrains surrogate over-optimization [11], steering trajectories away from adversarial optima toward native-like *k*-mer composition. Simultaneously, the differentiable penalty *Q*(*z*) (*w*_*Q*_ = 0.15) is evaluated on unperturbed probabilities softmax(*z*_*s*_) to softly penalize extreme GC content and homopolymers, mitigating synthesis artifacts [37] (Appendix A).

The latent state *z*_*s*_ evolves under a linear step-size schedule with coordinate-wise clipping:

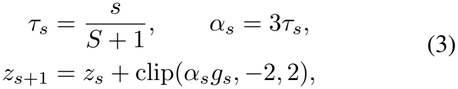

where *τ*_*s*_ indexes optimization progress across horizon *S*. This schedule applies conservative early updates to explore sequence space before amplifying late-stage steps to consolidate functional motifs, resetting across *R* restarts (*S* = 24, *R* = 1 for promoters/TFs; *S* = 48, *R* = 2 for enhancers; Appendix A).

### 3.4. Feasibility-First Trajectory Readout

Generation-stage feasibility *F* (*x*; *c, t*) ∈ *{*0, 1*}* couples AlphaGenome task criteria with the hard sequence-composition gate *I*_QC_, which bounds GC content, maximum single-base fraction, and homopolymer run length (Appendix E.2; Supplementary Table 8). Whereas *Q*(*z*) supplies smooth gradient guidance during evolution, *I*_QC_ enforces strict compliance at readout.

Because discrete trajectories over rugged landscapes are non-monotonic, FP-DDS concludes with a *feasibility-first* readout over all visited states across all restarts (Figure 2c). The readout filters for feasible states satisfying *F* (*x*; *c, t*) = 1 and selects *x*^*\**^ maximizing exact utility *U*_AG_(*x*; *c, t*); if no state satisfies *F* = 1, it returns the highest-utility sequence with its failure flag preserved (Algorithm 1, Appendix A). Generation and readout rely exclusively on AlphaGenome and *I*_QC_, reserving Enformer and Borzoi strictly for post-generation validation.

## 4. Experiments

### 4.1. Experimental Setup

#### Benchmark Cohort and Baselines

We evaluate GENECraft across 2,185 human regulatory loci from the ATGC-Gen benchmark [39]: 512 K562 promoters, 512 melanocyte enhancers, and 1,161 GM12878 binding elements across 18 TFs (equal-factor macro-averaged; Appendix E.1). We compare GENECraft against seven baselines under standard deployment protocols (Appendix N): gradient-based Fast SeqProp [30], evolutionary search AdaLead [38], dinucleotide shuffle, and generative models Carbon, Evo2, NTv3, and DRAKES [1, 9, 10, 41]. Fast SeqProp and AdaLead optimize *U*_AG_ directly, while generative baselines apply best-of-*N* AlphaGenome reranking (*N* = 64 for promoters/TFs; *N* = 128 for enhancers). We also evaluate an internal AlphaGenome-only (*w*_*C*_ = 0) baseline across all 2,185 loci (Supplementary Table 9). Every method produces one locked sequence per locus with all constraint failures retained.

#### Controlled Ablation and Modularity Protocols

To isolate the impact of in-loop Carbon guidance, we construct a stratified 336-locus cohort (96 promoters, 96 enhancers, 144 TFs) and evaluate *w*_*C*_ ∈ *{*0, 0.03, 0.1, 0.3*}* while holding all other components, initializations, and Gumbel noise strictly fixed (Supplementary Table 10B–C). We further dissect controller mechanics and backbone modularity on 24-locus cohorts (Table 4): (B) shared-component enhancer controls aligning objectives, gates, readout, and compute budgets across FP-DDS, constant step size (*α*_*s*_ = 1.5), and Fast SeqProp variants; (C) a five-arm enhancer component ablation; and (D) four modular oracle–prior pairings (AlphaGenome/Enformer *×* Carbon/Evo2) across 24 TF loci.

#### Evaluation Metrics

To prevent reward hacking of the optimization oracle, each candidate *i* is evaluated post-generation under a stringent multi-oracle conjunction termed *full-joint success*:

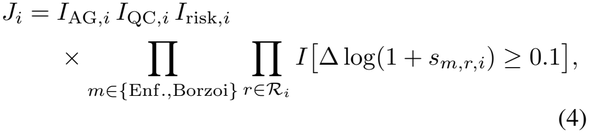

where *I*_AG,*i*_ and *I*_QC,*i*_ verify common AlphaGenome bench-mark screening and hard compositional compliance. The held-out validation set *R*_*i*_ comprises TSS CAGE for promoters, DNase and H3K27ac for enhancers, and target ChIP-seq for TFs, with Δ log(1 + *s*_*m,r,i*_) measuring predicted log-activity gain over the native sequence under external oracles *m* ∈ *{*Enformer, Borzoi*}*. For enhancers, *I*_risk,*i*_ enforces chromatin specificity in both external predictors by bounding CAGE peak, mean H3K4me3, and the H3K4me1/H3K4me3 ratio (Supplementary Table 7; *I*_risk,*i*_ *≡* 1 otherwise).

Methods are ranked by the primary evaluation block: full-joint success (*J*_*i*_), constituent pass rates, and external activity gains. The unranked diagnostic block reports conditional log-likelihoods (nats/bp) under Carbon and Evo2, descriptive composition (GC%, CpG *O/E*), constraint compliance, and native *k*-mer similarity via dinucleotide and 4-mer Jensen– Shannon (JS) divergences (bits) [29]. Accompanying native intervals (N) provide descriptive baselines rather than universal validity thresholds; these reference intervals contextualize reported cohort averages and do not replace per-sequence quality-control gates (Q). Protocol and statistical details appear in Appendices E.2, E.3, and F.

### 4.2. Comparison with State-of-the-Art Baselines

#### Promoter Design Comparison

In Table 1, GENECraft achieves a state-of-the-art **62.70%** full-joint success rate, outperforming the top gradient baseline, Fast SeqProp (45.31%), and generative baseline, Evo2 (5.27%). Unconditional priors (Carbon, Evo2) preserve native-like *k*-mer syntax but lack functional steering, limiting best-of-64 reranking to under 5.3% joint success. Conversely, Fast SeqProp and reward-finetuned DRAKES reach moderate activity pass rates (55.27% and 56.64%) by exploiting high-GC, repetitive sequences that inflate oracle scores while degrading compositional compliance (78.91% and 1.95%). By combining step-wise Carbon gradients with differentiable quality penalties, GENECraft confines optimization to syntactically plausible DNA, securing **100.00%** compositional compliance and the highest held-out Enformer (1.934) and Borzoi (0.515) CAGE gains.

**Table 1.** Promoter design benchmark (*n* = 512 K562 loci). Held-out Enformer and Borzoi oracles evaluate all designs with failures retained; primary metrics rank performance while diagnostics remain unranked. Bold and underline denote the best and second-best distinct displayed values per metric within each comparable panel, including ties; unranked reference diagnostics are excluded. Reference intervals denote descriptive native envelopes (N: P05–P95, or [0, P95] for JS_2_*/*JS_4_) and fixed quality-gate bounds (Q; Appendix E.2).

| **Primary Evaluation Metrics** (*n* = 512) |  |  |  |  |  |  |  |  |
| Metric | **GENECraft (Ours)** | Fast SeqProp [30] | AdaLead [38] | Dinucleotide shuffle | Carbon [1] | Evo2 [10] | NTv3 [9] | DRAKES [41] |
|---|---|---|---|---|---|---|---|---|
| Full joint (%) ↑ | **62.70** | 45.31 | 11.52 | 0.20 | 4.49 | 5.27 | 1.17 | 0.78 |
| Activity pass (%) ↑ | **62.70** | 55.27 | 15.23 | 1.95 | 29.49 | 33.40 | 4.30 | 56.64 |
| Quality pass (%) ↑ | **100.00** | 78.91 | 99.80 | 39.26 | 39.26 | 39.45 | 44.14 | 1.95 |
| Enf. CAGE ↑ | **1.934** | 1.400 | -0.180 | -1.269 | 0.168 | 0.165 | -1.270 | 1.168 |
| Borzoi CAGE ↑ | **0.515** | 0.447 | -0.463 | -0.858 | -0.010 | -0.023 | -0.809 | 0.447 |
| **Reference Sequence Diagnostics** |  |  |  |  |  |  |  |  |
| Metric | GENECraft | Fast SeqProp | AdaLead | Shuffle | Carbon | Evo2 | NTv3 | DRAKES |
| Carbon LL [-1.335, -0.718]<sup>N</sup> | -1.319 | -1.382 | -1.448 | -1.332 | -1.213 | -1.254 | -1.225 | -1.298 |
| Evo2 LL [-1.322, -0.680]<sup>N</sup> | -1.312 | -1.383 | -1.449 | -1.325 | -1.253 | -1.200 | -1.225 | -1.297 |
| Dinucleotide JS [0.0000, 0.1554]<sup>N</sup> | 0.0840 | 0.0909 | 0.0801 | 0.0000 | 0.0389 | 0.0377 | 0.1668 | 0.0539 |
| 4-mer JS [0.0000, 0.1398]<sup>N</sup> | 0.1117 | 0.1091 | 0.1076 | 0.0571 | 0.0797 | 0.0786 | 0.1438 | 0.0896 |
| GC content (%) [35, 65]<sup>Q</sup> | 53.90 | 56.24 | 50.47 | 66.48 | 65.60 | 66.47 | 37.67 | 72.19 |
| GC-bound pass (%) | 100.00 | 86.33 | 100.00 | 41.99 | 45.51 | 41.60 | 58.40 | 2.15 |
| CpG O/E [0.244, 1.125]<sup>N</sup> | 1.096 | 1.337 | 1.047 | 0.801 | 0.876 | 0.796 | 0.894 | 1.096 |
| Longest HP (bp) [1, 10]<sup>Q</sup> | 4.60 | 5.86 | 4.71 | 5.83 | 6.52 | 5.09 | 20.33 | 5.59 |
| Max. base fraction [0.25, 0.40]<sup>Q</sup> | 0.297 | 0.345 | 0.282 | 0.381 | 0.374 | 0.377 | 0.419 | 0.395 |

#### Enhancer Design Comparison

Enhancer synthesis must drive chromatin accessibility (DNase) and histone acetylation (H3K27ac) while suppressing promoter leakage (CAGE, H3K4me3). GENECraft achieves **65.23%** full-joint success, whereas external baselines collapse, with runner-up AdaLead at 2.15% (Table 2). Reward-finetuned DRAKES over-optimizes H3K27ac while incurring negative DNase gains and severe promoter leakage (0.20% risk pass). Conversely, discrete evolutionary search (AdaLead) preserves 100% sequence quality via local mutations but lacks multi-track gradient directionality, stalling at 3.71% activity pass. GENECraft succeeds because RMS-normalized logit pull-backs balance competing epigenetic updates while in-loop Carbon guidance discourages unnatural promoter-like CpG islands, jointly securing high activity (83.79%), risk suppression (72.27%), and quality compliance (100.00%).

**Table 2.**
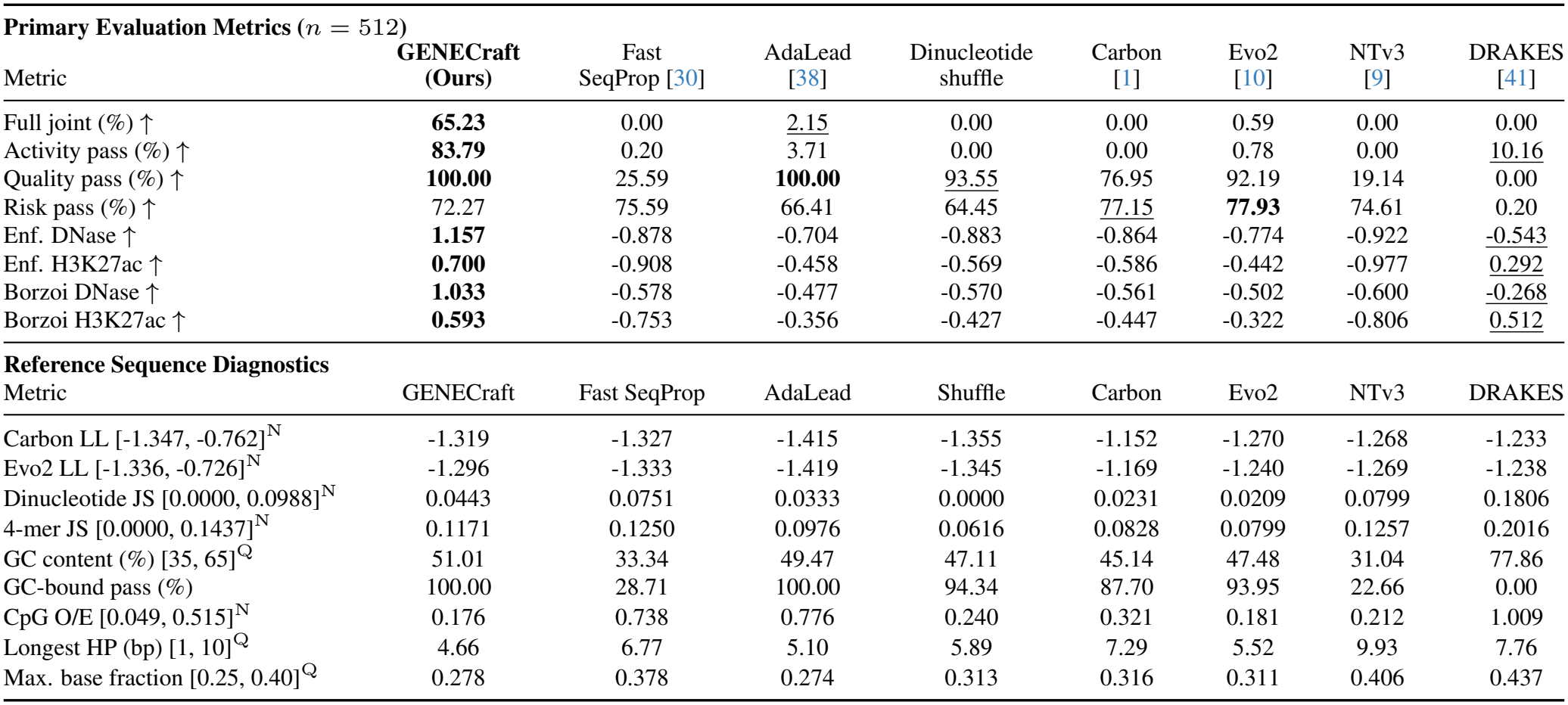
Comparison on the multi-constraint enhancer benchmark (*n* = 512 melanocyte loci). Primary metrics evaluate joint compliance across activity, sequence quality, and promoter-leakage risk gates alongside external Δ log(1 + *s*) DNase and H3K27ac gains. Reference interval conventions (N, Q) follow Table 1.

| **Primary Evaluation Metrics** (*n* = 512) |  |  |  |  |  |  |  |
|---|---|---|---|---|---|---|---|
| Metric | **GENECraft (Ours)** | Fast SeqProp [30] | AdaLead [38] | Dinucleotide shuffle | Carbon [1] | Evo2 [10] | NTv3 [9] | DRAKES [41] |
|---|---|---|---|---|---|---|---|---|
| Full joint (%) ↑ | **65.23** | 0.00 | 2.15 | 0.00 | 0.00 | 0.59 | 0.00 | 0.00 |
| Activity pass (%) ↑ | **83.79** | 0.20 | 3.71 | 0.00 | 0.00 | 0.78 | 0.00 | 10.16 |
| Quality pass (%) ↑ | **100.00** | 25.59 | **100.00** | 93.55 | 76.95 | 92.19 | 19.14 | 0.00 |
| Risk pass (%) ↑ | 72.27 | 75.59 | 66.41 | 64.45 | 77.15 | **77.93** | 74.61 | 0.20 |
| Enf. DNase ↑ | **1.157** | -0.878 | -0.704 | -0.883 | -0.864 | -0.774 | -0.922 | -0.543 |
| Enf. H3K27ac ↑ | **0.700** | -0.908 | -0.458 | -0.569 | -0.586 | -0.442 | -0.977 | 0.292 |
| Borzoi DNase ↑ | **1.033** | -0.578 | -0.477 | -0.570 | -0.561 | -0.502 | -0.600 | -0.268 |
| Borzoi H3K27ac ↑ | **0.593** | -0.753 | -0.356 | -0.427 | -0.447 | -0.322 | -0.806 | 0.512 |
| **Reference Sequence Diagnostics** |  |  |  |  |  |  |  |  |
| Metric | GENECraft | Fast SeqProp | AdaLead | Shuffle | Carbon | Evo2 | NTv3 | DRAKES |
| Carbon LL [-1.347, -0.762]<sup>N</sup> | -1.319 | -1.327 | -1.415 | -1.355 | -1.152 | -1.270 | -1.268 | -1.233 |
| Evo2 LL [-1.336, -0.726]<sup>N</sup> | -1.296 | -1.333 | -1.419 | -1.345 | -1.169 | -1.240 | -1.269 | -1.238 |
| Dinucleotide JS [0.0000, 0.0988]<sup>N</sup> | 0.0443 | 0.0751 | 0.0333 | 0.0000 | 0.0231 | 0.0209 | 0.0799 | 0.1806 |
| 4-mer JS [0.0000, 0.1437]<sup>N</sup> | 0.1171 | 0.1250 | 0.0976 | 0.0616 | 0.0828 | 0.0799 | 0.1257 | 0.2016 |
| GC content (%) [35, 65]<sup>Q</sup> | 51.01 | 33.34 | 49.47 | 47.11 | 45.14 | 47.48 | 31.04 | 77.86 |
| GC-bound pass (%) | 100.00 | 28.71 | 100.00 | 94.34 | 87.70 | 93.95 | 22.66 | 0.00 |
| CpG O/E [0.049, 0.515]<sup>N</sup> | 0.176 | 0.738 | 0.776 | 0.240 | 0.321 | 0.181 | 0.212 | 1.009 |
| Longest HP (bp) [1, 10]<sup>Q</sup> | 4.66 | 6.77 | 5.10 | 5.89 | 7.29 | 5.52 | 9.93 | 7.76 |
| Max. base fraction [0.25, 0.40]<sup>Q</sup> | 0.278 | 0.378 | 0.274 | 0.313 | 0.316 | 0.311 | 0.406 | 0.437 |

#### Transcription Factor Binding Comparison

Across 1,161 loci spanning 18 TFs, GENECraft reaches **94.56%** macro-averaged full-joint success (96.38% micro-averaged), outperforming the strongest external baseline, Fast SeqProp (17.49%), by over 77 percentage points (Table 3). Optimizing oracle utility without contextual sequence-prior regularization, Fast SeqProp incurs elevated dinucleotide divergence (0.0952) and weak cross-oracle Borzoi transfer (*−*0.016). In contrast, GENECraft pairs scheduled functional updates with bidirectional Carbon guidance to embed canonical motifs within compatible flanking syntax. This halves dinucleotide divergence (0.0495) and supports cross-oracle generalization on held-out Enformer (1.337) and Borzoi (1.060), achieving *≥* 95% joint success on 13 of 18 targets (Appendix J).

**Table 3.**
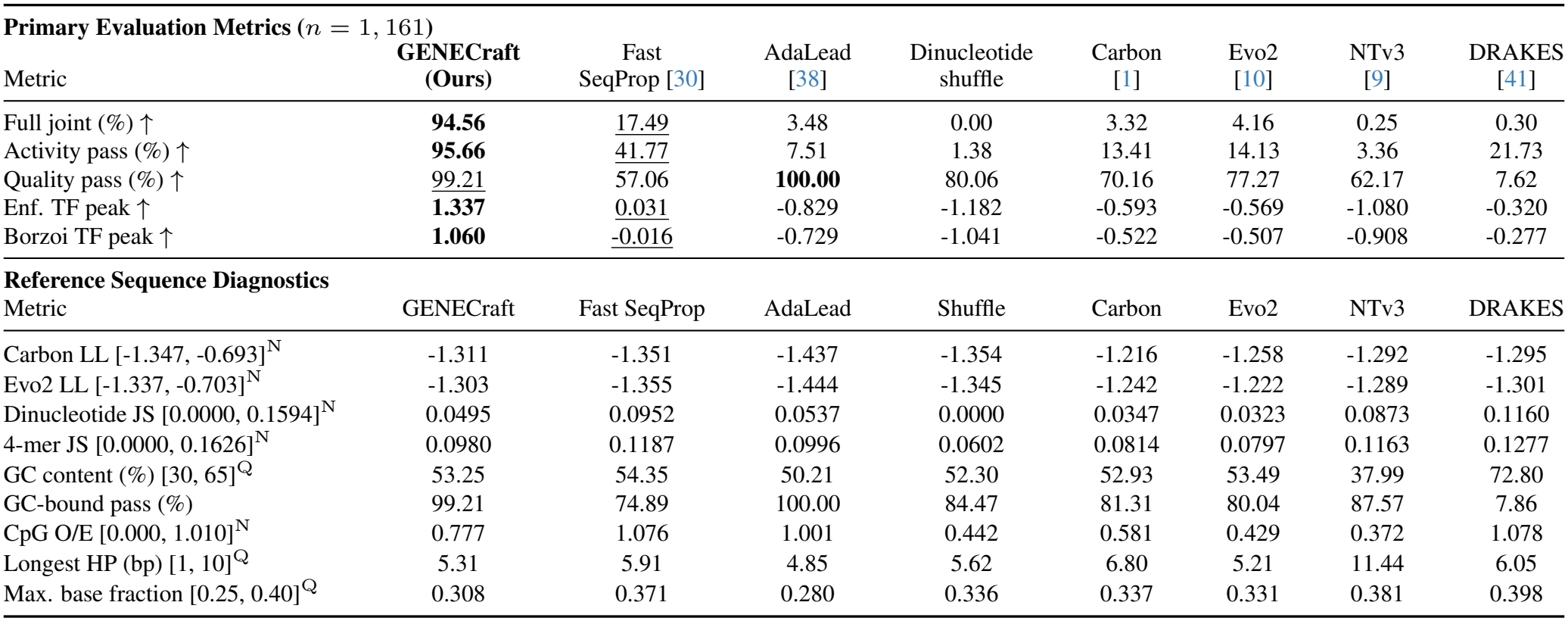
Comparison on the transcription factor binding benchmark (*n* = 1, 161 GM12878 loci). Results report equal-weighted macro-averages across 18 TFs on held-out Enformer and Borzoi oracles. Per-target breakdowns are provided in Supplementary Table 19. Reference interval conventions (N, Q) follow Table 1.

| Metric | GENECraft (Ours) | Fast SeqProp [30] | AdaLead [38] | Dinucleotide shuffle | Carbon [1] | Evo2 [10] | NTv3 [9] | DRAKES [41] |
|---|---|---|---|---|---|---|---|---|
| **Primary Evaluation Metrics** (*n* = 1, 161) |  |  |  |  |  |  |  |  |
| Metric | **GENECraft (Ours)** | Fast SeqProp [30] | AdaLead [38] | Dinucleotide shuffle | Carbon [1] | Evo2 [10] | NTv3 [9] | DRAKES [41] |
| Full joint (%) ↑ | **94.56** | 17.49 | 3.48 | 0.00 | 3.32 | 4.16 | 0.25 | 0.30 |
| Activity pass (%) ↑ | **95.66** | 41.77 | 7.51 | 1.38 | 13.41 | 14.13 | 3.36 | 21.73 |
| Quality pass (%) ↑ | 99.21 | 57.06 | **100.00** | 80.06 | 70.16 | 77.27 | 62.17 | 7.62 |
| Enf. TF peak ↑ | **1.337** | 0.031 | -0.829 | -1.182 | -0.593 | -0.569 | -1.080 | -0.320 |
| Borzoi TF peak ↑ | **1.060** | -0.016 | -0.729 | -1.041 | -0.522 | -0.507 | -0.908 | -0.277 |
| **Reference Sequence Diagnostics** |  |  |  |  |  |  |  |  |
| Metric | GENECraft | Fast SeqProp | AdaLead | Shuffle | Carbon | Evo2 | NTv3 | DRAKES |
| Carbon LL [-1.347, -0.693]<sup>N</sup> | -1.311 | -1.351 | -1.437 | -1.354 | -1.216 | -1.258 | -1.292 | -1.295 |
| Evo2 LL [-1.337, -0.703]<sup>N</sup> | -1.303 | -1.355 | -1.444 | -1.345 | -1.242 | -1.222 | -1.289 | -1.301 |
| Dinucleotide JS [0.0000, 0.1594]<sup>N</sup> | 0.0495 | 0.0952 | 0.0537 | 0.0000 | 0.0347 | 0.0323 | 0.0873 | 0.1160 |
| 4-mer JS [0.0000, 0.1626]<sup>N</sup> | 0.0980 | 0.1187 | 0.0996 | 0.0602 | 0.0814 | 0.0797 | 0.1163 | 0.1277 |
| GC content (%) [30, 65]<sup>Q</sup> | 53.25 | 54.35 | 50.21 | 52.30 | 52.93 | 53.49 | 37.99 | 72.80 |
| GC-bound pass (%) | 99.21 | 74.89 | 100.00 | 84.47 | 81.31 | 80.04 | 87.57 | 7.86 |
| CpG O/E [0.000, 1.010]<sup>N</sup> | 0.777 | 1.076 | 1.001 | 0.442 | 0.581 | 0.429 | 0.372 | 1.078 |
| Longest HP (bp) [1, 10]<sup>Q</sup> | 5.31 | 5.91 | 4.85 | 5.62 | 6.80 | 5.21 | 11.44 | 6.05 |
| Max. base fraction [0.25, 0.40]<sup>Q</sup> | 0.308 | 0.371 | 0.280 | 0.336 | 0.337 | 0.331 | 0.381 | 0.398 |

**Table 4.**
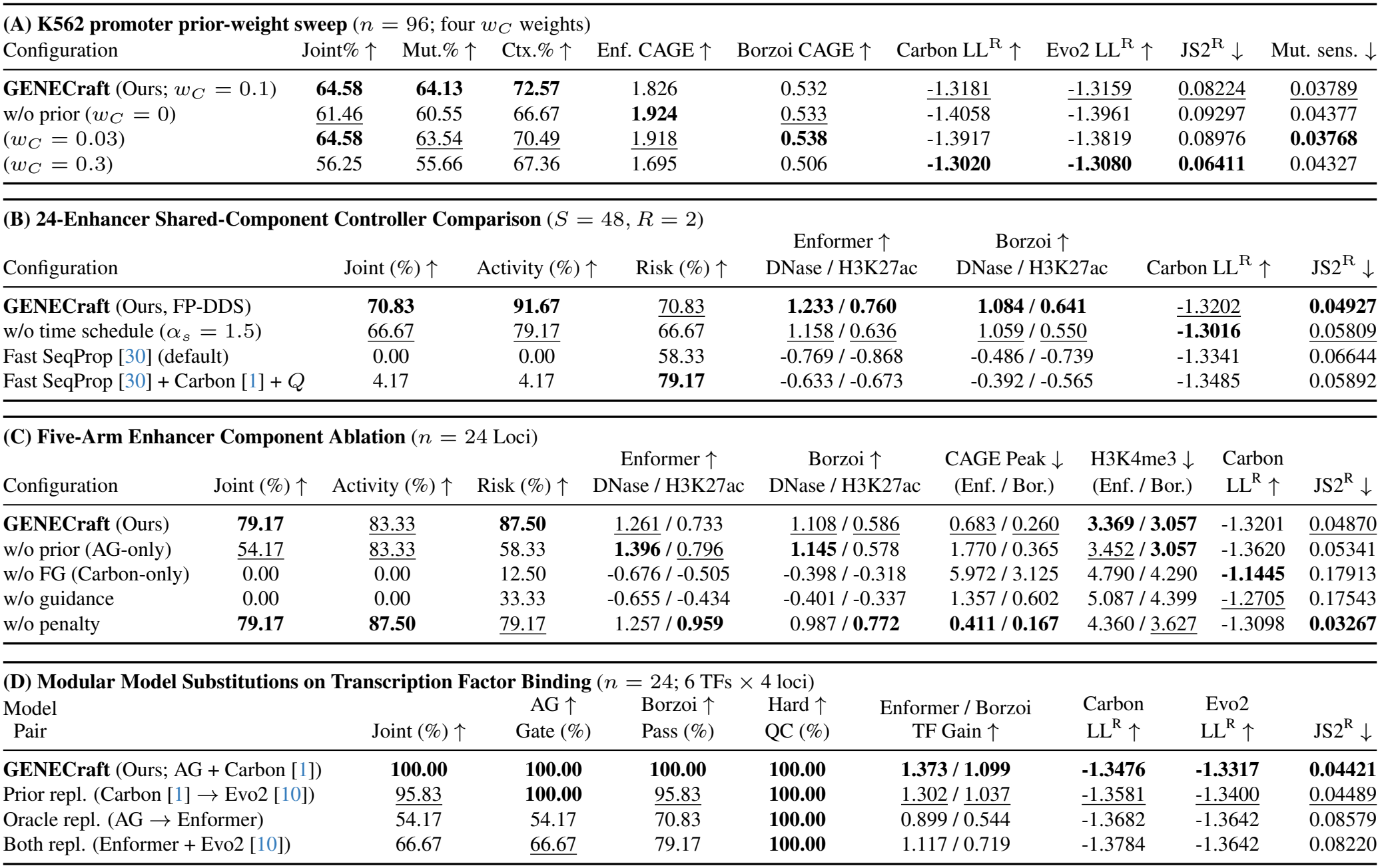
GENECraft ablation, control, and modularity analyses. (A) Carbon weight sweep (*w*_*C*_ ∈ *{*0, 0.03, 0.1, 0.3*}*) across 96 K562 promoters. (B) Controller controls on 24 enhancers (*S* = 48, *R* = 2). (C) Five-arm enhancer ablation (*n* = 24). (D) Modular oracle-prior pairings across 24 TF-binding loci. Mut.% and Ctx.% denote unconditional full-joint success rates across all loci after mutations and context swaps; ^R^: reference diagnostic; LL: log-likelihood (nats/bp); JS_2_: dinucleotide divergence (bits); Sens.: Borzoi CAGE mutation sensitivity. Bold and underline denote the best and second-best distinct displayed values per metric within each panel, including ties, with paired Enformer/Borzoi readouts ranked separately.

| **(A) K562 promoter prior-weight sweep** (*n* = 96; four *w*<sub>*C*</sub> weights) |  |  |  |  |  |  |  |  |  |
| Configuration | Joint% ↑ | Mut.% ↑ | Ctx.% ↑ | Enf. CAGE ↑ | Borzoi CAGE ↑ | Carbon LL<sup>R</sup> ↑ | Evo2 LL<sup>R</sup> ↑ | JS2<sup>R</sup> ↓ | Mut. sens. ↓ |
|---|---|---|---|---|---|---|---|---|---|
| **GENECraft** (Ours; *w*<sub>*C*</sub> = 0.1) | **64.58** | **64.13** | **72.57** | 1.826 | 0.532 | -1.3181 | -1.3159 | 0.08224 | 0.03789 |
| w/o prior (*w*<sub>*C*</sub> = 0) | 61.46 | 60.55 | 66.67 | **1.924** | 0.533 | -1.4058 | -1.3961 | 0.09297 | 0.04377 |
| (*w*<sub>*C*</sub> = 0.03) | **64.58** | 63.54 | 70.49 | 1.918 | **0.538** | -1.3917 | -1.3819 | 0.08976 | **0.03768** |
| (*w*<sub>*C*</sub> = 0.3) | 56.25 | 55.66 | 67.36 | 1.695 | 0.506 | **-1.3020** | **-1.3080** | **0.06411** | 0.04327 |
| **(B) 24-Enhancer Shared-Component Controller Comparison** (*S* = 48, *R* = 2) |  |  |  |  |  |  |  |  |  |
| Configuration | Joint (%) ↑ | Activity (%) ↑ | Risk (%) ↑ | Enformer ↑<br>DNase / H3K27ac | Borzoi ↑<br>DNase / H3K27ac | Carbon LL<sup>R</sup> ↑ | JS2<sup>R</sup> ↓ |  |  |
| **GENECraft** (Ours, FP-DDS) | **70.83** | **91.67** | 70.83 | **1.233 / 0.760** | **1.084 / 0.641** | -1.3202 | **0.04927** |  |  |
| w/o time schedule (*α*<sub>*s*</sub> = 1.5) | 66.67 | 79.17 | 66.67 | 1.158 / 0.636 | 1.059 / 0.550 | **-1.3016** | 0.05809 |  |  |
| Fast SeqProp [30] (default) | 0.00 | 0.00 | 58.33 | -0.769 / -0.868 | -0.486 / -0.739 | -1.3341 | 0.06644 |  |  |
| Fast SeqProp [30] + Carbon [1] + *Q* | 4.17 | 4.17 | **79.17** | -0.633 / -0.673 | -0.392 / -0.565 | -1.3485 | 0.05892 |  |  |
| **(C) Five-Arm Enhancer Component Ablation** (*n* = 24 Loci) |  |  |  |  |  |  |  |  |  |
| Configuration | Joint (%) ↑ | Activity (%) ↑ | Risk (%) ↑ | Enformer ↑<br>DNase / H3K27ac | Borzoi ↑<br>DNase / H3K27ac | CAGE Peak ↓<br>(Enf. / Bor.) | H3K4me3 ↓<br>(Enf. / Bor.) | Carbon<br>LL<sup>R</sup> ↑ | JS2<sup>R</sup> ↓ |
| **GENECraft** (Ours) | **79.17** | 83.33 | **87.50** | 1.261 / 0.733 | 1.108 / 0.586 | 0.683 / 0.260 | **3.369 / 3.057** | -1.3201 | 0.04870 |
| w/o prior (AG-only) | 54.17 | 83.33 | 58.33 | **1.396** / 0.796 | **1.145** / 0.578 | 1.770 / 0.365 | 3.452 / **3.057** | -1.3620 | 0.05341 |
| w/o FG (Carbon-only) | 0.00 | 0.00 | 12.50 | -0.676 / -0.505 | -0.398 / -0.318 | 5.972 / 3.125 | 4.790 / 4.290 | **-1.1445** | 0.17913 |
| w/o guidance | 0.00 | 0.00 | 33.33 | -0.655 / -0.434 | -0.401 / -0.337 | 1.357 / 0.602 | 5.087 / 4.399 | -1.2705 | 0.17543 |
| w/o penalty | **79.17** | **87.50** | 79.17 | 1.257 / **0.959** | 0.987 / **0.772** | **0.411 / 0.167** | 4.360 / 3.627 | -1.3098 | **0.03267** |
| **(D) Modular Model Substitutions on Transcription Factor Binding** (*n* = 24; 6 TFs × 4 loci) |  |  |  |  |  |  |  |  |  |
| Model<br>Pair | Joint (%) ↑ | AG ↑<br>Gate (%) | Borzoi ↑<br>Pass (%) | Hard ↑<br>QC (%) | Enformer / Borzoi<br>TF Gain ↑ | Carbon<br>LL<sup>R</sup> ↑ | Evo2<br>LL<sup>R</sup> ↑ | JS2<sup>R</sup> ↓ |  |
| **GENECraft** (Ours; AG + Carbon [1]) | **100.00** | **100.00** | **100.00** | **100.00** | **1.373 / 1.099** | **-1.3476** | **-1.3317** | **0.04421** |  |
| Prior repl. (Carbon [1] → Evo2 [10]) | 95.83 | **100.00** | 95.83 | **100.00** | 1.302 / 1.037 | -1.3581 | -1.3400 | 0.04489 |  |
| Oracle repl. (AG → Enformer) | 54.17 | 54.17 | 70.83 | **100.00** | 0.899 / 0.544 | -1.3682 | -1.3642 | 0.08579 |  |
| Both repl. (Enformer + Evo2 [10]) | 66.67 | 66.67 | 79.17 | **100.00** | 1.117 / 0.719 | -1.3784 | -1.3642 | 0.08220 |  |

### 4.3. Ablation Studies and Analysis

#### Impact of In-Loop Carbon Prior

The 96-locus K562 promoter sweep (Table 4A; Supplementary Tables 10B–C, 14, and 15) shows how in-loop Carbon regularization coordinates functional compliance, native similarity, and robustness. Relative to optimization without Carbon (*w*_*C*_ = 0), *w*_*C*_ = 0.1 raises full-joint success (64.58% vs. 61.46%) and maximizes joint success under mutation (64.13% vs. 60.55%) and context swaps (72.57% vs. 66.67%), while improving Carbon and Evo2 log-likelihoods, dinucleotide divergence JS_2_ (0.08224 vs. 0.09297, *−*11.5%), and descriptive Borzoi CAGE mutation sensitivity (*−*13.4%). Lighter weighting (*w*_*C*_ = 0.03) ties nominal yield (64.58%) but weakens perturbation joint success and prior alignment, whereas *w*_*C*_ = 0.3 over-regularizes yield (56.25%). Balancing functional activity with sequence prior knowledge, we therefore select *w*_*C*_ = 0.1 as the default Carbon guidance weight.

#### Shared-Component Controller Comparison

To isolate the algorithmic contribution of the FP-DDS controller, we match Fast SeqProp to our exact *U*_AG_ objective, Carbon prior, *Q* penalty, feasibility gate, readout, and compute budget (∼ 46.2 s; Table 4B, Appendix L). Despite shared guidance objectives and readout, component-matched Fast SeqProp (FSP + Carbon + *Q*) achieves only 1*/*24 joint compliance with negative external activity gains, whereas FP-DDS reaches **17/24** (*p* = 9.16 *×* 10^*−*5^). Moreover, our linear step-size ramp outperforms a constant step schedule (*α*_*s*_ = 1.5), improving external activity and native dinucleotide similarity (JS_2_: 0.0493 vs. 0.0581).

#### Enhancer Component Ablation

Dissecting individual guidance forces on the 24-locus enhancer cohort confirms their complementary roles (Table 4C, Figure 3). Removing functional gradients (*w/o FG*) collapses joint success from 79.2% (19/24) to 0.0%, confirming that active functional steering is indispensable. While removing Carbon (*w/o prior*) maintains raw activity pass (20/24), it degrades chromatin-risk compliance from 87.5% (21/24) to 58.3% (14/24), lowering joint success to 54.2%. Specifically, Carbon co-guidance suppresses off-target Enformer CAGE peaks from 1.770 to 0.683 (*p* = 0.0350; Figure 3a,b). Similarly, omitting the soft penalty *Q*(*z*) (*w/o penalty*) inflates undesirable H3K4me3 promoter marks across both external oracles (Figure 3c,d) and increases risk failures.

**Figure 3.**
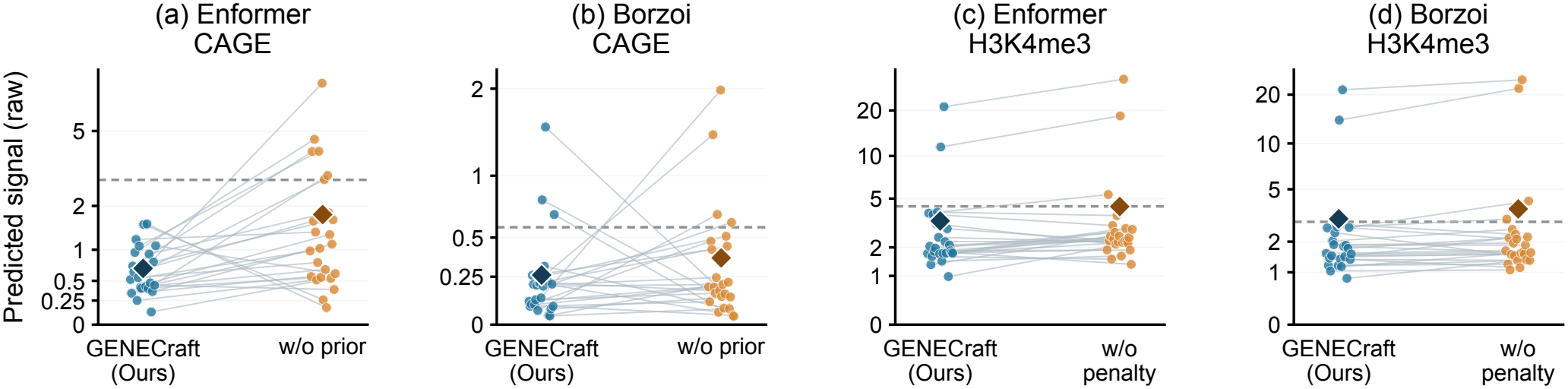
Paired enhancer chromatin-risk ablations across 24 matched loci. **(a**,**b)** Predicted Enformer and Borzoi CAGE peaks comparing full GENECraft against prior removal (*w/o prior, w*_*C*_ = 0). **(c**,**d)** Predicted mean H3K4me3 signals comparing full GENECraft against penalty removal (*w/o penalty, w*_*Q*_ = 0). Dashed lines denote risk ceilings; diamonds mark cohort means.

#### Modularity and Model Substitution Comparison

Finally, we verify that FP-DDS serves as a plug-and-play interface across distinct foundation models by crossing two functional oracles (AlphaGenome, Enformer) with two gLM priors (Carbon, Evo2) on 24 TF loci (Table 4D). All four pairings achieve 100% (24/24) hard-constraint compliance without hyperparameter retuning. While our default AlphaGenome+Carbon pairing delivers the best overall performance (24/24 joint success; 1.099 held-out Borzoi gain), substituting Evo2 as the prior performs comparably (23/24 joint; 1.037 Borzoi gain), and coupling Enformer with Evo2 or Carbon also yields high transferability. This confirms that FP-DDS provides a generalizable, model-agnostic framework for controllable regulatory DNA synthesis.

### 4.4. Case Study

#### Coordinated Latent-Logit Trajectory

To illustrate how GENECraft coordinates complementary guidance forces, Fig. 4a tracks a 445-bp GM12878 *BHLHE40* locus across 24 updates. In the projected PCA plane, AlphaGenome functional gradients and Carbon priors exhibit early cooperation (update 4), mid-trajectory divergence where Carbon restrains prior-deviating oracle ascent (update 15), and later realignment (update 18; exact cosines in Appendix P). This coupled trajectory elevates predicted AlphaGenome utility from 0.75 to 131.0 while improving exact Carbon log-likelihood from *−*1.436 to *−*1.310 nats/bp, achieving initial constraint feasibility at update 5 and selecting the highest-utility feasible design at update 24.

**Figure 4.**
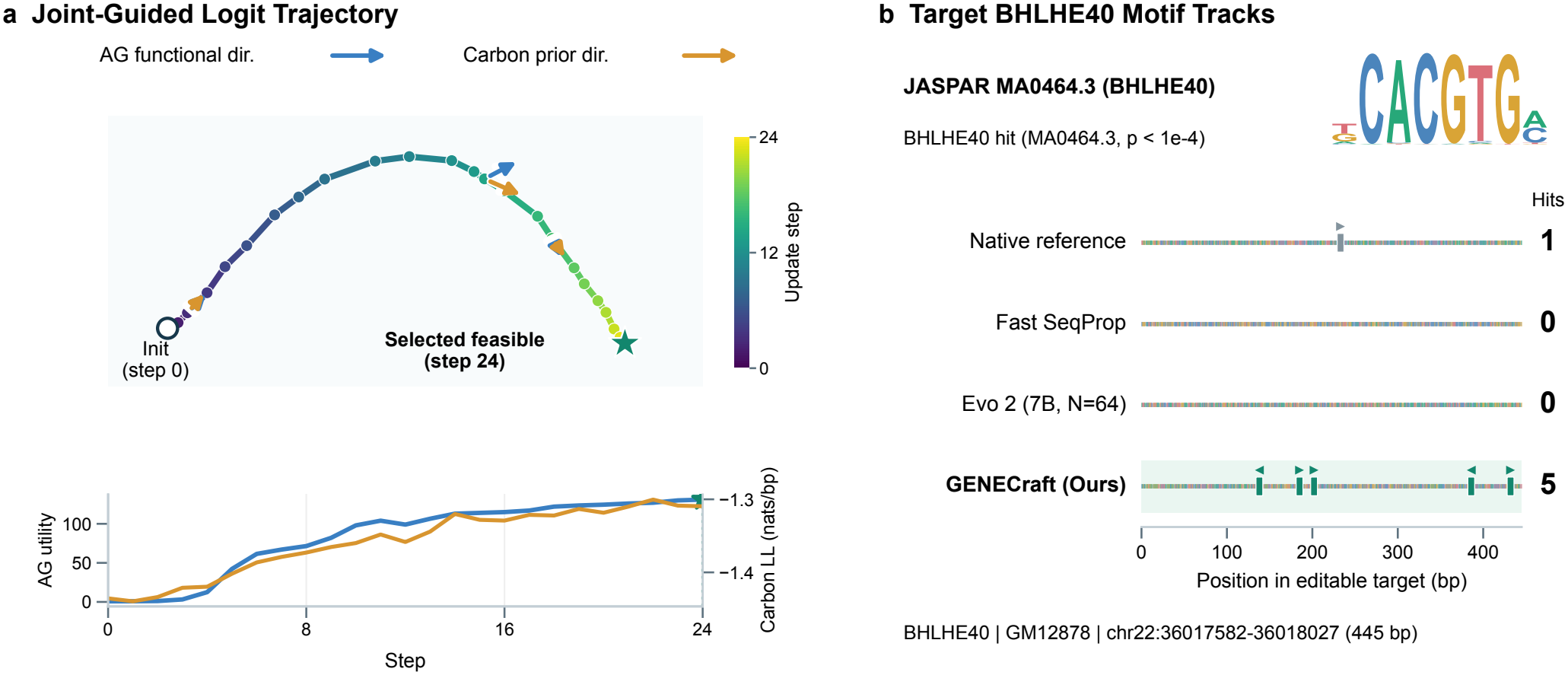
Joint-guided optimization dynamics and target-motif acquisition at a *BHLHE40* locus. **(a)** Latent-logit PCA trajectory (0–24), with unit-length arrows showing projected AlphaGenome and Carbon guidance directions (updates 4, 15, 18), a star marking the selected feasible design, and inset utility and likelihood traces. **(b)** Positional JASPAR *BHLHE40* motif tracks (*p <* 10^*−*4^) for native, GENECraft, Fast SeqProp, and Evo 2 sequences (Appendix P).

#### Targeted Canonical Motif Acquisition

Iterative joint functional-prior updates translate this trajectory into targeted motif recruitment supporting high predicted activity (Fig. 4b). Post hoc JASPAR scanning of the 445-bp locus for the canonical *BHLHE40* motif (MA0464.3, *p <* 10^*−*4^) identifies 5 matches in GENECraft versus 1 in the native sequence and 0 in Fast SeqProp and AlphaGenome-ranked best-of-64 Evo 2. Across the 17-TF, 1,139-locus grammar audit (Appendix R; Supplementary Table 30), GENECraft similarly elevates canonical motif presence to 64.13% versus 57.96% for AlphaGenome-only designs and 38.40% natively while maintaining 100% cross-locus uniqueness, coupling motif acquisition with sequence-level regularity.

## 5. Conclusion

GENECraft introduces a training-free Function- and Prior-Constrained Discrete Dynamical System (FP-DDS) that integrates genomic language models, sequence-to-function oracles, and explicit compositional constraints along discrete-anchored latent-logit trajectories. Across 2,185 human regulatory loci, GENECraft achieves 74.16% task-macro full-joint success, outperforming seven optimization, evolutionary, and generative baselines under Enformer and Borzoi cross-oracle validation. Objective-matched controls further establish the advantage of the FP-DDS controller (17*/*24 vs. 1*/*24 enhancer success), while Carbon and quality regularization strengthen compositional similarity, off-target risk control, and selected stability endpoints.

## LLM Usage Statement

During preparation of this manuscript and revision, Gemini assisted with manuscript writing, editing, and drafting figure captions and table descriptions; Gemini, Qwen, and DeepSeek provided pre-submission feedback. OpenAI Codex assisted with LATEX editing, document layout, statistical analysis, programmatic plotting from experimental archives, implementation of supplementary ablation and control scripts, and numerical verification, while Qoder previously assisted with LATEX formatting.

## Supplementary Material

### A. Discrete Dynamics and Schedule Details

Each restart initializes DNA uniformly via a coordinate-derived seed and fixed Gumbel noise, scoring the seed sequence before noisy decoding. Functional and prior gradients are evaluated at the decoded one-hot sequence and propagated through Eq. 1 at *τ* = 1. Carbon’s forward and reverse-complement gradients are coordinate-aligned before averaging, while task adapters handle negative-strand promoter orientation.

For unperturbed nucleotide probabilities *q* = softmax(*z*) (excluding fixed Gumbel noise), expected GC fraction 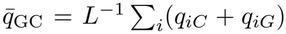, and target length *L >* 1, the differentiable sequence-quality objective is

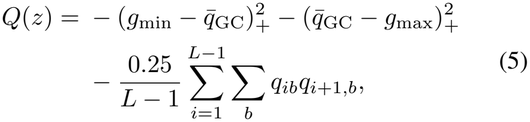

where (*a*)_+_ = max(*a*, 0) and [*g*_min_, *g*_max_] are task-specific GC bounds; these terms penalize global GC violations and adjacent-nucleotide homopolymers. Each gradient component, including *∇*_*z*_*Q*(*z*), is RMS-normalized across the *L×*4 tensor, while the hard gate *I*_QC_ enforces exact single-base fraction and homopolymer run-length limits.

Within each restart, step size *α*_*s*_ = 3*s/*(*S* + 1) grows linearly over *s* ∈ *{*1, …, *S}*, scaling the functional oracle (*α*_*s*_), generative prior (*w*_*C*_*α*_*s*_, default *w*_*C*_ = 0.1), and quality penalty (*w*_*Q*_*α*_*s*_, *w*_*Q*_ = 0.15) so *α*_*s*_ sets overall scale and *w*_*C*_ sets relative prior weight. Post-scaling clipping clip(*·, −*2, 2) bounds updates to |*z*_*s*+1,*ib*_ *− z*_*s,ib*_| *≤* 2 for late-trajectory stability. Fixed Gumbel noise is constant across steps, and matched ablation arms share schedules, initializations, and restart budgets.

The step-weight schedule *α*_*s*_ scales from 3*/*25 = 0.12 to 72*/*25 = 2.88 for *S* = 24 and from 3*/*49 *≈* 0.0612 to 144*/*49 *≈* 2.9388 for *S* = 48. Because GENECraft executes a direct discrete iteration schedule, increasing *S* simultaneously expands the optimization horizon, total compute budget, and cumulative discrete trajectory updates.

Candidate selection pools all decoded trajectories across restarts, prioritizing generation-feasible sequences (*F* = 1) and breaking ties by exact AlphaGenome utility *U*_AG_ (then encounter order); if none is feasible, the highest-utility sequence and its failure flag are retained. Generation feasibility uses only AlphaGenome and composition checks, reserving Enformer and Borzoi for post-generation validation.

### A.1. Deterministic Differentiable 6-mer Token Bridge

To reconcile AlphaGenome’s one-hot representation of *x* ∈ *{*A, C, G, T*}*^*L*^ with Carbon’s causal vocabulary *V*_full_ (containing *V*_DNA_ ≅ *{*A, C, G, T*}*^6^ of |*V*_DNA_| = 4^6^ = 4096 non-overlapping 6-mers) without adapter training, we construct a deterministic differentiable relaxation in shared nucleotide logit space directly from Carbon’s frozen embeddings *E*_*v*_ ∈ ℝ^*d*^ (*v* ∈ *V*_DNA_) and output projection.

#### Algorithm 1

GENECraft: Function-Constrained Discrete Sequence Dynamics

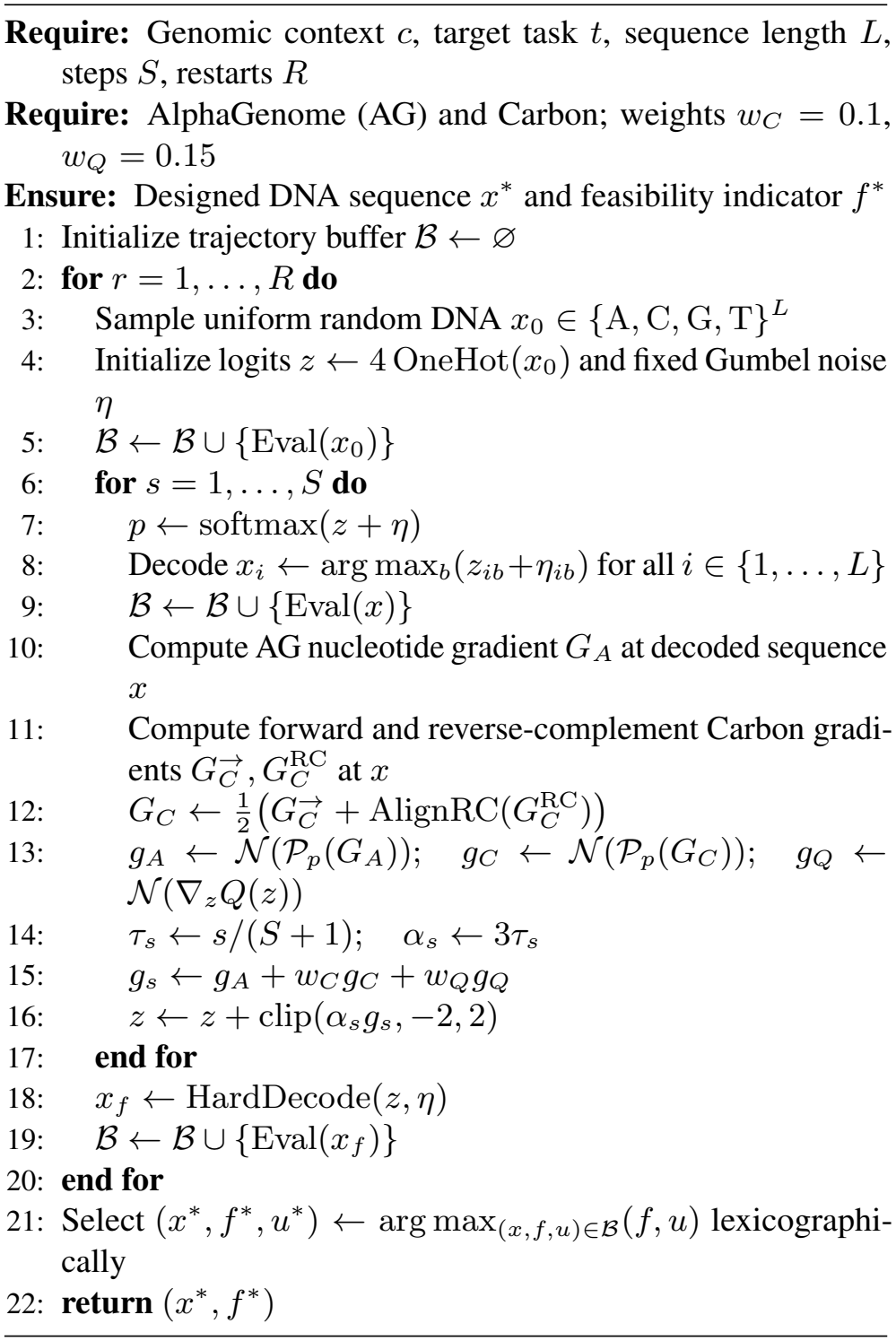

Each row *P*_*i*_ = (*P*_*i*,A_, *P*_*i*,C_, *P*_*i*,G_, *P*_*i*,T_) ∈ Δ^3^ lies on the nucleotide probability simplex, satisfying ∑_*b*∈*{*A,C,G,T*}*_ *P*_*i,b*_ = 1.

#### Exact 6-mer Factorization and Expected Embeddings

Flanks *c*_L_ and *c*_R_ have lengths divisible by 6 to align token boundaries in both orientations. Target length *L* decomposes as *L* = 6*n* + *r* into *n* = ⌊*L/*6⌋ full 6-mer blocks and a trailing *r* = *L* mod 6 ∈ *{*0, …, 5*}*-bp block. From the relaxed probability matrix *P* ∈ [0, 1]^*L×*4^ over (A, C, G, T), each full 6-mer block *t* ∈ *{*0, …, *n−*1*}* induces a factorized distribution *π*_*t*_ ∈ Δ^4096^ over 6-mers *v* = (*v*_0_, …, *v*_5_) ∈ *V*_DNA_:

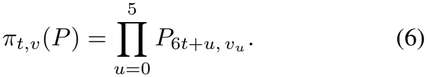

Conditioned on the *C* discrete prefix tokens of *c*_L_, we replace the *n* target token positions in the input embedding sequence with their expected embeddings over the fixed DNA embedding rows *E*_*v*_ under *π*_*t*_(*P* ):

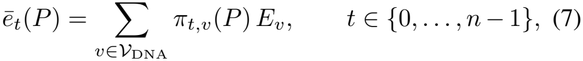

keeping context embeddings fixed and detached.

### Full-Vocabulary Causal Normalization and DNA-Subset Scoring

A causal forward pass over the concatenated context and expected target embeddings yields pre-softmax logits *z*_*t,u*_(*P* ) across *u* ∈ *V*_full_ at the prediction rows for the *n* full target blocks (plus row (*n* + 1) when *r >* 0). Let

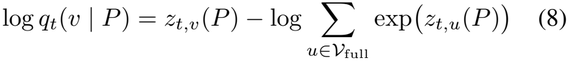

be the exact full-vocabulary log-softmax probability of DNA 6-mer *v* ∈ *V*_DNA_ at step *t* (conditioned on *c*_L_ and blocks *< t*). The surrogate scores the expected log-probability over *V*_DNA_:

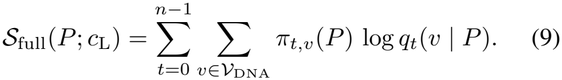

When *r >* 0, the trailing *r* nucleotides form a partial 6-mer scored at row *n* by summing full-vocabulary probabilities *q*_*n*_(*v* | *P* ) over all DNA 6-mers sharing that *r*-base prefix:

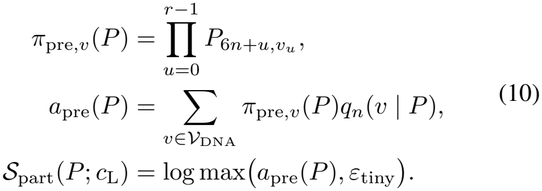

with *ε*_tiny_ ensuring numerical stability. Normalizing by sequence length gives the per-base forward surrogate score:

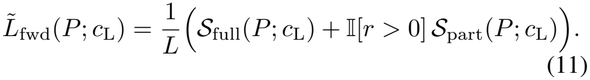

#### Bidirectional Coordinate Alignment

Carbon scores both orientations for strand symmetry. The reverse-complement operator *R* reverses positions (*i* ↦ *L −* 1 *− i*) and permutes bases ((A, C, G, T) ↦ (T, G, C, A)), yielding (*RP* )_*i,b*_ = *P*_*L−*1*−i*, 3*−b*_. Conditioning on rc(*c*_R_) as left context for *RP* defines 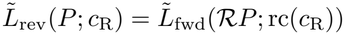, and pulling back its gradient through *R* wrt *P* gives the bidirectional objective and gradient:

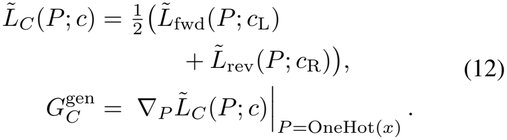

For negative-strand (*−*) promoters oriented 5^*′*^ → 3^*′*^ relative to the transcription start site (TSS), the coordinate adapter maps candidates to genomic coordinates before evaluation and pulls gradients back via 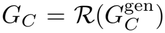 for positive-strand promoters, enhancers, and TF-binding loci, 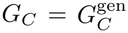.

#### One-Hot Vertex Consistency and Differentiable Surrogate Gradients

Eqs. (6)– 12) are exact at discrete vertices: for *P* = OneHot(*x*) (*x* ∈ *{*A, C, G, T*}*^*L*^), *π*_*t,v*_(*P* ) reduces to 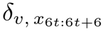 on the realized 6-mer at block *t, ē*_*t*_(*P* ) equals 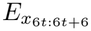, and *π*_pre,*v*_(*P* ) indicates the 4^6*−r*^ tokens matching prefix *x*_6*n*:*L*_. Thus,

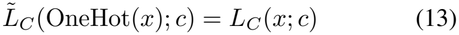

Because *π*_*t,v*_(*P* ) is multilinear in *P* before Carbon’s non-linear transformer layers, Eq. (13) guarantees exact vertex consistency for 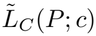. Across 165 comparisons from 55 designs, unbatched `bf16` forward evaluations matched batched hard-token scoring within 9.17 *×* 10^*−*8^ nats/bp median and 0.003504 nats/bp maximum discrepancy (0.2829% max relative difference). Thus, *∇*_*P*_ 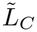 supplies continuous proposer gradients, while logging, feasibility, and selection evaluate exact discrete sequences *x*.

### A.2. Exact Task Utility and Feasibility Definitions

For editable sequence *x* ∈ *{*A, C, G, T*}*^*L*^ at [*s*_t_, *e*_t_) within context window *W* (*W* = 65,536 bp; *L* = 500 bp for enhancers), AlphaGenome predicts 1D tracks at resolution Δ_res_. Restricting track *y* ∈ ℝ^*B*^ to evaluation slice *I* = [*s, e*) (length *N* = *e − s*), the regional and peak-window (width *w*_peak_) means are:

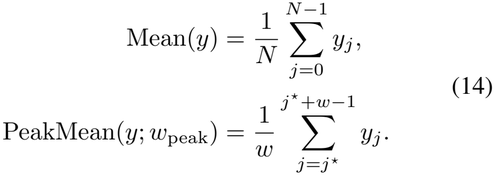

with *w* = min(max(1, *w*_peak_), *N* ), *j*_max_ = arg max_*j*_ *y*_*j*_, and window start *j*^⋆^ = min(max(0, *j*_max_ *−*⌊*w/*2⌋), *N −w*) centered at the track argmax and clamped to [0, *N − w*].

#### Promoter Utility and Generation Feasibility

On oriented CAGE/PRO-cap track *y* over slice *I* (*N* bins), `promoter_upstream` and `tss_window_bp` define TSS bin *b*_tss_ = min(*N −* 1, max(0, round(`promoter_upstream` */*Δ_res_))), width *w*_tss_ = max(1, ⌈`tss_window_bp` */*Δ_res_⌉), and clamped window *I*_tss_ = [lo, hi) (lo = max(0, min(*b*_tss_ *−* ⌊*w*_tss_*/*2⌋, *N − w*_tss_)), hi = min(*N*, lo + *w*_tss_)).

With focal mean 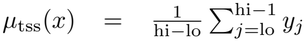, flanking mean 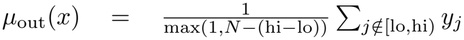, *λ*_off_ = `offtarget_weight`, and *ε* = 10^*−*6^:

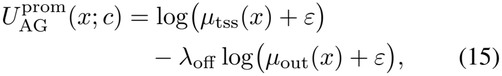

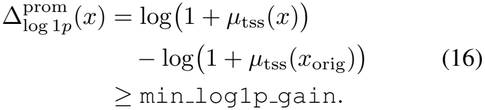

where Eq. (16) defines generation-stage functional feasibility relative to native *x*_orig_.

#### TF-Binding Utility, Generator Feasibility, and Common Benchmark Screen

For TF-binding design (`guidance_mode` = `maximize`, `activity_transform` = `raw` ), AlphaGenome scores ChIP track *y* over *I* via 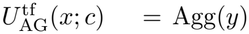. Generator feasibility requires strict native improvement (PeakMean(*y*(*x*); *w*_peak_) *>* PeakMean(*y*(*x*_orig_); *w*_peak_)), whereas benchmark success requires PeakMean(*y*(*x*); *w*_peak_) to reach the condition-specific native *p*_50_ threshold with gain 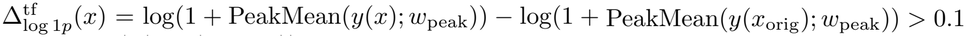.

#### Enhancer Composite Utility and Generation Feasibility

A single AlphaGenome pass evaluates three heads over target slice *I* (500 bp) and risk slice *I*_risk_ (`h3k4me3_window_bp` = 2048 bp): (1) `DNASE` (`CL:2000045`, tr. 0, *w*_D_ = 1.0); (2) `CHIP_HISTONE` (`CL:2000045`, tr. 0, *w*_H_ = 2.0) for H3K27ac (ch. 0), H3K4me3 (ch. 1), and H3K4me1 (ch. 2); and (3) penalty-only `CAGE` (`CL:0002566`, `CL:0002567`, tr. 0, *w*_CAGE_ = 1.0). With signals *A*_D_(*x*) = Mean(*y*_DNASE_[*I*]), *A*_H27_(*x*) = Mean(*y*_H3K27ac_[*I*]), *M*_H4me1_(*x*) = Mean(*y*_H3K4me1_[*I*_risk_]), *R*_H4me3_(*x*) = Mean(*y*_H3K4me3_[*I*_risk_]), *R*_CAGE_(*x*) = PeakMean(*y*_CAGE_[*I*]; *w*_peak_) and parameters *λ*_H4me3_ = 4.0, *τ*_H4me3_ = 0.7974365234375002, *λ*_H4me1_ = 0.5, *λ*_CAGE_ = 10.0, *τ*_CAGE_ = 0.0016947865486145054:

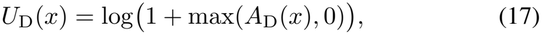

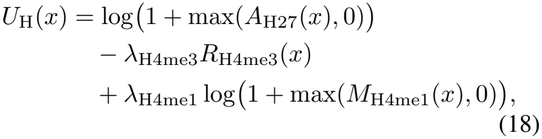

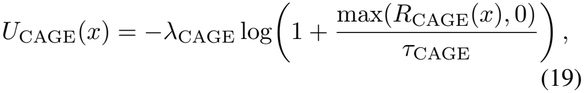

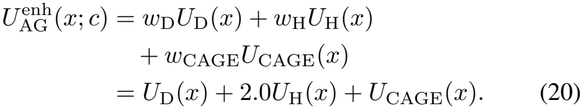

Enhancer generation feasibility requires: (i) for *m* ∈ *{*DNASE, H3K27AC*}* and *s* ∈ *{*mean, peak mean*}*, log(1 + *m*_*s*_(*x*)) *−* log(1 + *m*_*s*_(*x*_orig_)) *≥ γ*_*m*_ (*γ*_DNASE_ = *γ*_H3K27AC_ = 0.1) and *m*_mean_(*x*) *> m*_mean_(*x*_orig_); and (ii) risk caps *R*_H4me3_(*x*) *≤ τ*_H4me3_ and *R*_CAGE_(*x*) *≤ τ*_CAGE_.

#### Quality Control, Feasibility, and Evaluation Interfaces

Sequence quality and oracle evaluations operate at three interfaces:

1. **Soft Compositional Penalty** *Q*(*z*): During continuous updates, *Q*(*z*) acts on softmax(*z*) with weight *w*_*Q*_ = 0.15 using task-specific GC bounds [*g*_min_, *g*_max_] ([0.35, 0.65] for promoters and enhancers; [0.30, 0.65] for TF binding) to discourage GC skew and adjacent-base runs, without entering discrete ranking or feasibility.
2. **Hard Compositional Gate** *I*_QC_(*x*) **and Changed-Sequence Feasibility:** On discrete *x, I*_QC_(*x*) = 1 iff

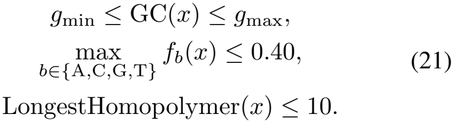

where [*g*_min_, *g*_max_] = [0.35, 0.65] for promoters and enhancers and [0.30, 0.65] for TF-binding elements. Total generator feasibility *F* (*x*; *c, t*) = 1 requires *I*_QC_(*x*) = 1, the separate non-triviality condition *x* ≠ *x*_orig_ (`require_changed` = `true` ), and the task-specific functional constraints above.
3. **In-Loop vs. External Oracles:** Default GENECraft (AlphaGenome + Carbon) generation and candidate selection rely solely on *F* (*x*; *c, t*) and *U*_AG_(*x*; *c, t*), holding out Enformer, Borzoi, and Evo2 strictly for post-generation validation; by contrast, the dedicated predictor/prior model-swap benchmark arms intentionally substitute Enformer or Evo2 inside the optimization loop.

### B. Natural-Sequence Initialization

To isolate initialization effects beyond *de novo* design, we compare random and native starts on 84 matched loci (24 promoters, 24 enhancers, 36 TF loci; two per TF), initializing *z*_0_ = 4 OneHot(*x*_0_) from *x*_0_ under identical per-locus Gumbel noise (distinct across enhancer restarts), dense AG– Carbon weights, schedules, step budgets, selection rules, and five-model evaluation thresholds.

Native initialization increases Carbon and external Evo2 log-likelihoods across all tasks while reducing dinucleotide divergence by 56.7%, 22.7%, and 42.5% for promoters, enhancers, and TF binding, respectively (Table 5). Independent functional gains and activity-pass rates also improve overall, while task-macro full-joint success remains comparable between random and natural starts (65.74% vs. 65.28%).

**Table 5.**
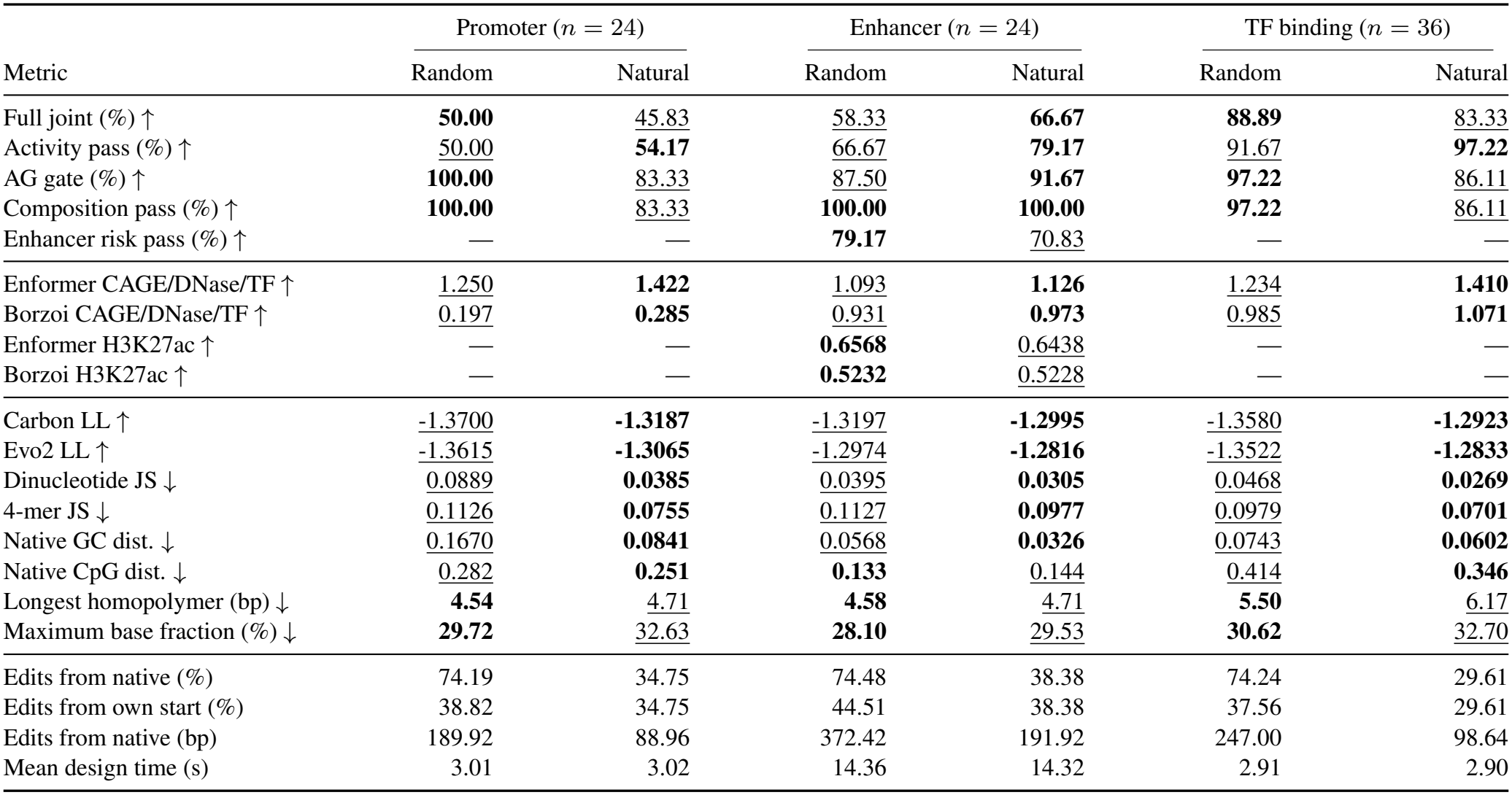
Matched initialization ablation. Random versus native initialization. Metrics: native-relative Δ log(1 + *s*) (activity pass: external gains *≥*0.1), bidirectional log-likelihood (LL, nats/bp), dinucleotide JS divergence (bits), native GC and CpG *O/E* distances, edit fractions, and warm H100 runtimes. TF values macro-average 18 factors; Bold and underline denote the best and second-best distinct displayed values per metric within each comparable panel, including ties; unranked reference diagnostics are excluded.

| Metric | Promoter (*n* = 24) |  | Enhancer (*n* = 24) |  | TF binding (*n* = 36) |  |
|---|---|---|---|---|---|---|
|  | Random | Natural | Random | Natural | Random | Natural |
| Full joint (%) ↑ | **50.00** | 45.83 | 58.33 | **66.67** | **88.89** | 83.33 |
| Activity pass (%) ↑ | 50.00 | **54.17** | 66.67 | **79.17** | 91.67 | **97.22** |
| AG gate (%) ↑ | **100.00** | 83.33 | 87.50 | **91.67** | **97.22** | 86.11 |
| Composition pass (%) ↑ | **100.00** | 83.33 | **100.00** | **100.00** | **97.22** | 86.11 |
| Enhancer risk pass (%) ↑ | — | — | **79.17** | 70.83 | — | — |
| Enformer CAGE/DNase/TF ↑ | 1.250 | **1.422** | 1.093 | **1.126** | 1.234 | **1.410** |
| Borzoi CAGE/DNase/TF ↑ | 0.197 | **0.285** | 0.931 | **0.973** | 0.985 | **1.071** |
| Enformer H3K27ac ↑ | — | — | **0.6568** | 0.6438 | — | — |
| Borzoi H3K27ac ↑ | — | — | **0.5232** | 0.5228 | — | — |
| Carbon LL ↑ | -1.3700 | **-1.3187** | -1.3197 | **-1.2995** | -1.3580 | **-1.2923** |
| Evo2 LL ↑ | -1.3615 | **-1.3065** | -1.2974 | **-1.2816** | -1.3522 | **-1.2833** |
| Dinucleotide JS ↓ | 0.0889 | **0.0385** | 0.0395 | **0.0305** | 0.0468 | **0.0269** |
| 4-mer JS ↓ | 0.1126 | **0.0755** | 0.1127 | **0.0977** | 0.0979 | **0.0701** |
| Native GC dist. ↓ | 0.1670 | **0.0841** | 0.0568 | **0.0326** | 0.0743 | **0.0602** |
| Native CpG dist. ↓ | 0.282 | **0.251** | **0.133** | 0.144 | 0.414 | **0.346** |
| Longest homopolymer (bp) ↓ | **4.54** | 4.71 | **4.58** | 4.71 | **5.50** | 6.17 |
| Maximum base fraction (%) ↓ | **29.72** | 32.63 | **28.10** | 29.53 | **30.62** | 32.70 |
| Edits from native (%) | 74.19 | 34.75 | 74.48 | 38.38 | 74.24 | 29.61 |
| Edits from own start (%) | 38.82 | 34.75 | 44.51 | 38.38 | 37.56 | 29.61 |
| Edits from native (bp) | 189.92 | 88.96 | 372.42 | 191.92 | 247.00 | 98.64 |
| Mean design time (s) | 3.01 | 3.02 | 14.36 | 14.32 | 2.91 | 2.90 |

**Table 6.** Modular enhancer backbone substitutions. Joint-pass rates and warm H200 runtimes across *n* = 24 enhancer loci (48 steps, 2 restarts) when pairing functional oracles (AlphaGenome, Enformer) with sequence priors (Carbon, Evo2) under held-out Borzoi evaluation.

| **Modular Backbone Substitutions on Enhancer Activation (*n* = 24)** |  |  |  |  |  |  |  |  |  |
|---|---|---|---|---|---|---|---|---|---|
| Model combination | Joint% ↑ | Borzoi% ↑ | Quality% ↑ | Risk% ↑ | Min. gain ↑ | Carbon LL<sup>R</sup> ↑ | Evo2 LL<sup>R</sup> ↑ | JS2<sup>R</sup> ↓ | Time (s) ↓ |
| GENECraft (Ours; AG + Carbon) | **79.17** | **87.50** | **100.00** | **87.50** | 0.490 | **-1.3201** | -1.2984 | 0.04870 | **13.95** |
| Prior repl. (Carbon → Evo2) | 70.83 | **87.50** | **100.00** | 75.00 | 0.498 | -1.3260 | **-1.2921** | **0.04704** | 168.77 |
| Oracle repl. (AG → Enformer) | 4.17 | 75.00 | **100.00** | 66.67 | **0.556** | -1.3293 | -1.3154 | 0.08226 | 58.50 |
| Both repl. (Enformer + Evo2) | 12.50 | 75.00 | **100.00** | 79.17 | 0.495 | -1.3374 | -1.3050 | 0.08117 | 214.39 |

**Table 7.** Empirical enhancer risk-gate thresholds. Inclusive native quantiles (*P*_99_ CAGE peak ceiling, *P*_80_ H3K4me3 mean ceiling, *P*_10_ H3K4me1*/*(H3K4me3 + 10^*−*6^) floor) calibrated independently on raw Enformer and Borzoi predictions across *n* = 50 reference sequences from `Genomic_MEL_regions.fa` .

| Constraint | Enformer | Borzoi |
|---|---|---|
| CAGE peak ≤ | 2.818158 | 0.573727 |
| H3K4me3 mean ≤ | 4.374960 | 2.899119 |
| H3K4me1/H3K4me3 ≥ | 1.443216 | 1.068096 |

**Table 8.**
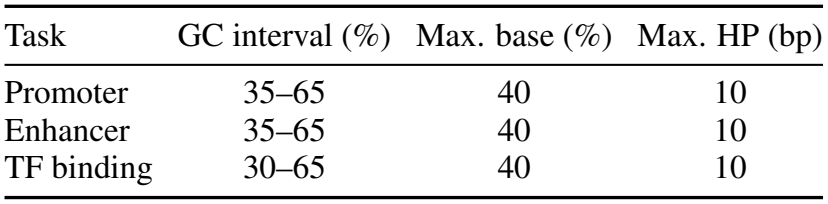
Hard sequence-quality thresholds. Inclusive simultaneous bounds for *I*_QC_ on the editable target of length *L* (excluding flanks); HP denotes the longest contiguous homopolymer run.

**Table 9.** Full-cohort internal architecture comparison. Evaluation across 2,185 loci (Tables 1–3; failed designs retained; equal-weight macro-averages across 18 TFs and tasks). Continuous joint guidance (GENECraft) is compared against sequential AG–Carbon refinement using functional readouts, bidirectional log-likelihood (LL, nats/bp), and native-relative dinucleotide divergence (JS2, bits).

| Task | System | Joint% ↑ | QC% ↑ | Enf. target(s) ↑ | Borzoi target(s) ↑ | Carbon LL ↑ | Evo2 LL ↑ | JS2 ↓ |
|---|---|---|---|---|---|---|---|---|
| Promoter | AG-only | 61.72 | 99.80 | **1.977** | 0.509 | -1.4088 | -1.3976 | 0.09668 |
| Promoter | Sequential AG–Carbon | **63.28** | 99.80 | 1.969 | **0.524** | -1.3950 | -1.3854 | 0.09157 |
| Promoter | GENECraft (Ours) | 62.70 | **100.00** | 1.934 | 0.515 | **-1.319** | **-1.312** | **0.08399** |
| Enhancer | AG-only | 56.05 | 99.22 | 0.975/0.547 | 0.804/0.419 | -1.3658 | -1.3507 | 0.05293 |
| Enhancer | Sequential AG–Carbon | 56.64 | 99.22 | 0.996/0.562 | 0.827/0.429 | -1.3593 | -1.3434 | 0.05158 |
| Enhancer | GENECraft (Ours) | **65.23** | **100.00** | **1.157/0.700** | **1.033/0.593** | **-1.3188** | **-1.2957** | **0.04433** |
| TF binding | AG-only | 88.89 | **100.00** | 1.149 | 0.891 | -1.3880 | -1.3800 | 0.07018 |
| TF binding | Sequential AG–Carbon | 89.13 | **100.00** | 1.152 | 0.894 | -1.3796 | -1.3716 | 0.06779 |
| TF binding | GENECraft (Ours) | **94.56** | 99.21 | **1.337** | **1.060** | **-1.311** | **-1.303** | **0.04948** |
| Equal-task macro | AG-only | 68.89 | – |  |  |  |  |  |
| Equal-task macro | Sequential AG–Carbon | 69.69 | – |  |  |  |  |  |
| Equal-task macro | GENECraft (Ours) | **74.16** | – |  |  |  |  |  |

**Table 10.**
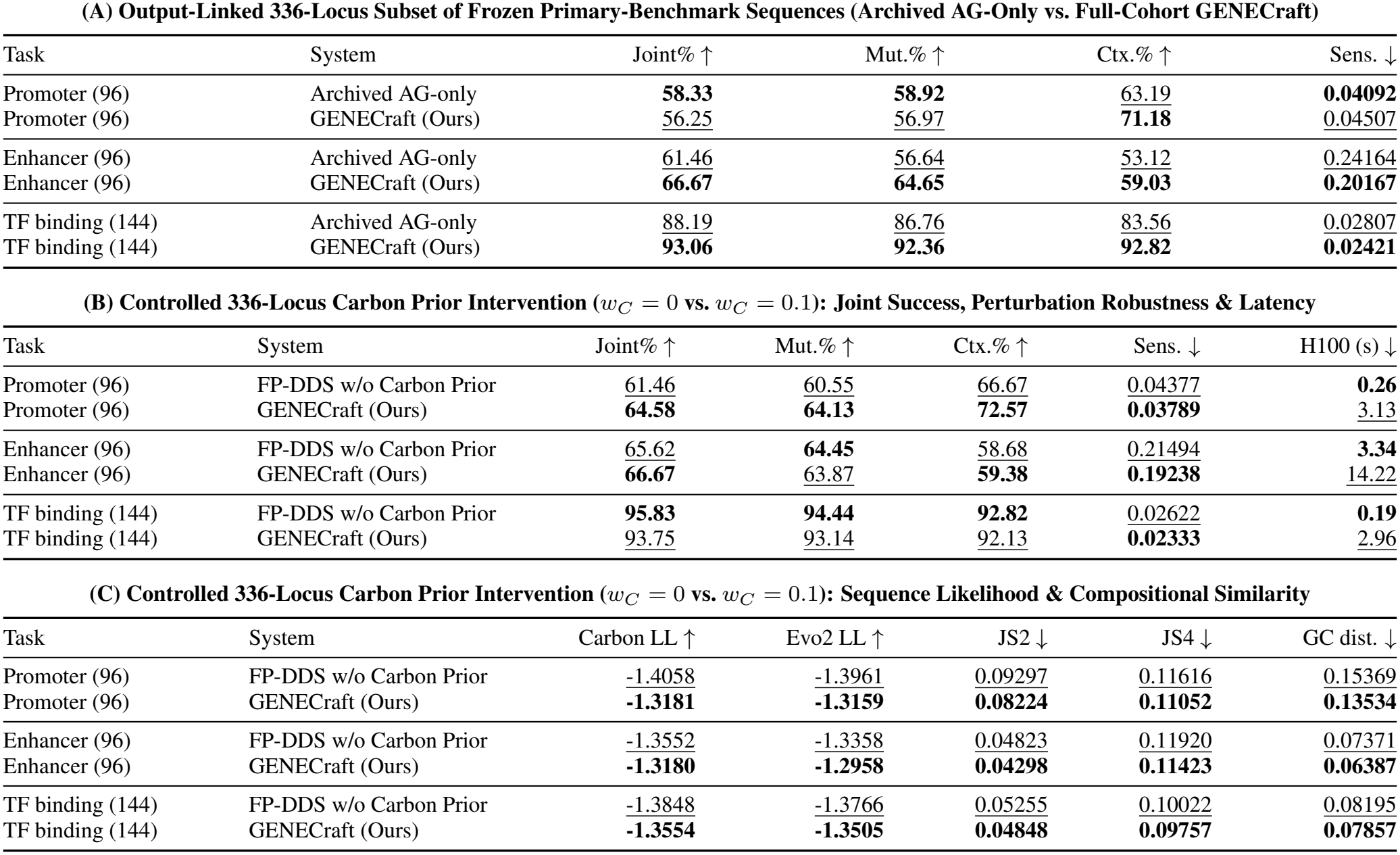
336-locus perturbation and compositional evaluation. Panel A evaluates output-linked frozen primary-benchmark sequences (2,185-locus cohort restricted to 336 loci). Panels B and C compare *w*_*C*_ = 0 versus default *w*_*C*_ = 0.1 from the independent, seed- and noise-matched intervention (*w*_*C*_ ∈ *{*0, 0.03, 0.1, 0.3*}*; effect profile in Tables 12–16). Mut. and Ctx. report full-joint success after substitutions and background swaps; Sens. measures Borzoi CAGE/binding or H3K27ac sensitivity (lower indicates higher stability). Warm H100 median latencies exclude loading/evaluation. Panel C reports log-likelihood (nats/bp), native-relative JS divergence (bits), and GC error, retaining failed designs.

| **(A) Output-Linked 336-Locus Subset of Frozen Primary-Benchmark Sequences (Archived AG-Only vs. Full-Cohort GENECraft)** |  |  |  |  |  |  |
|---|---|---|---|---|---|---|
| Task | System | Joint% ↑ | Mut.% ↑ | Ctx.% ↑ | Sens. ↓ |  |
| Promoter (96) | Archived AG-only | **58.33** | **58.92** | 63.19 | **0.04092** |  |
| Promoter (96) | GENECraft (Ours) | 56.25 | 56.97 | **71.18** | 0.04507 |  |
| Enhancer (96) | Archived AG-only | 61.46 | 56.64 | 53.12 | 0.24164 |  |
| Enhancer (96) | GENECraft (Ours) | **66.67** | **64.65** | **59.03** | **0.20167** |  |
| TF binding (144) | Archived AG-only | 88.19 | 86.76 | 83.56 | 0.02807 |  |
| TF binding (144) | GENECraft (Ours) | **93.06** | **92.36** | **92.82** | **0.02421** |  |
| **(B) Controlled 336-Locus Carbon Prior Intervention** (*w*<sub>*C*</sub> = 0 **vs.** *w*<sub>*C*</sub> = 0.1)**: Joint Success, Perturbation Robustness & Latency** |  |  |  |  |  |  |
| Task | System | Joint% ↑ | Mut.% ↑ | Ctx.% ↑ | Sens. ↓ | H100 (s) ↓ |
| Promoter (96) | FP-DDS w/o Carbon Prior | 61.46 | 60.55 | 66.67 | 0.04377 | **0.26** |
| Promoter (96) | GENECraft (Ours) | **64.58** | **64.13** | **72.57** | **0.03789** | 3.13 |
| Enhancer (96) | FP-DDS w/o Carbon Prior | 65.62 | **64.45** | 58.68 | 0.21494 | **3.34** |
| Enhancer (96) | GENECraft (Ours) | **66.67** | 63.87 | **59.38** | **0.19238** | 14.22 |
| TF binding (144) | FP-DDS w/o Carbon Prior | **95.83** | **94.44** | **92.82** | 0.02622 | **0.19** |
| TF binding (144) | GENECraft (Ours) | 93.75 | 93.14 | 92.13 | **0.02333** | 2.96 |
| **(C) Controlled 336-Locus Carbon Prior Intervention** (*w*<sub>*C*</sub> = 0 **vs.** *w*<sub>*C*</sub> = 0.1)**: Sequence Likelihood & Compositional Similarity** |  |  |  |  |  |  |
| Task | System | Carbon LL ↑ | Evo2 LL ↑ | JS2 ↓ | JS4 ↓ | GC dist. ↓ |
| Promoter (96) | FP-DDS w/o Carbon Prior | -1.4058 | -1.3961 | 0.09297 | 0.11616 | 0.15369 |
| Promoter (96) | GENECraft (Ours) | **-1.3181** | **-1.3159** | **0.08224** | **0.11052** | **0.13534** |
| Enhancer (96) | FP-DDS w/o Carbon Prior | -1.3552 | -1.3358 | 0.04823 | 0.11920 | 0.07371 |
| Enhancer (96) | GENECraft (Ours) | **-1.3180** | **-1.2958** | **0.04298** | **0.11423** | **0.06387** |
| TF binding (144) | FP-DDS w/o Carbon Prior | -1.3848 | -1.3766 | 0.05255 | 0.10022 | 0.08195 |
| TF binding (144) | GENECraft (Ours) | **-1.3554** | **-1.3505** | **0.04848** | **0.09757** | **0.07857** |

Gate-level decomposition clarifies this trade-off: because several native targets lie near compositional limits, four natural-start promoters and five activity-passing TF designs exceed the 65% GC cap (two with 14-bp homopolymers), and enhancer chromatin-risk compliance shifts from 19/24 to 17/24. Natural-start optimization still edits 29.6–38.4% of bases while improving prior likelihood and *k*-mer fidelity. Source-context cluster-bootstrap (2,000 replicates) 95% confidence intervals for the natural-minus-random full-joint success difference are [*−*25.0, 21.7], [*−*12.5, 29.2], and [*−*21.2, 6.5] percentage points for promoters, enhancers, and TFs across 13, 24, and 17 context clusters (unadjusted).

### C. Multiple Functional Oracles and Sequence Priors

GENECraft’s modular nucleotide-gradient interface directly supports multi-oracle and multi-prior ensembles, such as combining AlphaGenome and Enformer functional gradients with Carbon and Evo2 prior gradients:

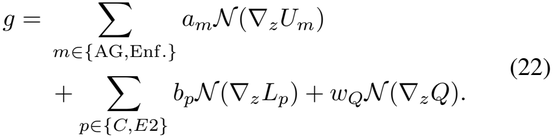

Through a unified nucleotide-space gradient interface, GENECraft’s FP-DDS framework pairs diverse functional predictors and genomic priors under identical discrete dynamics. All evaluated runs apply synchronized every-step two-model guidance (Equation 22 formalizes an unevaluated four-model extension), enabling controlled comparisons across enhancer (*n* = 24, 48 steps, *×*2 restarts; Supplementary Table 6) and TF (Table 4D) backbone swaps.

### D. Full-Benchmark Metric Provenance

All 21,850 manifest-verified evaluations across ten systems on 2,185 loci use a unified pipeline for native-relative *k*-mer and composition metrics. Functional readouts orient negative-strand promoters transcriptionally, whereas prior log-likelihoods average both strands; Carbon is in-loop for Carbon-guided or sampled arms, while Evo2 remains strictly external for default GENECraft.

In the archived raw-score parity audit, re-scoring archived sequences confirms numerical reproducibility, with mean signed log-likelihood differences of *−*0.000034 nats/bp for Carbon and +0.000002 nats/bp for Evo2 (maximum absolute deviations 0.007852 and 0.000904). Full-cohort enhancers report the re-scored AlphaGenome-only joint-success count of 287 (vs. 288 previously). Released parity records, SHA256 hashes, and scripts cover all ten systems (Tables 1–3).

#### Enhancer chromatin-risk gate

Table 7 specifies the six Enformer/Borzoi enhancer specificity checks, raw CAGE peak and mean H3K4me3 ceilings plus H3K4me1*/*(H3K4me3 + 10^*−*6^) floors, summarized by *I*_risk,*i*_ and combined with DNase/H3K27ac activity pass for full-joint success.

### E. Additional Experimental Detail

#### E.1. Benchmark Data and Candidate Generation

##### Data lineage and held-out loci

Candidate genomic loci are curated from the DNA Gen benchmark [39] (https://huggingface.co/datasets/xingyusu/DNA_Gen ) and mapped to GRCh38 with native flanking contexts. Promoter loci derive from FANTOM-CAT robust TSS annotations [21]; after flanking verification and pilot exclusion, score-blind SHA256 coordinate hashing selects 512 of 802 candidates (targeted to K562 AlphaGenome CAGE). Enhancer loci from `Genomic_MEL_regions.fa` represent human melanoma chromatin-accessibility regions [4]; from 3,324 exact-matched records (of 3,885 total), excluding 47 pilot loci and applying score-blind hashing yields 512 melanocyte-targeted test enhancers. TF-binding targets from `gm12878_500_test_df.csv` use ENCODE clustered GM12878 ChIP-seq peaks [40] strictly separated from both training and calibration splits; excluding two development factors (RUNX3 and TCF12) yields 1,161 held-out test peaks across 18 TFs (editable peak widths 65–500 bp). TF loci require an unambiguous 32, 768-bp genomic window for long-context external evaluation alongside the 4, 096-bp generation context.

##### Sequence geometry

Total context windows and central editable target lengths are 8, 192*/*256 bp for promoters, 65, 536*/*500 bp for enhancers, and 4, 096*/*65–500 bp for TF binding (preserving native peak widths). Optimization modifies only the central target region while keeping flanking contexts fixed.

##### External Readout Mapping and Threshold Provenance

In the 24-locus enhancer study, Enformer and Borzoi evaluate foreskin melanocyte male newborn DNase/H3K27ac/H3K4me3/H3K4me1 and light/dark melanocyte CAGE tracks. Under Appendix A.2 geometry, activity gains use native-relative Δ log(1 + *s*) and risk/composition gates use archived JSON cutoffs; Appendix I assesses threshold sensitivity.

##### Default Baseline Code Paths and Evaluation Accounting

Across the 2,185-locus benchmark (Tables 1–3), GENECraft uses *S* = 24, *R* = 1 (promoters/TFs) and *S* = 48, *R* = 2 (enhancers); Fast SeqProp (with learnable instance normalization *γ, β*) and AdaLead maximize *U*_AG_; and generative baselines apply best-of-64 or 128 AlphaGenome reranking alongside dinucleotide shuffling, retaining one SHA256-verified candidate per locus.

##### Perturbation and factorial ablation protocols

The perturbation study evaluates 2,352 parent sequences across seven arms: matched *w*_*C*_ ∈ *{*0, 0.03, 0.1, 0.3*}* alongside archived AG-only, wild-type-only sequential AG–Carbon, and dense GENECraft references. Six arms undergo single, double, 1%, and 5% mutational burdens plus three context swaps across 672 natural donors, yielding 48,384 mutated and 6,048 context-swapped variants (56,784 total records, retaining all failures).

Sequential refinement follows *w*_*C*_ = 0 generation with up to 8 greedy prior-favored edits (32 in deep TF), whereas coupled arms fuse AG–Carbon gradients every step (dense) or fourth step (sparse). H200 wall-clock medians exclude model loading and external evaluation.

Across 24 enhancer loci (500-bp window, 65,536-bp context, 48 steps, 2 restarts; Table 4C), we compare `GENECraft` (*w*_prior_ = 0.10, *w*_*Q*_ = 0.15) against matched ablations omitting Carbon gradients (`w/o prior` ), functional gradients (`w/o FG` ), both (`w/o guidance` ), or soft penalty (`w/o penalty` ). Removing functional gradients collapses joint success (0*/*24), whereas Carbon guidance elevates joint pass (13*/*24 to 19*/*24) by curbing external risk violations.

We crossed AlphaGenome and Enformer with Carbon and Evo2 across 24 TF cases (six TFs *×* four loci) under matched settings (24 steps, 1 restart, every-step fusion, prior weight 0.1, *w*_*Q*_ = 0.15, shared initializations/Gumbel seeds; H200 for AG+Carbon, H20 otherwise).

Evo2 used the Evo2-7B `bf16` path; Enformer+Evo2 used non-reentrant activation rematerialization with validated input gradients (0.07–0.12% cross-host relative difference, within 0.390625% half-epsilon tolerance). Selection used only the active oracle (Borzoi held out), evaluating all five models on standard TF criteria (Δ log(1 + TF peak) *>* 0.1, peak *≥* p50, Enformer/Borzoi *≥* 0.1), likelihoods (6, 144 bp flanks), and JS_2_.

Native-relative compositional divergence is quantified via dinucleotide and 4-mer Jensen–Shannon divergence [29] (base-2 logarithms, unit pseudocounts), absolute GC error, CpG error (CpG_*O/E*_ = *n*_*CG*_*L/* max(1, *n*_*C*_*n*_*G*_)), homopolymer run, maximum base fraction, compositiongate (*I*_QC_) pass, local GC variance, *k*-mer entropy, and G-quadruplex counts. Prior log-likelihoods remain diagnostics outside *J*_*i*_.

#### E.2. Sequence Quality Criteria and Rationale

Descriptive composition and hard sequence-quality metrics are evaluated on the editable target *x* of length *L*, excluding flanks. GC content is *f*_GC_(*x*) = (*n*_*C*_ + *n*_*G*_)*/L*, passing when *f*_GC_(*x*) lies within [*g*_min_, *g*_max_] (35–65% for promoters/enhancers; 30–65% for TF elements). Here *f*_max_(*x*) = max_*b*_ *n*_*b*_*/L* (*b* ∈ *{A, C, G, T}*) is the maximum single-base fraction, and *l*_HP_(*x*) is the longest contiguous homopolymer (HP) run in *x*.

Hard quality compliance is defined by the binary gate

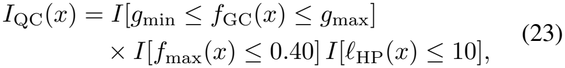

with inclusive thresholds in Table 8. Whereas *Q*(*z*) (Appendix A) supplies a differentiable guidance penalty, *I*_QC_(*x*) enforces a binary feasibility and evaluation contract. Screening GC balance, single-base dominance, and homopolymers follows standard synthetic DNA practice to mitigate synthesis, amplification, and sequencing errors [37]. Within *I*_QC_(*x*), these three operational benchmark constraints play complementary roles on the editable target; CpG *O/E* is tracked descriptively without a hard cutoff.

Diagnostic reference intervals in Tables 1–3 pair unchanged per-sequence *I*_QC_(*x*) gates (Q, defined above) with descriptive native envelopes (N) derived without new model scoring from the 2,185 benchmark targets (512 promoters, 512 enhancers, and 1,161 TF loci weighted equally across 18 TFs). Native Carbon and Evo2 log-likelihoods are recovered exactly from archived designed-minus-delta likelihoods, remaining identical across all three methods. For likelihoods and CpG *O/E*, N reports task-specific (or TF-balanced) native 5th–95th percentiles. For JS_2_ and JS_4_, N spans [0, P95] across eight deterministic nearest-length distinct same-condition/assay native donor pairs per locus, center-cropped to the shorter sequence length under unit-pseudocount base-2 JS. Because N is purely descriptive, all pass/fail gates remain unchanged.

The coordinate-hash-selected 336-locus cohort (96 promoters, 96 enhancers, 144 TFs; Table 10) supports two distinct analyses: (i) output-linked perturbation of frozen primary-benchmark AG-only and GENECraft sequences (Panel A, matching all 672 parent labels); and (ii) independent *de novo* weight interventions (*w*_*C*_ ∈ *{*0, 0.03, 0.1, 0.3*}*; Panels B–C, Table 12) under matched initializations, Gumbel noise, and hash-rotated same-GPU execution.

**Table 11.** Factorial architecture study. Full-joint success counts across 72 loci (three 24-locus folds) spanning the functional-oracle, sequence-prior, refinement-depth, and guidance-frequency axes.

| Configuration | Fold 0 | Fold 1 | Fold 2 | Total/72 |
|---|---|---|---|---|
| AG-only | 20/24 | 16/24 | 19/24 | 55/72 |
| AG + Carbon, sequential (8) | 19/24 | **17/24** | 18/24 | 54/72 |
| AG + Carbon, sequential (32) | 19/24 | **17/24** | **20/24** | **56/72** |
| AG + Evo2, sequential (8) | **21/24** | 16/24 | 19/24 | **56/72** |
| Enformer-only | 11/24 | 9/24 | 8/24 | 28/72 |
| Enformer + Carbon, sequential (8) | 10/24 | 10/24 | 8/24 | 28/72 |
| Enformer + Evo2, sequential (8) | 11/24 | 10/24 | 8/24 | 29/72 |
| AG + Carbon, every fourth step | 20/24 | 15/24 | 17/24 | 52/72 |
| GENECraft (Ours), every step | 20/24 | 16/24 | **20/24** | **56/72** |

**Table 12.**
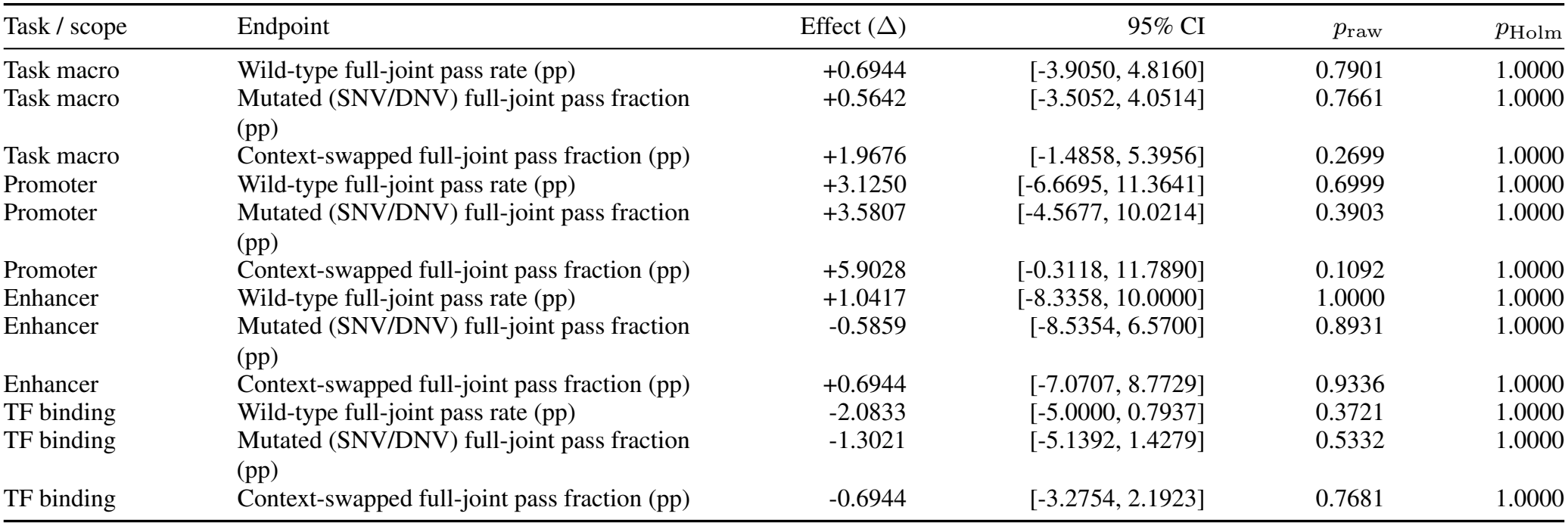
Confirmatory functional-success contrasts evaluating full-joint gate satisfaction across wild-type designs, pooled SNV/DNV substitutions, and swapped genomic contexts.

**Table 13.** Confirmatory evolutionary-plausibility contrasts evaluating bidirectional target-only conditional log-likelihoods under Carbon and external Evo2 across native flanking contexts.

| Task / scope | Endpoint | Effect (Δ) | 95% CI | *p*<sub>raw</sub> | *p*<sub>Holm</sub> |
|---|---|---|---|---|---|
| Promoter | Carbon log-likelihood (nats/bp) | +0.0877 | [0.0837, 0.0913] | 1.00 × 10<sup>−4</sup> | 0.0012 |
| Promoter | Evo2 log-likelihood (nats/bp) | +0.0802 | [0.0755, 0.0862] | 1.00 × 10<sup>−4</sup> | 0.0012 |
| Enhancer | Carbon log-likelihood (nats/bp) | +0.0372 | [0.0334, 0.0413] | 1.00 × 10<sup>−4</sup> | 0.0012 |
| Enhancer | Evo2 log-likelihood (nats/bp) | +0.0401 | [0.0356, 0.0445] | 1.00 × 10<sup>−4</sup> | 0.0012 |
| TF binding | Carbon log-likelihood (nats/bp) | +0.0293 | [0.0229, 0.0335] | 1.00 × 10<sup>−4</sup> | 0.0012 |
| TF binding | Evo2 log-likelihood (nats/bp) | +0.0262 | [0.0203, 0.0319] | 1.00 × 10<sup>−4</sup> | 0.0012 |

**Table 14.** Confirmatory contrasts for native-relative sequence composition distances.

| Task / scope | Endpoint | Effect (Δ) | 95% CI | *p*<sub>raw</sub> | *p*<sub>Holm</sub> |
|---|---|---|---|---|---|
| Promoter | Dinucleotide JS<sub>2</sub> divergence (bits) | -0.0107 | [-0.0155, -0.0070] | 1.00 × 10<sup>−4</sup> | 0.0024 |
| Promoter | 4-mer JS<sub>4</sub> divergence (bits) | -0.0056 | [-0.0086, -0.0030] | 0.0005 | 0.0080 |
| Promoter | Absolute native GC error | -0.0184 | [-0.0262, -0.0122] | 1.00 × 10<sup>−4</sup> | 0.0024 |
| Promoter | Longest homopolymer run (bp) | +0.5104 | [0.2319, 0.7500] | 0.0052 | 0.0520 |
| Enhancer | Dinucleotide JS<sub>2</sub> divergence (bits) | -0.0052 | [-0.0082, -0.0024] | 0.0011 | 0.0132 |
| Enhancer | 4-mer JS<sub>4</sub> divergence (bits) | -0.0050 | [-0.0082, -0.0019] | 0.0030 | 0.0330 |
| Enhancer | Absolute native GC error | -0.0098 | [-0.0157, -0.0045] | 0.0006 | 0.0090 |
| Enhancer | Longest homopolymer run (bp) | +0.1354 | [-0.0714, 0.3628] | 0.2847 | 0.4530 |
| TF binding | Dinucleotide JS<sub>2</sub> divergence (bits) | -0.0041 | [-0.0075, -0.0015] | 0.0006 | 0.0090 |
| TF binding | 4-mer JS<sub>4</sub> divergence (bits) | -0.0027 | [-0.0055, -0.0004] | 0.0242 | 0.1477 |
| TF binding | Absolute native GC error | -0.0034 | [-0.0115, 0.0017] | 0.2242 | 0.4530 |
| TF binding | Longest homopolymer run (bp) | +0.3750 | [0.1105, 0.6160] | 0.0153 | 0.1224 |

**Table 15.** Confirmatory functional-stability contrasts evaluating continuous oracle sensitivity under single/double nucleotide substitutions and swapped genomic flanking backgrounds.

| Task / scope | Endpoint | Effect (Δ) | 95% CI | *p*<sub>raw</sub> | *p*<sub>Holm</sub> |
|---|---|---|---|---|---|
| Promoter | Enformer CAGE mutation sensitivity (\|Δ\|) | -0.0057 | [-0.0256, 0.0089] | 0.6297 | 1.0000 |
| Promoter | Enformer CAGE context sensitivity (\|Δ\|) | -0.0073 | [-0.1354, 0.1257] | 0.9130 | 1.0000 |
| Promoter | Borzoi CAGE mutation sensitivity (\|Δ\|) | -0.0059 | [-0.0159, 0.0009] | 0.1964 | 1.0000 |
| Promoter | Borzoi CAGE context sensitivity (\|Δ\|) | -0.0368 | [-0.1252, 0.0467] | 0.4300 | 1.0000 |
| Enhancer | Enformer DNase mutation sensitivity (\|Δ\|) | +0.0032 | [-0.0030, 0.0093] | 0.3175 | 1.0000 |
| Enhancer | Enformer DNase context sensitivity (\|Δ\|) | -0.0279 | [-0.0585, -0.0006] | 0.0576 | 0.8474 |
| Enhancer | Enformer H3K27ac mutation sensitivity (\|Δ\|) | +0.0007 | [-0.0034, 0.0046] | 0.7409 | 1.0000 |
| Enhancer | Enformer H3K27ac context sensitivity (\|Δ\|) | -0.0194 | [-0.0414, 0.0023] | 0.0754 | 0.9801 |
| Enhancer | Borzoi DNase mutation sensitivity (\|Δ\|) | -0.0004 | [-0.0056, 0.0043] | 0.8796 | 1.0000 |
| Enhancer | Borzoi DNase context sensitivity (\|Δ\|) | -0.0257 | [-0.0457, -0.0064] | 0.0115 | 0.2185 |
| Enhancer | Borzoi H3K27ac mutation sensitivity (\|Δ\|) | −9.16 × 10<sup>−5</sup> | [-0.0023, 0.0020] | 0.9312 | 1.0000 |
| Enhancer | Borzoi H3K27ac context sensitivity (\|Δ\|) | -0.0226 | [-0.0371, -0.0094] | 0.0015 | 0.0405 |
| TF binding | Enformer TF mutation sensitivity (\|Δ\|) | -0.0014 | [-0.0060, 0.0018] | 0.3454 | 1.0000 |
| TF binding | Enformer TF context sensitivity (\|Δ\|) | -0.0201 | [-0.0685, 0.0149] | 0.2500 | 1.0000 |
| TF binding | Borzoi TF mutation sensitivity (\|Δ\|) | -0.0029 | [-0.0054, -0.0014] | 0.0019 | 0.0456 |
| TF binding | Borzoi TF context sensitivity (\|Δ\|) | -0.0389 | [-0.0812, -0.0121] | 0.0037 | 0.0851 |

**Table 16.** Confirmatory contrasts for functional specificity endpoints.

| Task / scope | Endpoint | Effect (Δ) | 95% CI | *p*<sub>raw</sub> | *p*<sub>Holm</sub> |
|---|---|---|---|---|---|
| Promoter | Enformer CAGE specificity margin | -0.0224 | [-0.1255, 0.0654] | 0.6903 | 1.0000 |
| Promoter | Borzoi CAGE specificity margin | -0.0095 | [-0.0468, 0.0302] | 0.6309 | 1.0000 |
| Enhancer | Enformer DNase specificity margin | +0.0130 | [-0.0549, 0.0842] | 0.7186 | 1.0000 |
| Enhancer | Enformer H3K27ac specificity margin | -0.0420 | [-0.1660, 0.0782] | 0.5003 | 1.0000 |
| Enhancer | Borzoi DNase specificity margin | +0.0288 | [-0.0380, 0.0972] | 0.4066 | 1.0000 |
| Enhancer | Borzoi H3K27ac specificity margin | -0.0173 | [-0.1312, 0.0892] | 0.7542 | 1.0000 |
| TF binding | Enformer TF specificity margin | +0.0075 | [-0.0340, 0.0410] | 0.8140 | 1.0000 |
| TF binding | Borzoi TF specificity margin | −6.22 × 10<sup>−5</sup> | [-0.0155, 0.0163] | 0.9937 | 1.0000 |

#### E.3. Detailed Evaluation, Statistical Inference, and Benchmark Synthesis

##### Continuous Functional Endpoints

Alongside binary success, main tables report continuous per-track gains Δ log(1+ *s*), while “activity pass” requires exceeding 0.1 on all required external tracks simultaneously. In ablations, “minimum gain” is the cohort-averaged worst-case per-sequence Δ log(1 + *s*) across required external tracks. *Functional specificity* is the target gain minus mean off-target gain across external predictors, and *functional stability* is the absolute perturbation deviation |Δ log(1 + *s*)| under mutation or context swap (lower indicates higher robustness).

##### Bidirectional Contextual Likelihood Protocol

Sequence plausibility is evaluated via bidirectional target-only conditional log-likelihood within 6, 144 bp native flanks, preserving Carbon 6-mer phase and full-vocabulary normalization. Main benchmarks, promoter sweeps (main Table 4A), and TF swaps (Table 4D) report Carbon and Evo2 likelihoods alongside JS_2_; held-out Evo2 concordance on AG+Carbon confirms cross-model plausibility gains.

##### Clustered Inference and Multiplicity Control

To account for spatial correlation, statistical inference clusters full-benchmark sequences into 510 non-overlapping genomic-context groups and matched perturbation analyses into 127 source-context groups. We compute confidence intervals via 2,000 cluster-bootstrap replicates and two-sided *p*-values using 10,000 paired cluster sign-flip permutations with a plus-one Monte Carlo correction [16, 34]. Our prespecified primary confirmatory contrast compares default GENECraft (*w*_*C*_ = 0.1) against the AlphaGenome-only ablation (*w*_*C*_ = 0). We apply the Holm–Bonferroni procedure [20] separately across five predefined endpoint families: joint success, prior likelihood, sequence composition, functional stability, and specificity. All failed designs and perturbed variants are retained (Appendix F; boldface denotes panel-wise optima).

##### Benchmark-Wide Synthesis, Latency, and Initialization Robustness

Across the 2,185-locus benchmark, GENECraft surpasses sequential AG–Carbon in task-macro joint success (74.16% vs. 69.69%, +4.48 points; Holm *p <* 0.001; Table 9) and all matched likelihood/JS2 contrasts (Tables 10–12) at 3.13 s–14.22 s median H100 latency. Native initialization preserves functionality while lowering JS2 by 22.7–56.7% (Appendix B, Table 5). Continuous Carbon guidance regularizes composition and strengthens perturbation stability (Tables 12–16).

### F. Complete Confirmatory Contrasts

Tables 12–16 report 54 prespecified contrasts comparing default GENECraft (*w*_C_ = 0.1) against *w*_C_ = 0 across 336 loci and five endpoint families, using paired cluster sign-flip tests (10,000 permutations, plus-one correction), 2,000-replicate cluster-bootstrap 95% CIs, and within-family Holm adjustment across the five full families spanning all 108 archived contrasts.

### G. Implementation Audit

All 210 primary five-model evaluation jobs (56,784 sequence records), the 120-design enhancer study, and the 96-design TF study (480 records) completed with verified SHA256 hashes, lineage bindings, and full-cohort denominators. Cluster inference respects 127 matched-cohort (510 full-benchmark) source-context groups, and latencies measure in-loop execution only.

### H. Enhancer Chromatin-Risk Decomposition and Exploratory Continuous Tests

Complementing our 336-locus confirmatory benchmark (Tables 10–12), a five-arm enhancer study (*n* = 24; Table 4C, Figure 3, Table 17) isolates Carbon (*w*_*C*_) and compositional penalty *Q*(*z*). Removing functional gradients collapses joint success from 19*/*24 to 0*/*24 (Holm *p* = 1.53 *×* 10^*−*5^), whereas Carbon raises joint pass over prior ablation (13*/*24; +25 points, Holm *p* = 0.1406) by expanding chromatin-risk compliance (14*/*24 to 21*/*24).

**Table 17.** Paired continuous chromatin-risk contrasts (24 enhancer loci). Mean differences (full GENECraft *−* ablation) across Enformer and Borzoi CAGE, H3K4me3, and H3K4me1*/*H3K4me3 readouts, reporting 2,000-replicate bootstrap 95% CIs, 10,000-permutation sign-flip *p*-values, and 12-test Holm adjustments.

|  | Target readout | Endpoint | Effect (Δ) | 95% CI | *p*<sub>raw</sub> | *p*<sub>Holm</sub> |
|---|---|---|---|---|---|---|
| w/o prior | Enformer | CAGE | -1.087 | [-1.869, -0.414] | 0.0035 | 0.0350 |
| w/o prior | Enformer | H3K4me3 | -0.083 | [-0.437, 0.331] | 0.6981 | 1.0000 |
| w/o prior | Enformer | H3K4me1/H3K4me3 | +0.487 | [0.296, 0.689] | 1.00 × 10<sup>−4</sup> | 0.0012 |
| w/o prior | Borzoi | CAGE | -0.105 | [-0.352, 0.119] | 0.4105 | 1.0000 |
| w/o prior | Borzoi | H3K4me3 | -0.001 | [-0.592, 0.665] | 0.9978 | 1.0000 |
| w/o prior | Borzoi | H3K4me1/H3K4me3 | +0.128 | [-0.001, 0.267] | 0.0731 | 0.5487 |
| w/o penalty | Enformer | CAGE | +0.272 | [0.133, 0.429] | 0.0006 | 0.0066 |
| w/o penalty | Enformer | H3K4me3 | -0.991 | [-2.186, -0.164] | 0.0158 | 0.1422 |
| w/o penalty | Enformer | H3K4me1/H3K4me3 | +0.177 | [-0.010, 0.355] | 0.0822 | 0.5487 |
| w/o penalty | Borzoi | CAGE | +0.093 | [-0.010, 0.194] | 0.1024 | 0.5487 |
| w/o penalty | Borzoi | H3K4me3 | -0.570 | [-1.365, -0.027] | 0.0686 | 0.5487 |
| w/o penalty | Borzoi | H3K4me1/H3K4me3 | -0.156 | [-0.317, 0.003] | 0.0731 | 0.5487 |

**Table 18.** External chromatin-risk gate violations (24 enhancer loci). Marginal locus failure counts per Enformer and Borzoi risk gate across five ablation arms; counts overlap at multi-violation loci.

|  | Enf.<br>CAGE | Enf.<br>H3K4me3 | Enf.<br>Ratio | Bor.<br>CAGE | Bor.<br>H3K4me3 | Bor.<br>Ratio | Any<br>failure |
|---|---|---|---|---|---|---|---|
| GENECraft (Ours) | **0** | **2** | **2** | <u>3</u> | **2** | **2** | **3** |
| w/o prior (AG-only) | 6 | <u>3</u> | **2** | 4 | <u>3</u> | **2** | 10 |
| w/o FG (Carbon-only) | 15 | **2** | <u>7</u> | 18 | <u>3</u> | <u>4</u> | 21 |
| w/o guidance | <u>4</u> | <u>3</u> | 14 | 7 | <u>3</u> | <u>4</u> | 16 |
| w/o penalty | **0** | <u>3</u> | **2** | **2** | 4 | **2** | <u>5</u> |

Decomposing the six external risk gates reveals complementary roles (Table 17). Removing the Carbon prior lowers risk compliance from 21*/*24 to 14*/*24 via Enformer and Borzoi CAGE violations, whereas Carbon significantly lowers Enformer CAGE peaks (difference = *−*1.087, Holm *p* = 0.0350) and raises H3K4me1*/*H3K4me3 (+0.487, Holm *p* = 0.0012). Conversely, *Q*(*z*) curbs H3K4me3 excursions while maintaining 100% hard QC compliance.

### I. Post-Hoc Threshold Sensitivity Analysis

We evaluated post-hoc threshold sensitivity on the locked candidate sequences while holding generation and the common AlphaGenome selection gate fixed (Figures 5–7).

**Figure 5.**
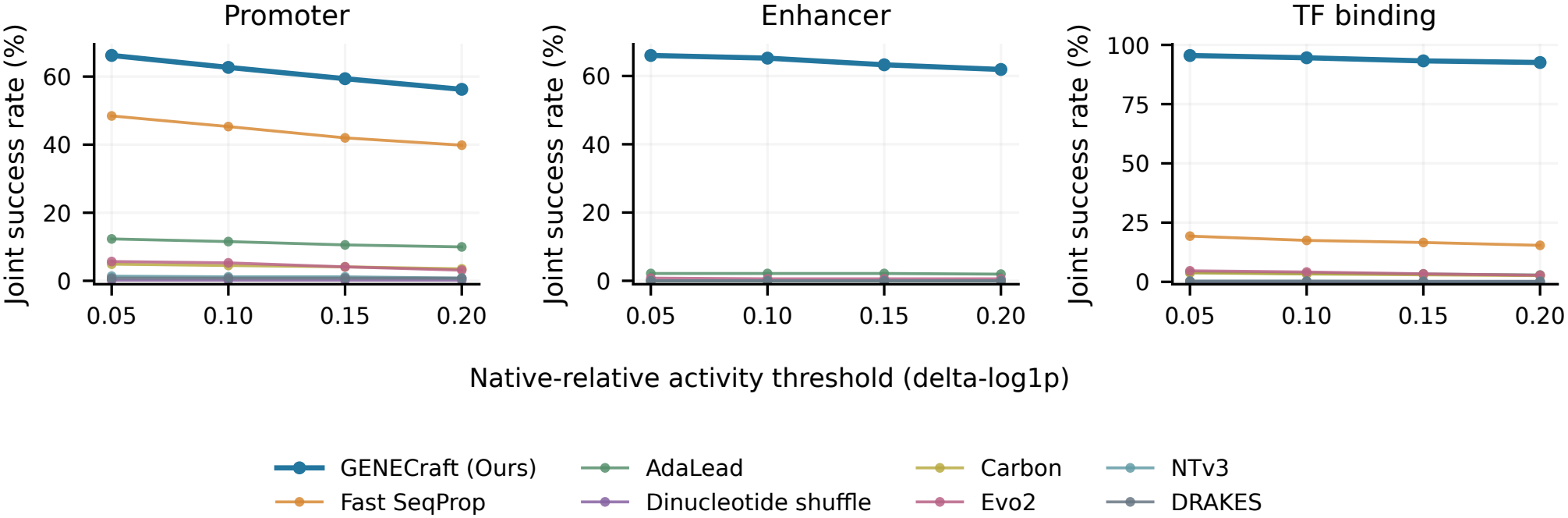
Cross-oracle activity-threshold sensitivity (2,185 loci). Full-joint pass rates for locked candidate sequences across Δ log(1 + *s*) cutoffs (*γ* ∈ *{*0.05, 0.10, 0.15, 0.20*}*) for GENECraft and seven baselines.

Sweeping the external native-relative activity-gain cutoff Δ log(1 + *s*) *≥ γ* across *γ* ∈ *{*0.05, 0.10, 0.15, 0.20*}* on the 2,185-locus benchmark (Figure 5), GENECraft maintains higher full-joint success than all seven external baselines across every cutoff (0.05, 0.10, 0.15, 0.20) and task.

Scaling each external enhancer risk threshold (CAGE ceiling, H3K4me3 ceiling, and H3K4me1*/*(H3K4me3 + 10^*−*6^) ratio floor) by *m* ∈ *{*0.8, 0.9, 1.0, 1.1, 1.2*}* across the 24-locus cohort (Figure 6; 15 settings) shows GENECraft consistently outperforming *w/o prior* by 16.67–29.17 percentage points in joint success. GENECraft matches or exceeds *w/o penalty* (*Q*(*z*) omitted) everywhere except under tightened H3K4me3 ceilings (0.8*×*, 0.9*×*).

**Figure 6.**
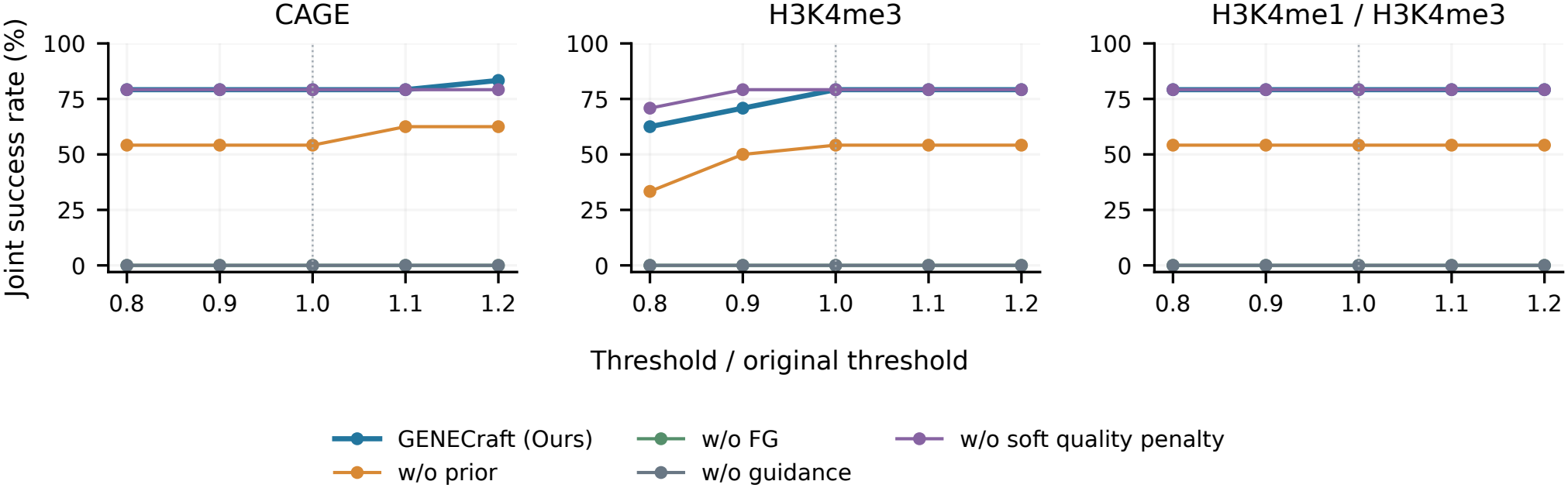
Enhancer risk-threshold sensitivity (24 loci). Full-joint pass rates across five ablation arms when scaling individual external chromatin-risk gate families by *{*0.8, 0.9, 1.0, 1.1, 1.2*}*.

Rescoring *I*_QC_ post-hoc under expanded GC bounds and varied homopolymer caps (Figure 7), with the generation-stage AlphaGenome gate fixed, likewise preserves GENECraft’s full-joint lead over all baselines.

**Figure 7.**
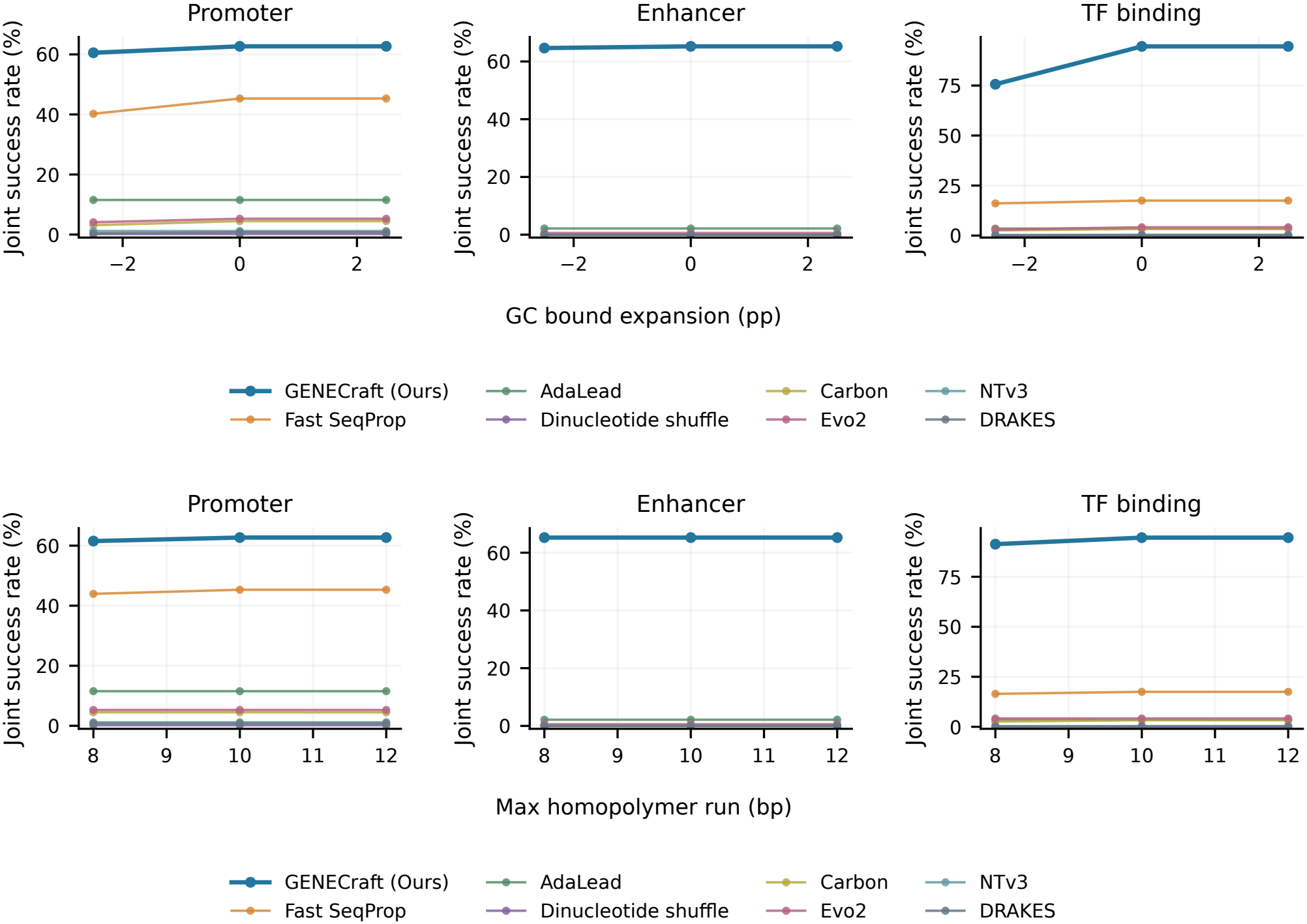
Sequence-quality threshold sensitivity (2,185 loci). Post-hoc full-joint pass rates for locked designs when shifting evaluation GC bounds by *±*2.5 percentage points and varying homopolymer caps (*L*_max_ ∈ *{*8, 10, 12*}*) under a fixed AlphaGenome feasibility gate.

### J. Per-Transcription-Factor Performance Breakdown

Table 19 details per-factor performance across all 18 GM12878 transcription factors (1,161 loci; *n* = 17 to *n* = 156, retaining all failures). To weight factors equally across varying peak counts, Table 3 reports macro-averages (94.56% full-joint success) alongside the 1,119*/*1,161 = 96.38% locus-level micro-average; this full cohort is distinct from the 6-TF, 24-locus backbone-swap subset in Table 4D. GENECraft achieves 100.00% full-joint success on 6/18 TFs, *≥* 95.00% on 13/18, and 79.25%–87.50% on the remaining 5, with 100.00% composition compliance (*I*_QC_) on 14/18 TFs and positive mean Enformer and Borzoi Δ log(1 + *s*) gains on every factor. Conversely, Fast Se-qProp reaches only 2.65%–35.26% full-joint success and yields negative mean Borzoi gains on 11/18 TFs.

**Table 19.**
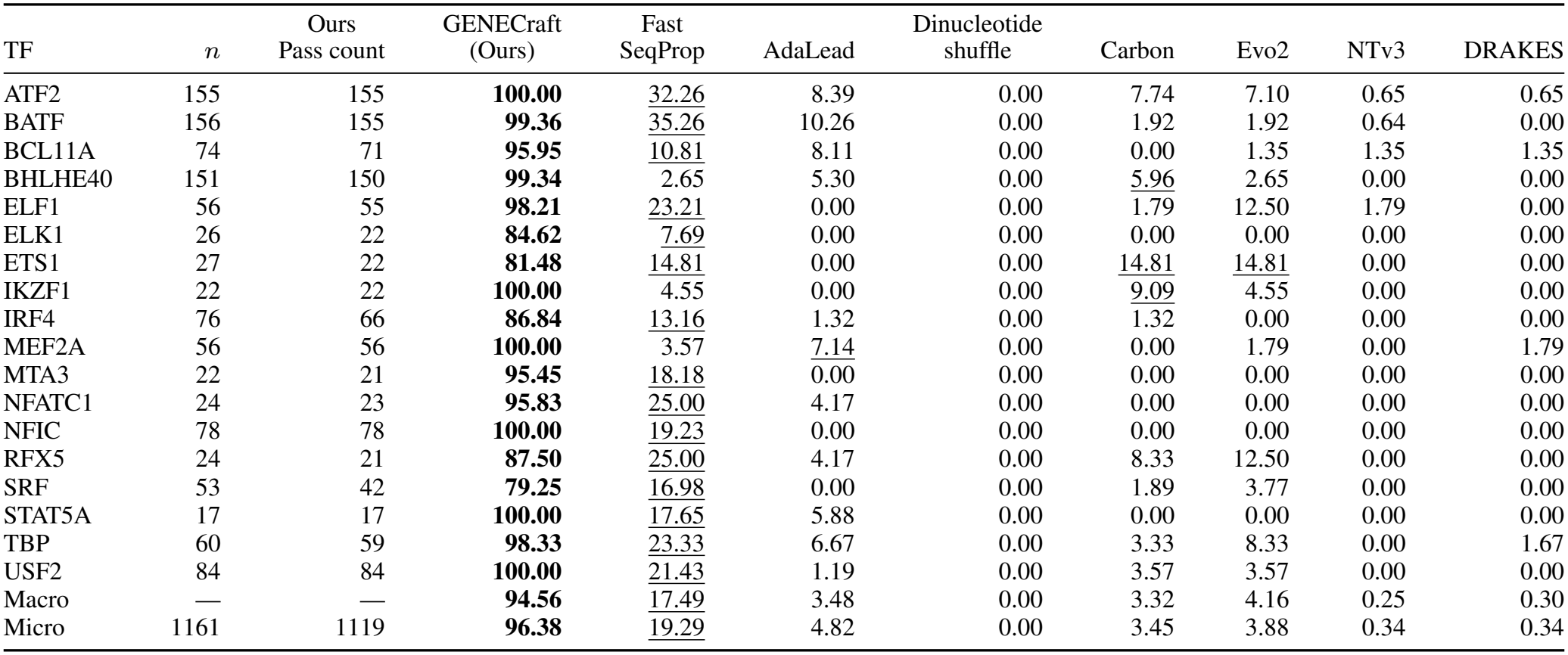
Per-factor GM12878 TF benchmark results. Locus counts and per-TF full-joint pass rates alongside equal-factor macro-averages and locus-weighted micro-averages across 1,161 binding loci and eight methods.

| TF | *n* | Ours<br>Pass count | GENECraft<br>(Ours) | Fast<br>SeqProp | AdaLead | Dinucleotide<br>shuffle | Carbon | Evo2 | NTv3 | DRAKES |
|---|---|---|---|---|---|---|---|---|---|---|
| ATF2 | 155 | 155 | **100.00** | 32.26 | 8.39 | 0.00 | 7.74 | 7.10 | 0.65 | 0.65 |
| BATF | 156 | 155 | **99.36** | 35.26 | 10.26 | 0.00 | 1.92 | 1.92 | 0.64 | 0.00 |
| BCL11A | 74 | 71 | **95.95** | 10.81 | 8.11 | 0.00 | 0.00 | 1.35 | 1.35 | 1.35 |
| BHLHE40 | 151 | 150 | **99.34** | 2.65 | 5.30 | 0.00 | 5.96 | 2.65 | 0.00 | 0.00 |
| ELF1 | 56 | 55 | **98.21** | 23.21 | 0.00 | 0.00 | 1.79 | 12.50 | 1.79 | 0.00 |
| ELK1 | 26 | 22 | **84.62** | 7.69 | 0.00 | 0.00 | 0.00 | 0.00 | 0.00 | 0.00 |
| ETS1 | 27 | 22 | **81.48** | 14.81 | 0.00 | 0.00 | 14.81 | 14.81 | 0.00 | 0.00 |
| IKZF1 | 22 | 22 | **100.00** | 4.55 | 0.00 | 0.00 | 9.09 | 4.55 | 0.00 | 0.00 |
| IRF4 | 76 | 66 | **86.84** | 13.16 | 1.32 | 0.00 | 1.32 | 0.00 | 0.00 | 0.00 |
| MEF2A | 56 | 56 | **100.00** | 3.57 | 7.14 | 0.00 | 0.00 | 1.79 | 0.00 | 1.79 |
| MTA3 | 22 | 21 | **95.45** | 18.18 | 0.00 | 0.00 | 0.00 | 0.00 | 0.00 | 0.00 |
| NFATC1 | 24 | 23 | **95.83** | 25.00 | 4.17 | 0.00 | 0.00 | 0.00 | 0.00 | 0.00 |
| NFIC | 78 | 78 | **100.00** | 19.23 | 0.00 | 0.00 | 0.00 | 0.00 | 0.00 | 0.00 |
| RFX5 | 24 | 21 | **87.50** | 25.00 | 4.17 | 0.00 | 8.33 | 12.50 | 0.00 | 0.00 |
| SRF | 53 | 42 | **79.25** | 16.98 | 0.00 | 0.00 | 1.89 | 3.77 | 0.00 | 0.00 |
| STAT5A | 17 | 17 | **100.00** | 17.65 | 5.88 | 0.00 | 0.00 | 0.00 | 0.00 | 0.00 |
| TBP | 60 | 59 | **98.33** | 23.33 | 6.67 | 0.00 | 3.33 | 8.33 | 0.00 | 1.67 |
| USF2 | 84 | 84 | **100.00** | 21.43 | 1.19 | 0.00 | 3.57 | 3.57 | 0.00 | 0.00 |
| Macro | — | — | **94.56** | 17.49 | 3.48 | 0.00 | 3.32 | 4.16 | 0.25 | 0.30 |
| Micro | 1161 | 1119 | **96.38** | 19.29 | 4.82 | 0.00 | 3.45 | 3.88 | 0.34 | 0.34 |

### K. Archival Two-Stage Study of Fixed versus Resampled Gumbel Noise

Table 20 reports an archival paired study (*n* = 50 promoters, *S* = 24, *R* = 1; *n* = 50 enhancers, *S* = 96, *R* = 4) comparing fixed per-restart against step-resampled Gumbel noise in an earlier two-stage pipeline (unregularized AlphaGenome generation then post-hoc Carbon refinement). Fixed noise raised single-seed joint passes from 28*/*50 to 31*/*50 (promoters) and 13*/*50 to 17*/*50 (enhancers), motivating deterministic decoding.

**Table 20.** Gumbel-noise diagnostics in sequential AG–Carbon. Final-pass counts comparing fixed versus step-resampled Gumbel noise across 50 promoters (*S* = 24, *R* = 1) and 50 enhancers (*S* = 96, *R* = 4).

| Task / scope | *n* | Steps (*S*) | Restarts (*R*) | Fresh<br>pass | Fixed<br>pass | Fixed-only<br>pass | Fresh-only<br>pass |
|---|---|---|---|---|---|---|---|
| Promoter | 50 | 24 | 1 | 28/50 | **31/50** | 5 | 2 |
| Enhancer | 50 | 96 | 4 | 13/50 | **17/50** | 5 | 1 |

### L. Shared-Component Controller Controls

To isolate the FP-DDS discrete dynamical controller from shared objective components, a five-arm study on the 24-locus enhancer cohort (*S* = 48, *R* = 2; Tables 26, 21–23) ran in rotated locus order. Every arm shares utility *U*_AG_, feasibility gate *F* (*x*; *c, t*) (with *I*_QC_), feasibility-first selection, and held-out Enformer/Borzoi thresholds, retaining all 120 final designs and 360 prefix readouts (24, 48, 96 backward calls).

**Table 21.**
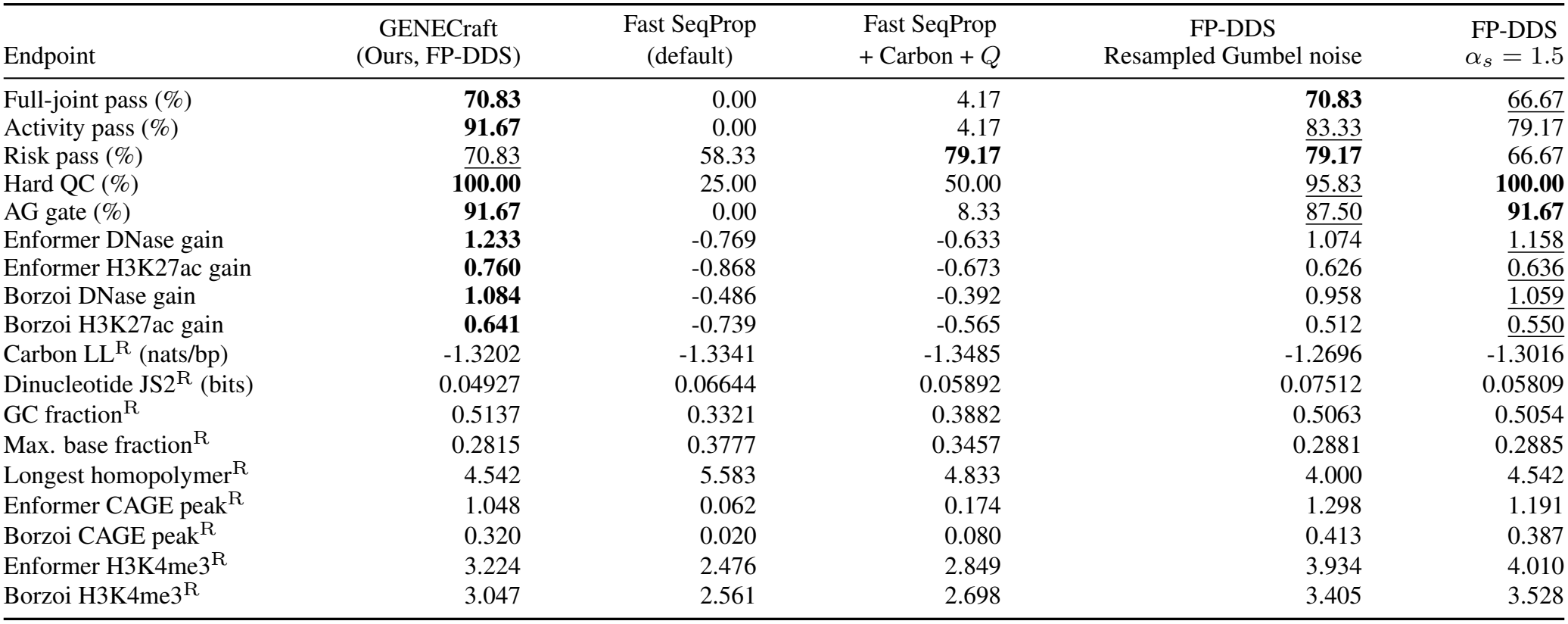
Matched five-arm controller benchmark across 24 enhancer loci (*S* = 48, *R* = 2), reporting joint/marginal pass rates and external Enformer/Borzoi activity gains (Δ log(1 + *s*)); ^R^ denotes unranked reference diagnostics.

| Endpoint | GENECraft<br>(Ours, FP-DDS) | Fast SeqProp<br>(default) | Fast SeqProp<br>+ Carbon + *Q* | FP-DDS<br>Resampled Gumbel noise | FP-DDS<br>α<sub>s</sub> = 1.5 |
|---|---|---|---|---|---|
| Full-joint pass (%) | **70.83** | 0.00 | 4.17 | **70.83** | 66.67 |
| Activity pass (%) | **91.67** | 0.00 | 4.17 | 83.33 | 79.17 |
| Risk pass (%) | 70.83 | 58.33 | **79.17** | **79.17** | 66.67 |
| Hard QC (%) | **100.00** | 25.00 | 50.00 | 95.83 | **100.00** |
| AG gate (%) | **91.67** | 0.00 | 8.33 | 87.50 | **91.67** |
| Enformer DNase gain | **1.233** | -0.769 | -0.633 | 1.074 | 1.158 |
| Enformer H3K27ac gain | **0.760** | -0.868 | -0.673 | 0.626 | 0.636 |
| Borzoi DNase gain | **1.084** | -0.486 | -0.392 | 0.958 | 1.059 |
| Borzoi H3K27ac gain | **0.641** | -0.739 | -0.565 | 0.512 | 0.550 |
| Carbon LL<sup>R</sup> (nats/bp) | -1.3202 | -1.3341 | -1.3485 | -1.2696 | -1.3016 |
| Dinucleotide JS2<sup>R</sup> (bits) | 0.04927 | 0.06644 | 0.05892 | 0.07512 | 0.05809 |
| GC fraction<sup>R</sup> | 0.5137 | 0.3321 | 0.3882 | 0.5063 | 0.5054 |
| Max. base fraction<sup>R</sup> | 0.2815 | 0.3777 | 0.3457 | 0.2881 | 0.2885 |
| Longest homopolymer<sup>R</sup> | 4.542 | 5.583 | 4.833 | 4.000 | 4.542 |
| Enformer CAGE peak<sup>R</sup> | 1.048 | 0.062 | 0.174 | 1.298 | 1.191 |
| Borzoi CAGE peak<sup>R</sup> | 0.320 | 0.020 | 0.080 | 0.413 | 0.387 |
| Enformer H3K4me3<sup>R</sup> | 3.224 | 2.476 | 2.849 | 3.934 | 4.010 |
| Borzoi H3K4me3<sup>R</sup> | 3.047 | 2.561 | 2.698 | 3.405 | 3.528 |

Under 96 AlphaGenome backward calls and shared gate *F* (Table 26), all three FP-DDS arms (canonical, resampled noise, constant step *α*_*s*_ *≡* 1.5) use coordinate-seeded discrete initialization *x*_0_ (*z*_1_ = 4 OneHot(*x*_0_)). Both Fast SeqProp arms retain Gaussian initialization *N* (0, 1), instance normalization (*γ, β*), straight-through sampling, and Adam (lr = 0.1; seed 0 + restart); the augmented arm adds RMS-normalized Carbon (*w*_*C*_ = 0.1) and *Q* (*w*_*Q*_ = 0.15) post-normalization forces before backpropagating through (*γ, β*).

Across the 24 enhancer loci (Table 4B), all arms execute 96 AlphaGenome backward passes per locus, and prior-guided arms add 192 Carbon gradient evaluations (96 steps *×* 2 strands). At matched median warm runtime (46.27 s vs. 46.17 s; Table 23), FP-DDS increases full-joint success by +70.83 pp over default FSP (Holm *p* = 6.10 *×* 10^*−*5^) and +66.67 pp over FSP + Carbon + *Q* (Holm *p* = 9.16 *×* 10^*−*5^; Table 22).

**Table 22.** Paired binary contrasts of full-joint success against FP-DDS across 24 enhancer loci, reporting discordant counts, percentage-point differences, 95% bootstrap confidence intervals (2,000 resamples), exact sign-flip *p*-values, and Holm-adjusted *p*-values.

| Configuration | FP-DDS only | Other only | Δ (pp) | 95% CI | *p*<sub>raw</sub> | *p*<sub>Holm</sub> |
|---|---|---|---|---|---|---|
| Fast SeqProp (default) | 17 | 0 | +70.83 | [54.17, 87.50] | 1.53 × 10<sup>−5</sup> | 6.10 × 10<sup>−5</sup> |
| Fast SeqProp + Carbon + *Q* | 16 | 0 | +66.67 | [45.83, 83.33] | 3.05 × 10<sup>−5</sup> | 9.16 × 10<sup>−5</sup> |
| FP-DDS (resampled noise) | 3 | 3 | +0.00 | [-20.83, 20.83] | 1.0000 | 1.0000 |
| FP-DDS (constant step, *α*<sub>*s*</sub> = 1.5) | 5 | 4 | +4.17 | [-20.83, 29.17] | 1.0000 | 1.0000 |

**Table 23.**
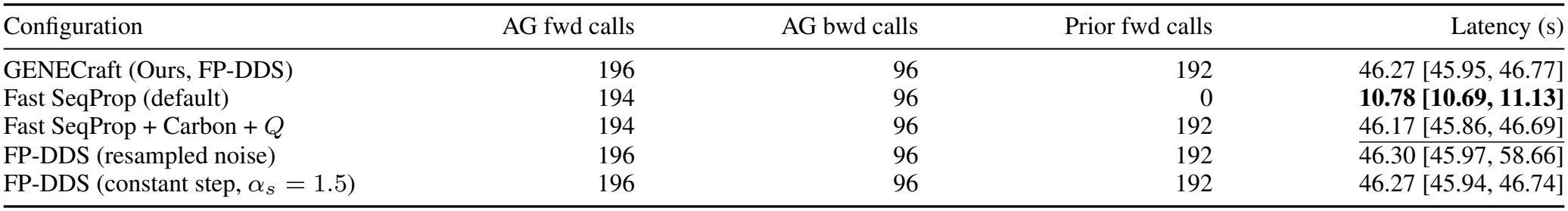
Per-locus inference call counts and warm generation latency (seconds; median and range across *n* = 24 loci) on dedicated H20 GPUs, excluding model loading, warmup, and post-hoc evaluation.

| Configuration | AG fwd calls | AG bwd calls | Prior fwd calls | Latency (s) |
|---|---|---|---|---|
| GENECraft (Ours, FP-DDS) | 196 | 96 | 192 | 46.27 [45.95, 46.77] |
| Fast SeqProp (default) | 194 | 96 | 0 | **10.78 [10.69, 11.13]** |
| Fast SeqProp + Carbon + *Q* | 194 | 96 | 192 | 46.17 [45.86, 46.69] |
| FP-DDS (resampled noise) | 196 | 96 | 192 | 46.30 [45.97, 58.66] |
| FP-DDS (constant step, α<sub>s</sub> = 1.5) | 196 | 96 | 192 | 46.27 [45.94, 46.74] |

Comparing the primary reference run (19*/*24 joint pass; Table 4C) and the independent controller-suite FP-DDS repeat (17*/*24; Table 21) across all 24 enhancer loci localizes the two-locus discrepancy to external risk gates (Table 32B). Hard QC (24*/*24 → 24*/*24) and AlphaGenome compliance (22*/*24 → 22*/*24) remain invariant, cross-oracle activity rises from 20*/*24 to 22*/*24, and six-gate risk compliance shifts from 21*/*24 to 17*/*24.

### M. Temporal Schedule and Noise Dynamics

At the 96-call horizon (Tables 21–25), binary full-joint success does not significantly separate standard FP-DDS (*α*_*s*_ = 3*s/*(*S* + 1)), step-resampled Gumbel noise (Holm *p* = 1.0), and constant step size *α*_*s*_ *≡* 1.5 (Holm *p* = 1.0). However, standard FP-DDS maximizes AlphaGenome utility, Enformer/Borzoi functional gains, and dinucleotide fidelity, whereas resampled noise and constant step size favor higher Carbon log-likelihood.

Prefix readouts at 24, 48, and 96 backward calls along the *S* = 48, *R* = 2 trajectories (Table 25) reveal distinct dynamics: standard FP-DDS maintains 17*/*24 → 17*/*24 → 17*/*24 full-joint success while expanding external activity (19*/*24 to 22*/*24), resampled noise improves monotonically (12*/*24 → 16*/*24 → 17*/*24), and constant step size fluctuates non-monotonically (14*/*24 → 19*/*24 → 16*/*24) as chromatin-risk compliance declines.

### N. Reference Optimizer Interfaces

Default Fast SeqProp initializes logits from *N* (0, 1) (seed = 0), applies 1D instance normalization (*γ* = 1, *β* = 0) and straight-through sampling (samples per step = 1), and runs 24 Adam steps (lr = 0.1, *β*_1_ = 0.9, *β*_2_ = 0.999, *ϵ* = 10^*−*8^; *S* = 48, *R* = 2, seed = 0 + restart in matched controller runs). AdaLead uses a 128-query budget with population 16, mutation rate 1.0 (1*/L*), threshold *τ* = 0.05, and recombination 0.0. Both record earliest-visit top feasible and overall candidates via `BestTracker` (unconstrained in `HardDNAOracle` ), excluding Enformer and Borzoi outside explicit model-swap ablations.

### O. Methodological and Empirical Distinctions Between GENECraft and Fast SeqProp

Unlike Fast SeqProp [30], GENECRAFT integrates AlphaGenome, the Carbon prior via our nucleotide-to-6-mer bridge (Appendix A.1), and differentiable quality guidance (*Q*) using RMS-normalized forces within clipped fixed-noise Gumbel trajectories (Tables 24–26). Under a matched 96-backward-pass budget across 24 enhancer loci (Tables 21– 23; Appendix M), GENECRAFT reaches 17*/*24 full-joint success versus 0*/*24 (base) and 1*/*24 (Carbon+quality-augmented Fast SeqProp; Holm *p* = 9.16 *×* 10^*−*5^) at matched runtime.

**Table 24.**
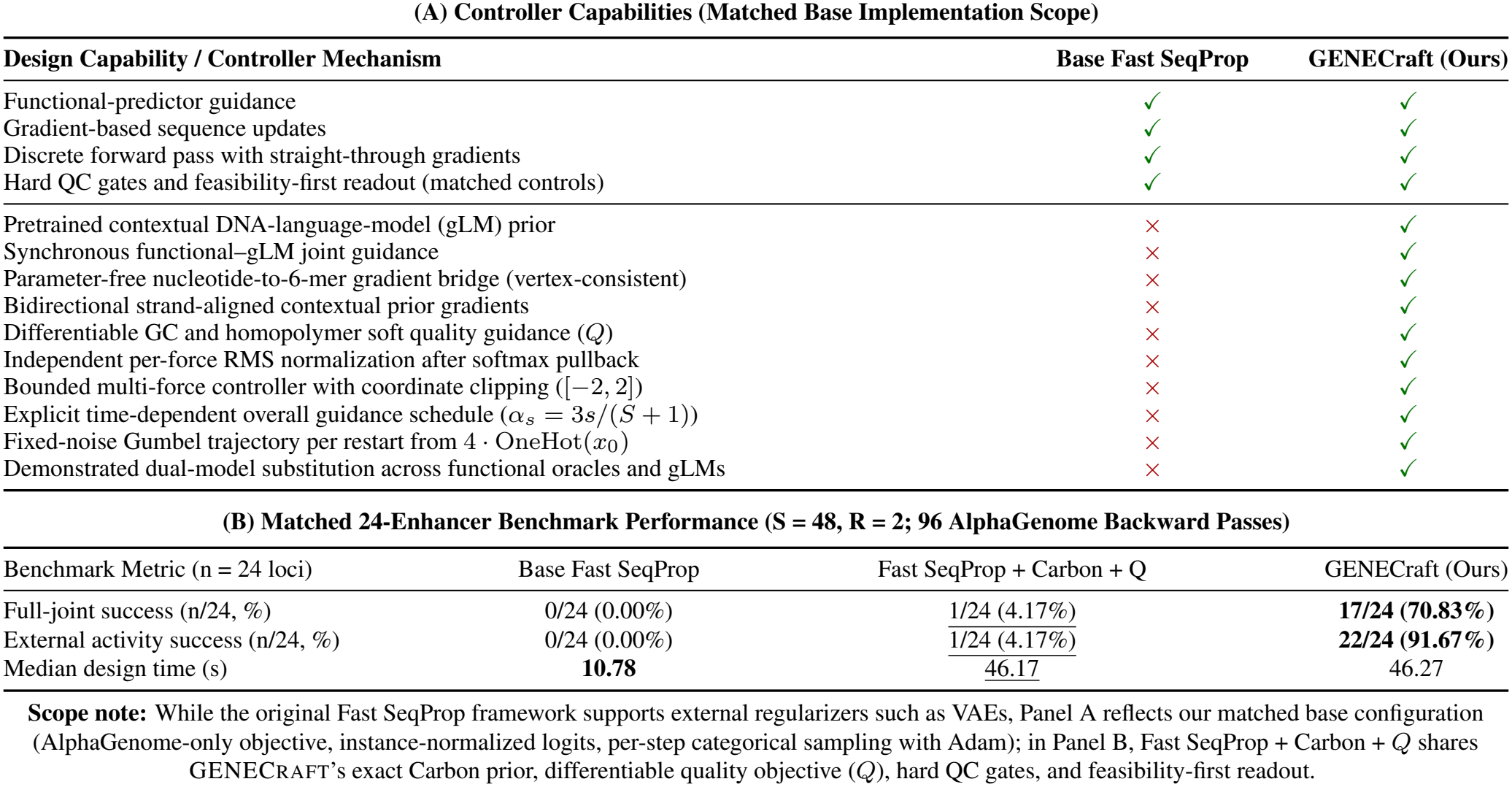
Architectural comparison (Panel A) and matched 24-enhancer benchmark (Panel B) for Fast SeqProp [30] and GENECRAFT under a 96-call AlphaGenome backward budget (*S* = 48, *R* = 2), including an augmented Fast SeqProp + Carbon + *Q* control (Tables 21–23; modularity in Table 4D).

| **(A) Controller Capabilities (Matched Base Implementation Scope)** |  |  |  |
|---|---|---|---|
| **Design Capability / Controller Mechanism** | **Base Fast SeqProp** | **GENECraft (Ours)** |  |
| Functional-predictor guidance | ✓ | ✓ |  |
| Gradient-based sequence updates | ✓ | ✓ |  |
| Discrete forward pass with straight-through gradients | ✓ | ✓ |  |
| Hard QC gates and feasibility-first readout (matched controls) | ✓ | ✓ |  |
| Pretrained contextual DNA-language-model (gLM) prior | × | ✓ |  |
| Synchronous functional–gLM joint guidance | × | ✓ |  |
| Parameter-free nucleotide-to-6-mer gradient bridge (vertex-consistent) | × | ✓ |  |
| Bidirectional strand-aligned contextual prior gradients | × | ✓ |  |
| Differentiable GC and homopolymer soft quality guidance (*Q*) | × | ✓ |  |
| Independent per-force RMS normalization after softmax pullback | × | ✓ |  |
| Bounded multi-force controller with coordinate clipping ([−2, 2]) | × | ✓ |  |
| Explicit time-dependent overall guidance schedule (*α*<sub>*s*</sub> = 3*s*/(*S* + 1)) | × | ✓ |  |
| Fixed-noise Gumbel trajectory per restart from 4 · OneHot(*x*<sub>0</sub>) | × | ✓ |  |
| Demonstrated dual-model substitution across functional oracles and gLMs | × | ✓ |  |
| **(B) Matched 24-Enhancer Benchmark Performance (S = 48, R = 2; 96 AlphaGenome Backward Passes)** |  |  |  |
| Benchmark Metric (n = 24 loci) | Base Fast SeqProp | Fast SeqProp + Carbon + Q | GENECraft (Ours) |
| Full-joint success (n/24, %) | 0/24 (0.00%) | 1/24 (4.17%) | **17/24 (70.83%)** |
| External activity success (n/24, %) | 0/24 (0.00%) | 1/24 (4.17%) | **22/24 (91.67%)** |
| Median design time (s) | **10.78** | 46.17 | 46.27 |

**Scope note:** While the original Fast SeqProp framework supports external regularizers such as VAEs, Panel A reflects our matched base configuration (AlphaGenome-only objective, instance-normalized logits, per-step categorical sampling with Adam); in Panel B, Fast SeqProp + Carbon + *Q* shares GENECraft's exact Carbon prior, differentiable quality objective (*Q*), hard QC gates, and feasibility-first readout.

**Table 25.**
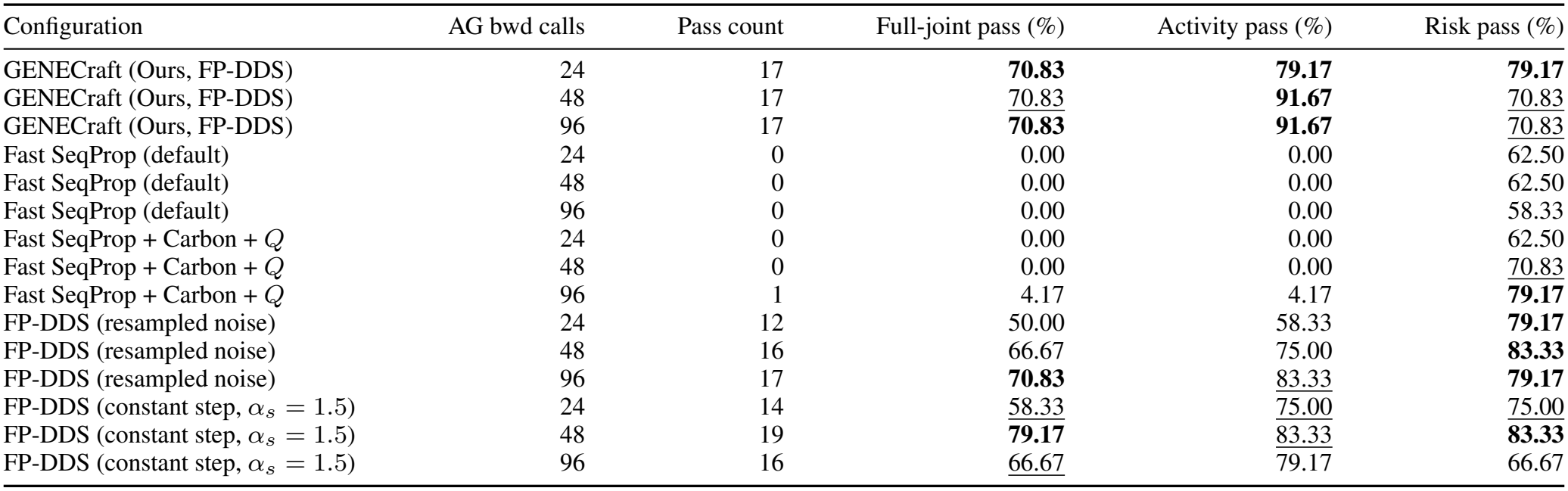
Anytime prefix readout across 15 arm–checkpoint combinations at 24, 48, and 96 cumulative AlphaGenome backward calls (*n* = 24 enhancers, locked *S* = 48, *R* = 2 trajectories) under feasibility-first exact readout.

| Configuration | AG bwd calls | Pass count | Full-joint pass (%) | Activity pass (%) | Risk pass (%) |
|---|---|---|---|---|---|
| GENECraft (Ours, FP-DDS) | 24 | 17 | **70.83** | **79.17** | **79.17** |
| GENECraft (Ours, FP-DDS) | 48 | 17 | 70.83 | **91.67** | 70.83 |
| GENECraft (Ours, FP-DDS) | 96 | 17 | **70.83** | **91.67** | 70.83 |
| Fast SeqProp (default) | 24 | 0 | 0.00 | 0.00 | 62.50 |
| Fast SeqProp (default) | 48 | 0 | 0.00 | 0.00 | 62.50 |
| Fast SeqProp (default) | 96 | 0 | 0.00 | 0.00 | 58.33 |
| Fast SeqProp + Carbon + *Q* | 24 | 0 | 0.00 | 0.00 | 62.50 |
| Fast SeqProp + Carbon + *Q* | 48 | 0 | 0.00 | 0.00 | 70.83 |
| Fast SeqProp + Carbon + *Q* | 96 | 1 | 4.17 | 4.17 | **79.17** |
| FP-DDS (resampled noise) | 24 | 12 | 50.00 | 58.33 | **79.17** |
| FP-DDS (resampled noise) | 48 | 16 | 66.67 | 75.00 | **83.33** |
| FP-DDS (resampled noise) | 96 | 17 | **70.83** | 83.33 | **79.17** |
| FP-DDS (constant step, α<sub>s</sub> = 1.5) | 24 | 14 | 58.33 | 75.00 | 75.00 |
| FP-DDS (constant step, α<sub>s</sub> = 1.5) | 48 | 19 | **79.17** | 83.33 | **83.33** |
| FP-DDS (constant step, α<sub>s</sub> = 1.5) | 96 | 16 | 66.67 | 79.17 | 66.67 |

**Table 26.**
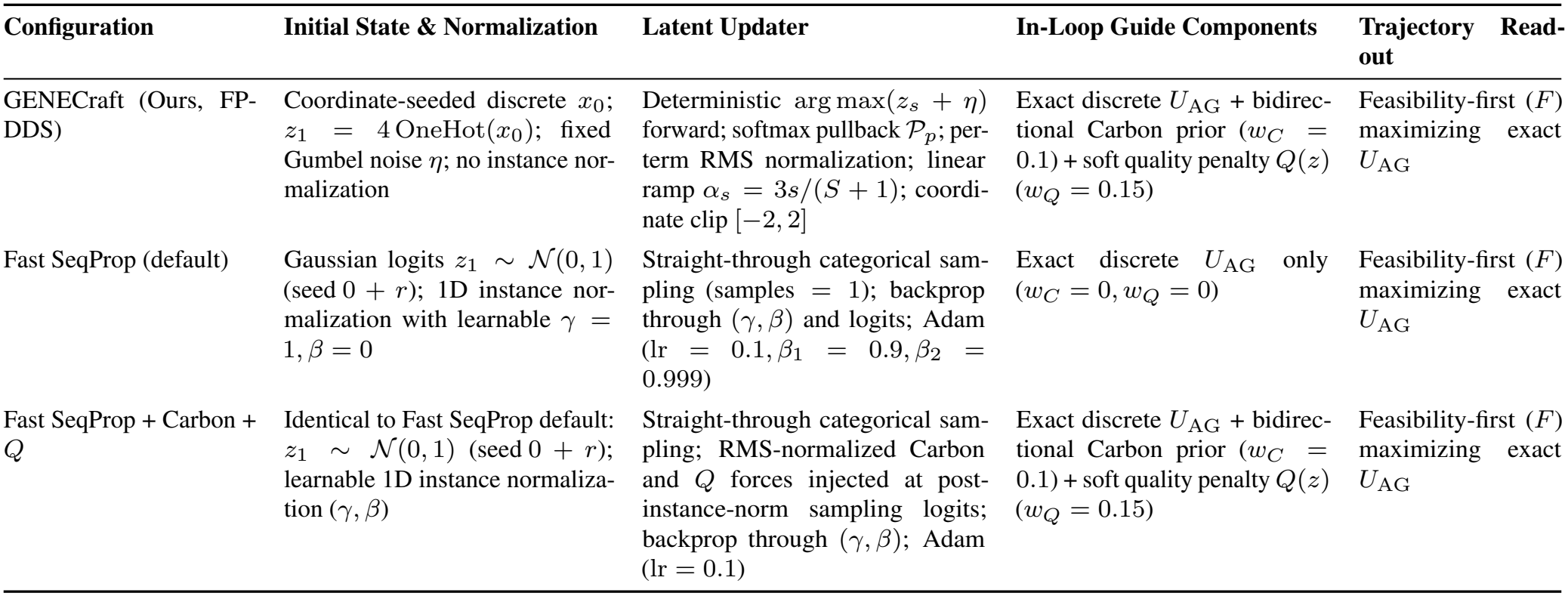
Optimizer configurations in the matched 24-locus enhancer controller study (*S* = 48, *R* = 2). Arms share utility *U*_AG_, feasibility gate *F*, feasibility-first readout, and 96 AlphaGenome backward calls per locus (Table 23); FP-DDS and Fast SeqProp + Carbon + *Q* also share Carbon and *Q* forces.

| Configuration | Initial State & Normalization | Latent Updater | In-Loop Guide Components | Trajectory Read-out |
|---|---|---|---|---|
| GENECraft (Ours, FP-DDS) | Coordinate-seeded discrete $x_0$; $z_1 = 4\,\mathrm{OneHot}(x_0)$; fixed Gumbel noise $\eta$; no instance nor-malization | Deterministic $\arg\max(z_s + \eta)$ forward; softmax pullback $\mathcal{P}_p$; per-term RMS normalization; linear ramp $\alpha_s = 3s/(S + 1)$; coordi-nate clip $[-2, 2]$ | Exact discrete $U_{\mathrm{AG}}$ + bidirec-tional Carbon prior ($w_C = 0.1$) + soft quality penalty $Q(z)$ ($w_Q = 0.15$) | Feasibility-first ($F$) maximizing exact $U_{\mathrm{AG}}$ |
| Fast SeqProp (default) | Gaussian logits $z_1 \sim \mathcal{N}(0, 1)$ (seed $0 + r$); 1D instance nor-malization with learnable $\gamma = 1, \beta = 0$ | Straight-through categorical sam-pling (samples $= 1$); backprop through $(\gamma, \beta)$ and logits; Adam ($\mathrm{lr} = 0.1, \beta_1 = 0.9, \beta_2 = 0.999$) | Exact discrete $U_{\mathrm{AG}}$ only ($w_C = 0, w_Q = 0$) | Feasibility-first ($F$) maximizing exact $U_{\mathrm{AG}}$ |
| Fast SeqProp + Carbon + $Q$ | Identical to Fast SeqProp default: $z_1 \sim \mathcal{N}(0, 1)$ (seed $0 + r$); learnable 1D instance normaliza-tion $(\gamma, \beta)$ | Straight-through categorical sam-pling; RMS-normalized Carbon and $Q$ forces injected at post-instance-norm sampling logits; backprop through $(\gamma, \beta)$; Adam ($\mathrm{lr} = 0.1$) | Exact discrete $U_{\mathrm{AG}}$ + bidirec-tional Carbon prior ($w_C = 0.1$) + soft quality penalty $Q(z)$ ($w_Q = 0.15$) | Feasibility-first ($F$) maximizing exact $U_{\mathrm{AG}}$ |

### P. Trajectory and Motif Case Protocol

Figure 4 replays a *S* = 24, *R* = 1 trajectory on the prese-lected 445-bp GM12878 *BHLHE40* locus (chr22:36017582– 36018027; 6,144-bp flanks; 4,096-bp AlphaGenome window) under default GENECRAFT weights (*w*_*C*_ = 0.1, *w*_*Q*_ = 0.15, *α*_*t*_ = 3*t/*25), locking all 25 states (within 1.33 *×* 10^*−*15^) alongside 24-step Fast SeqProp (lr 0.1) and 64-sample Evo 2 7B baselines. Bidirectional MOODS scans with JASPAR 2024 MA0464.3 (Appendix R; *p <* 10^*−*4^) detected hits at 0-based positions 134, 181, 198, 382, and 428. PCA across the 25 states captured 88.10% variance (PC1: 77.02%, PC2: 11.08%), with unit arrows at updates 4, 15, and 18 showing projected AlphaGenome and Carbon force alignment (2D cosines 0.9984, 0.6382, 0.9897; 1,780-D cosines 0.1336, *−*0.2678, 0.1725; updates 23–24 2D cosines *−*0.5788, *−*0.8865).

### Q. Locus-Level Provenance and Decomposition of Full-Cohort and 336-Locus Evaluations

Supplementary Table 27 links the full benchmark (Supplementary Table 9; *N* = 2,185) to the 336-locus perturbation analyses (Table 10). Archived perturbation records evaluate exact frozen benchmark sequences (672 parent records), whereas subset restriction isolates locus selection (Panel A). Separately, the independent de novo intervention (*w*_C_ ∈ *{*0, 0.03, 0.1, 0.3*}*; Panels B–C) runs `design_core.generate` under matched coordinate-hashed seeds and fixed Gumbel noise, improving compositional similarity and perturbation stability alongside high joint success (Table 12).

**Table 27.**
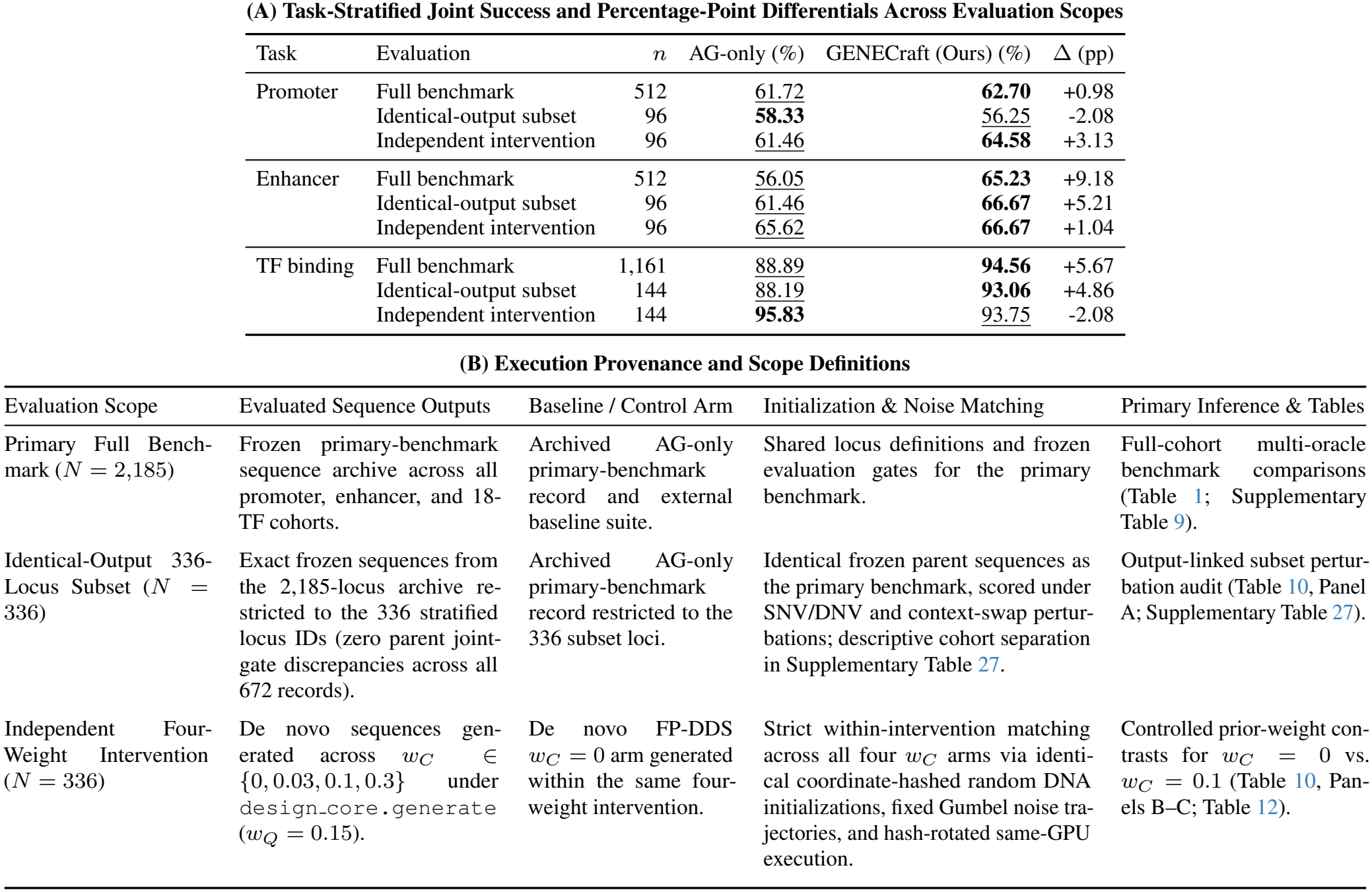
Three-layer locus-level decomposition of joint success rates and percentage-point (pp) differentials across the full cohort (*N* = 2,185; 18-TF macro-average), the output-linked 336-locus subset (Table 10, Panel A), and the independent 336-locus weight intervention (*w*_C_ = 0.1 vs. *w*_C_ = 0; Table 10, Panel B).

| (A) Task-Stratified Joint Success and Percentage-Point Differentials Across Evaluation Scopes |  |  |  |  |  |
| Task | Evaluation | *n* | AG-only (%) | GENECraft (Ours) (%) | Δ (pp) |
|---|---|---|---|---|---|
| Promoter | Full benchmark | 512 | 61.72 | **62.70** | +0.98 |
|  | Identical-output subset | 96 | **58.33** | 56.25 | -2.08 |
|  | Independent intervention | 96 | 61.46 | **64.58** | +3.13 |
| Enhancer | Full benchmark | 512 | 56.05 | **65.23** | +9.18 |
|  | Identical-output subset | 96 | 61.46 | **66.67** | +5.21 |
|  | Independent intervention | 96 | 65.62 | **66.67** | +1.04 |
| TF binding | Full benchmark | 1,161 | 88.89 | **94.56** | +5.67 |
|  | Identical-output subset | 144 | 88.19 | **93.06** | +4.86 |
|  | Independent intervention | 144 | **95.83** | 93.75 | -2.08 |
| (B) Execution Provenance and Scope Definitions |  |  |  |  |  |
| Evaluation Scope | Evaluated Sequence Outputs | Baseline / Control Arm | Initialization & Noise Matching | Primary Inference & Tables |  |
| Primary Full Bench-mark (*N* = 2,185) | Frozen primary-benchmark sequence archive across all promoter, enhancer, and 18-TF cohorts. | Archived AG-only primary-benchmark record and external baseline suite. | Shared locus definitions and frozen evaluation gates for the primary benchmark. | Full-cohort multi-oracle benchmark comparisons (Table 1; Supplementary Table 9). |  |
| Identical-Output 336-Locus Subset (*N* = 336) | Exact frozen sequences from the 2,185-locus archive re-stricted to the 336 stratified locus IDs (zero parent joint-gate discrepancies across all 672 records). | Archived AG-only primary-benchmark record restricted to the 336 subset loci. | Identical frozen parent sequences as the primary benchmark, scored under SNV/DNV and context-swap pertur-bations; descriptive cohort separation in Supplementary Table 27. | Output-linked subset pertur-bation audit (Table 10, Panel A; Supplementary Table 27). |  |
| Independent Four-Weight Intervention (*N* = 336) | De novo sequences gen-erated across *w*<sub>*C*</sub> ∈ {0, 0.03, 0.1, 0.3} under design_core.generate (*w*<sub>*Q*</sub> = 0.15). | De novo FP-DDS *w*<sub>*C*</sub> = 0 arm generated within the same four-weight intervention. | Strict within-intervention matching across all four *w*<sub>*C*</sub> arms via identi-cal coordinate-hashed random DNA initializations, fixed Gumbel noise tra-jectories, and hash-rotated same-GPU execution. | Controlled prior-weight con-trasts for *w*<sub>*C*</sub> = 0 vs. *w*<sub>*C*</sub> = 0.1 (Table 10, Pan-els B–C; Table 12). |  |

### R. Full-Cohort Motif Grammar, Repeat Structure, and Cross-Locus Diversity

Scanning 879 nonredundant JASPAR 2024 profiles [35] via MOODS [27] (both strands, 0.25 background, pseudocount 10^*−*4^, *p <* 10^*−*4^; Supplementary Table 30) across all 6,555 designs and 2,185 native targets shows GENECraft achieves the lowest motif-distribution Jensen–Shannon divergence from native DNA among the three evaluated internal design arms. GENECraft increases canonical motif presence and local stacking (*≥* 3 hits per 100 bp) while exhibiting 0.310%– 1.082% tandem repeat coverage and preserving 100.0% sequence uniqueness alongside high measured 6-mer Jaccard diversity.

Native motif-profile similarity is measured by base-2 JS divergence across 879 PWMs (including a no-hit bin) after greedy score-based overlap removal within each PWM and across shared-TF profiles. Sequence uniqueness is macro-averaged across TFs, and inter-sequence diversity is the 6-mer Jaccard distance across up to 5,000 fixed locus pairs per condition.

Our grammar audit separates global composition and constraint compliance from local target-motif multiplicity (Table 32A). Because hard QC and soft penalty *Q* govern GC balance, single-base fraction, and homopolymers without capping motif counts, local target-motif stacking (*≥* 3 hits per 100 bp) can rise alongside improved dinucleotide and 879-PWM JS divergence and high QC compliance.

### S. Supplementary Pilot Comparison with ATGC-Gen on 24 Enhancer Loci

On the 24-locus enhancer subset (Table 31), we compared GENECraft with ATGC-Gen [39] using its pretrained melanoma checkpoint (`fbd_mel_GPT.ckpt` ; 16 layers, 16 heads, 768 dimensions; cached logit error 5.72 *×* 10^*−*6^). We sampled 128 500-bp candidates per locus (temperature = 1.0, top-*k* = 4) balanced across all 47 melanoma classes, scoring the first draw and AG-ranked best-of-128 candidate with Enformer and Borzoi.

Although best-of-128 AG selection raised ATGC-Gen joint success from 0*/*24 to 11*/*24, GENECraft achieved 19*/*24 (79.17%). Both methods matched on QC (24*/*24) and chromatin-risk compliance (21*/*24), but GENECraft nearly doubled external activity success (20*/*24 vs. 11*/*24) with substantially higher Enformer and Borzoi log(1 + *x*) gains, confirming stronger functional activation rather than post-hoc filtering.

### T. Evaluation Calibration, Model Adapter Specification, and Execution Protocol

To ground functional evaluation in native genomic standards, six-TF thresholds were calibrated on 64 non-overlapping ChIP-seq peaks per factor in `EFO:0002784` over 4,096 bp (`guidance_objective=peak_mean`, `guidance peak width bp=256`, `score_center_width_bp=256` ; Table 28). Complementing these targets, enhancer risk cutoffs (Table 7) derive from Enformer and Borzoi predictions across *n* = 50 DeepMEL2 enhancers (`Genomic_MEL_regions.fa` ) via linear quantile interpolation (*h* = (*n −* 1)*q*), bounding CAGE peak (*P*_99_), H3K4me3 mean (*P*_80_), and the H3K4me1*/*(H3K4me3 + 10^*−*6^) ratio (*P*_10_).

**Table 28.** Natural ChIP-seq calibration quantiles (*n* = 64 loci per factor, ontology `EFO:0002784`, 4,096 bp window; `guidance_objective=peak_mean`, `guidance_peak_width_bp=256`, `score_center_width_bp=256` ) defining the common AG evaluation gate (*p*_50_).

| TF | *n* | *p*<sub>50</sub> | *p*<sub>80</sub> |
|---|---|---|---|
| BCL11A | 64 | 8.71875 | 10.7375 |
| BHLHE40 | 64 | 13.5625 | 21.15 |
| IKZF1 | 64 | 6.3515625 | 9.4375 |
| NFIC | 64 | 7.3984375 | 9.19375 |
| TBP | 64 | 7.9375 | 11.9875 |
| USF2 | 64 | 14.75 | 21.85 |

Enhancer cross-evaluation embeds the central 500 bp candidate into native GRCh38 context for Enformer (393,216 bp input, 128 bp bins) and Borzoi (524,288 bp input, 32 bp bins), starting outputs at input start + (input *−* output)*/*2 (Table 29). DNase and H3K27ac average overlapping central 500 bp bins, CAGE uses the peak bin-wise mean, and chromatin specificity spans 2,048 bp as H3K4me1*/*(H3K4me3 + 10^*−*6^).

**Table 29.**
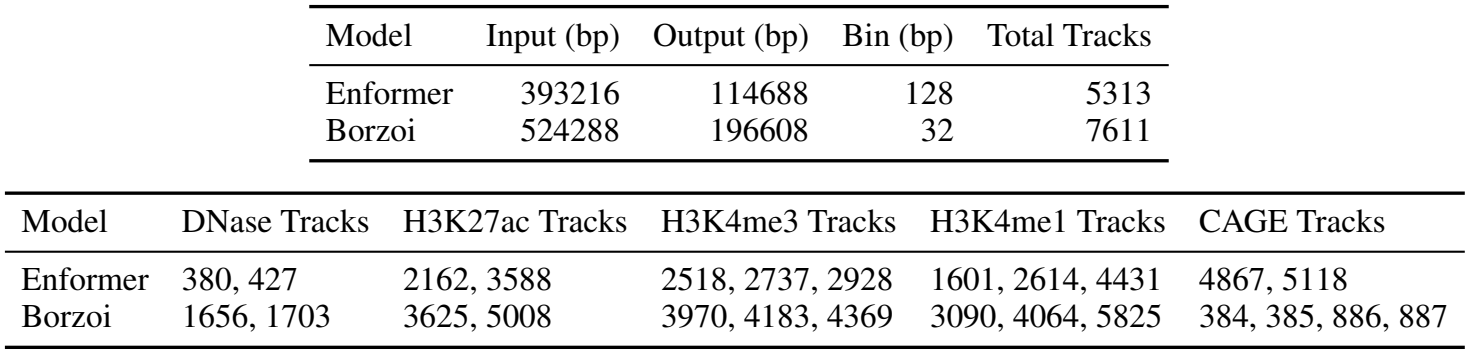
Genomic window geometry and zero-based melanocyte track identifiers for in-situ 500 bp enhancer evaluation in Enformer and Borzoi.

**Table 30.** Full-cohort motif grammar, tandem-repeat coverage, and cross-locus diversity across 2,185 native loci and 6,555 designs (equal weighting across 18 TFs; target-motif metrics span 17 JASPAR 2024 TFs across 1,139 loci).

| Task | Method | Motif-profile JS | Tandem repeats (%) | Exact / RC unique (%) | 6-mer distance |
|---|---|---|---|---|---|
| Promoter | Native | 0.0000 | 1.853 | 98.83 | 0.9394 |
|  | AG-only | 0.9148 | 0.442 | 99.80 | 0.9425 |
|  | Sequential AG–Carbon | 0.9089 | 0.665 | 99.80 | 0.9436 |
|  | **GENECraft (Ours)** | 0.8943 | 1.082 | 100.00 | 0.9405 |
| Enhancer | Native | 0.0000 | 0.564 | 100.00 | 0.9157 |
|  | AG-only | 0.8388 | 0.114 | 100.00 | 0.8668 |
|  | Sequential AG–Carbon | 0.8362 | 0.192 | 100.00 | 0.8660 |
|  | **GENECraft (Ours)** | 0.8353 | 0.924 | 100.00 | 0.8351 |
| TF binding | Native | 0.0000 | 0.781 | 100.00 | 0.9459 |
|  | AG-only | 0.8099 | 0.160 | 100.00 | 0.9097 |
|  | Sequential AG–Carbon | 0.8087 | 0.200 | 100.00 | 0.9099 |
|  | **GENECraft (Ours)** | 0.7828 | 0.310 | 100.00 | 0.9131 |
| Method (17 TFs) | Target-motif presence (%) | Hits / 100 bp | Stacking (%) |  |  |
| Native | 38.40 | 0.1612 | 0.90 |  |  |
| AG-only | 57.96 | 0.4658 | 8.48 |  |  |
| Sequential AG–Carbon | 58.37 | 0.4764 | 9.61 |  |  |
| **GENECraft (Ours)** | 64.13 | 0.5917 | 14.81 |  |  |
| *p* = 10<sup>−3</sup> sensitivity | AG-only | Sequential AG–Carbon | GENECraft (Ours) |  |  |
| Promoter motif-profile JS | 0.6830 | 0.6677 | 0.6439 |  |  |
| Enhancer motif-profile JS | 0.4756 | 0.4710 | 0.4583 |  |  |
| TF binding motif-profile JS | 0.5670 | 0.5625 | 0.5178 |  |  |
| Target-motif presence (%) | 79.62 | 80.31 | 84.63 |  |  |

**Table 31.**
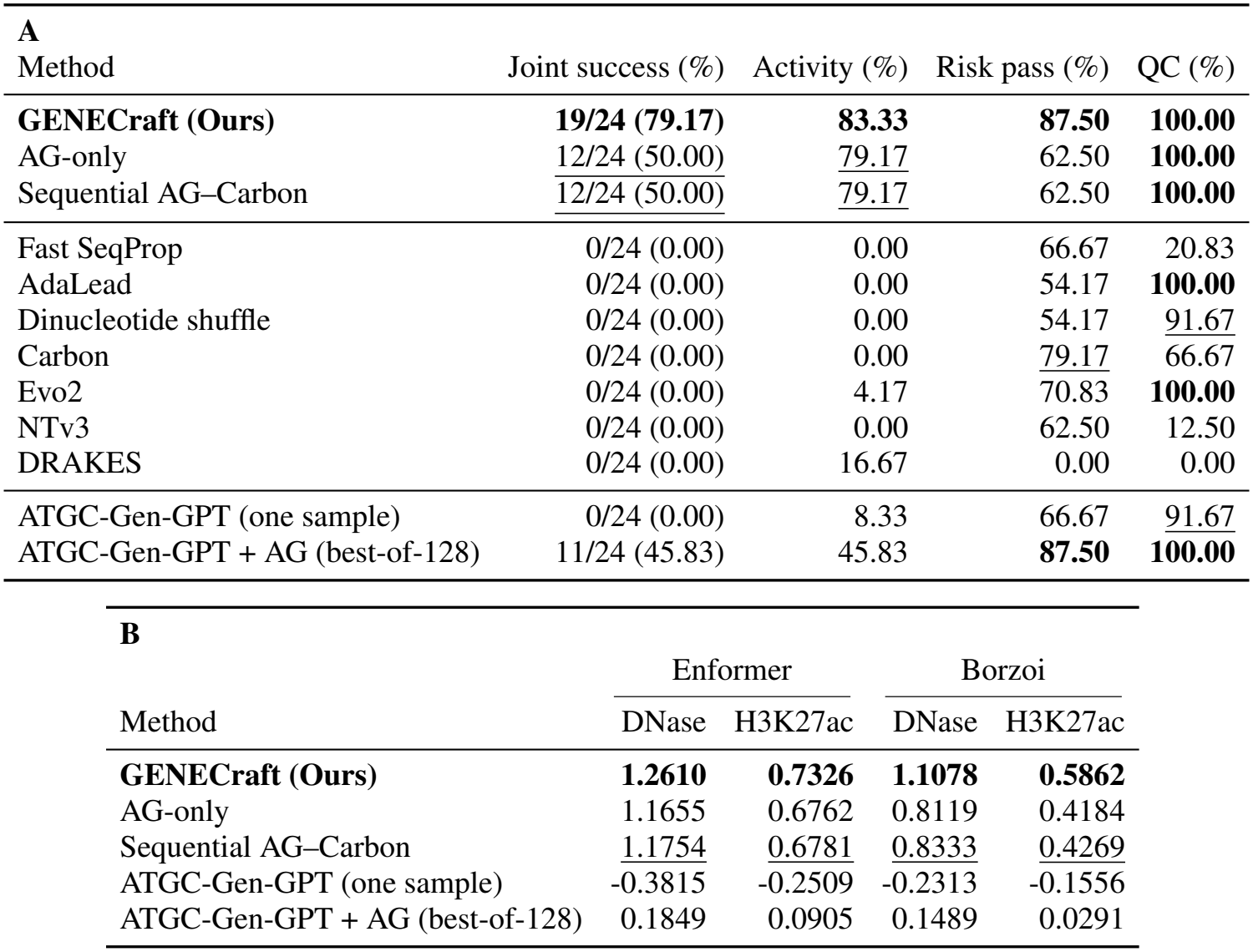
24-locus enhancer pilot comparing GENECraft, ablations, external baselines, and ATGC-Gen [39] (first draw and best-of-128 AG-selected). Panel A reports joint/activity success and risk/QC compliance across 12 methods (*n* = 24); Panel B reports mean native-relative log(1 + *x*) Enformer and Borzoi gains.

| A |  |  |  |  |
|---|---|---|---|---|
| Method | Joint success (%) | Activity (%) | Risk pass (%) | QC (%) |
| **GENECraft (Ours)** | **19/24 (79.17)** | **83.33** | **87.50** | **100.00** |
| AG-only | 12/24 (50.00) | 79.17 | 62.50 | **100.00** |
| Sequential AG–Carbon | 12/24 (50.00) | 79.17 | 62.50 | **100.00** |
| Fast SeqProp | 0/24 (0.00) | 0.00 | 66.67 | 20.83 |
| AdaLead | 0/24 (0.00) | 0.00 | 54.17 | **100.00** |
| Dinucleotide shuffle | 0/24 (0.00) | 0.00 | 54.17 | 91.67 |
| Carbon | 0/24 (0.00) | 0.00 | 79.17 | 66.67 |
| Evo2 | 0/24 (0.00) | 4.17 | 70.83 | **100.00** |
| NTv3 | 0/24 (0.00) | 0.00 | 62.50 | 12.50 |
| DRAKES | 0/24 (0.00) | 16.67 | 0.00 | 0.00 |
| ATGC-Gen-GPT (one sample) | 0/24 (0.00) | 8.33 | 66.67 | 91.67 |
| ATGC-Gen-GPT + AG (best-of-128) | 11/24 (45.83) | 45.83 | **87.50** | **100.00** |
| **B** |  |  |  |  |
|  | Enformer |  | Borzoi |  |
| Method | DNase | H3K27ac | DNase | H3K27ac |
| **GENECraft (Ours)** | **1.2610** | **0.7326** | **1.1078** | **0.5862** |
| AG-only | 1.1655 | 0.6762 | 0.8119 | 0.4184 |
| Sequential AG–Carbon | 1.1754 | 0.6781 | 0.8333 | 0.4269 |
| ATGC-Gen-GPT (one sample) | -0.3815 | -0.2509 | -0.2313 | -0.1556 |
| ATGC-Gen-GPT + AG (best-of-128) | 0.1849 | 0.0905 | 0.1489 | 0.0291 |

**Table 32.** Paired locus-level motif co-occurrence and enhancer gate transitions. **A**, Target-motif stacking (*≥* 3 instances per 100 bp) co-occurrence with improved dinucleotide JS, 879-motif JS, and hard-QC compliance across *n* = 1, 139 loci (17 TFs) versus AG-only and AG→Carbon. **B**, Gate-level pass transitions between primary and repeat FP-DDS runs (*n* = 24 enhancers).

| **(A) Motif-grammar co-occurrence** (*n* = 1,139; 17 TFs) |  |  |  |  |  |
| Reference | New stacking | Lower dinucleotide JS | Lower motif-profile JS | Hard QC pass |  |
|---|---|---|---|---|---|
| AG-only | 157 | 130 | 102 | 155 |  |
| Sequential AG→Carbon | 154 | 122 | 96 | 152 |  |
| **(B) Enhancer gate transitions** (*n* = 24) |  |  |  |  |  |
| Endpoint | Reference pass | Repeat pass | Both pass | Reference-only | Repeat-only |
| joint | 19 | 17 | 15 | 4 | 2 |
| activity | 20 | 22 | 20 | 0 | 2 |
| risk | 21 | 17 | 16 | 5 | 1 |
| QC | 24 | 24 | 24 | 0 | 0 |
| AG | 22 | 22 | 22 | 0 | 0 |
| Risk gate | Reference fail | Repeat fail | Lost compliance | Gained compliance |  |
| Enformer CAGE | 0 | 1 | 1 | 0 |  |
| Enformer H3K4me3 | 2 | 3 | 1 | 0 |  |
| Enformer H3K4me1/H3K4me3 | 2 | 2 | 0 | 0 |  |
| Borzoi CAGE | 3 | 4 | 4 | 3 |  |
| Borzoi H3K4me3 | 2 | 2 | 0 | 0 |  |
| Borzoi H3K4me1/H3K4me3 | 2 | 2 | 0 | 0 |  |

Table 33 summarizes the H20 execution environment and model provenance. Carbon 3B executes in GPU `bfloat16`, whereas controller updates and sampling run in NumPy `float64` . Initial sequences use `np.default_rng` seeded from the first 8 little-endian bytes of SHA256(`crossed-initial-v1|` + case id), with deterministic Gumbel noise *−* log(*−* log(*U* )) indexed by (case id, seed, restart).

**Table 33.** Model artifact provenance and software stack for the H20 supplementary analysis environment.

| Model | Artifact Identity | Supplementary Runtime Stack |
|---|---|---|
| AlphaGenome | manifest.ocdbt: 136ce6bd618b; _METADATA: df68f8d9ded2 | JAX/JAXlib 0.10.2; NumPy 2.2.2; Transformers 5.20.2 |
| Carbon 3B | carbon_checkpoint_sha256_manifest.json | GPU bfloat16; Transformers 5.20.2 |
| Enformer | saved_model.pb: c8f0af88f362; variables: 6a64f5fdb08b | TensorFlow 2.15.1; TF Hub 0.15.0; NumPy 1.26.4 |
| Borzoi | johahi/borzoi-replicate-0; 521db1033727 | Torch 2.6.0+cu118; Transformers 4.57.6; borzoi-pytorch 0.5.1 |
| Evo2 7B | bda0089f9258; SHA256 c66645929dc1 | bfloat16 |
| ATGC-Gen | fbd_mel_GPT.ckpt; 51a75d724306 | FP32; enhancer pilot |
| H20 | Python 3.11.11; driver 580.105.08 | Controller logits: NumPy float64 |

GRCh38 mapping confirms zero overlap (0*/*0; minimum distance 567,263 bp) between the 50 calibration enhancers and all evaluation cohorts across 500 bp targets and oracle contexts up to 524,288 bp (Supplementary Table 34). After the early *P*_95_ → *P*_99_ CAGE update on preliminary 8-locus and 50-design runs, the frozen *P*_99_ threshold governed all benchmark and ablation evaluations.

**Table 34.**
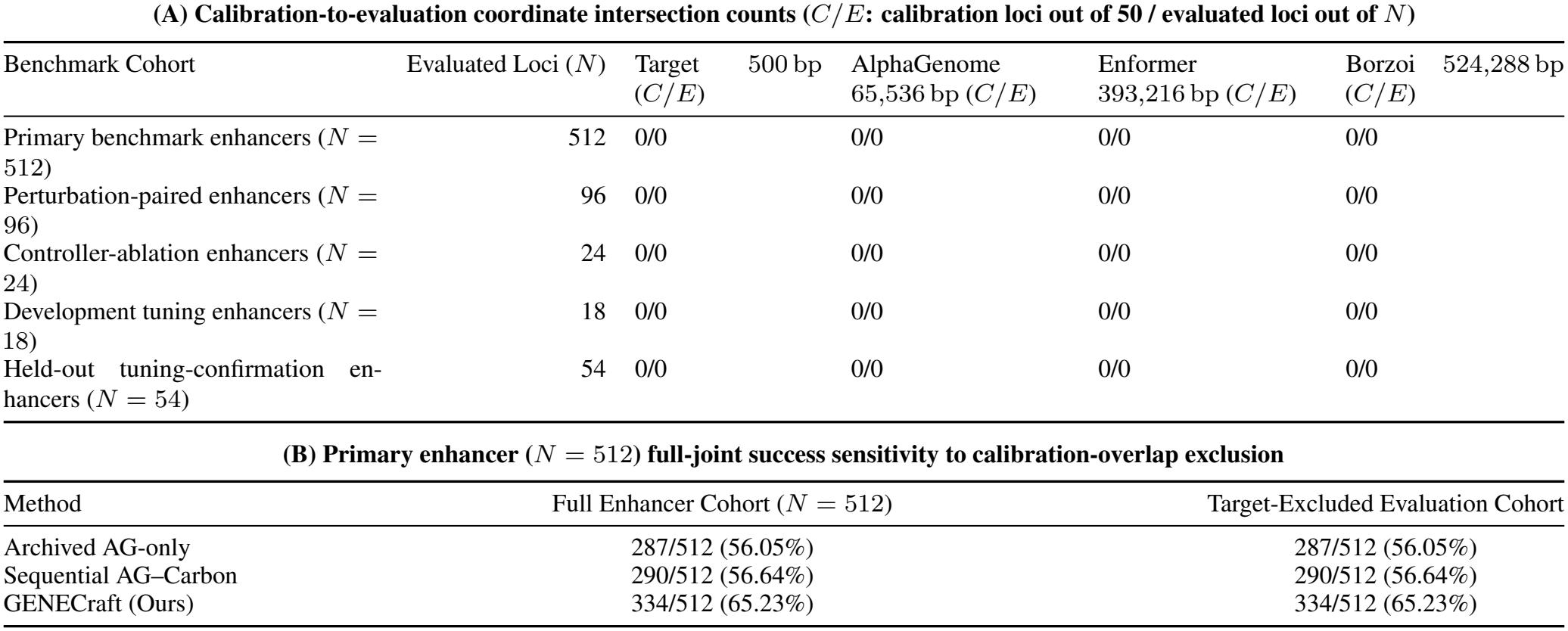
Coordinate overlap and exclusion sensitivity for 50 off-target calibration loci (0-based half-open GRCh38). Panel A reports intersection counts *C/E* (*C* calibration, *E* cohort loci) across 500 bp target and full 65,536 bp AlphaGenome, 393,216 bp Enformer, and 524,288 bp Borzoi windows. Panel B reports 512-enhancer joint success after overlap exclusion.

| **(A) Calibration-to-evaluation coordinate intersection counts (*C*/*E*: calibration loci out of 50 / evaluated loci out of *N*)** |  |  |  |  |  |
|---|---|---|---|---|---|
| Benchmark Cohort | Evaluated Loci (*N*) | Target 500 bp (*C*/*E*) | AlphaGenome 65,536 bp (*C*/*E*) | Enformer 393,216 bp (*C*/*E*) | Borzoi 524,288 bp (*C*/*E*) |
| Primary benchmark enhancers (*N* = 512) | 512 | 0/0 | 0/0 | 0/0 | 0/0 |
| Perturbation-paired enhancers (*N* = 96) | 96 | 0/0 | 0/0 | 0/0 | 0/0 |
| Controller-ablation enhancers (*N* = 24) | 24 | 0/0 | 0/0 | 0/0 | 0/0 |
| Development tuning enhancers (*N* = 18) | 18 | 0/0 | 0/0 | 0/0 | 0/0 |
| Held-out tuning-confirmation en-hancers (*N* = 54) | 54 | 0/0 | 0/0 | 0/0 | 0/0 |
| **(B) Primary enhancer (*N* = 512) full-joint success sensitivity to calibration-overlap exclusion** |  |  |  |  |  |
| Method | Full Enhancer Cohort (*N* = 512) | Target-Excluded Evaluation Cohort |  |  |  |
| Archived AG-only | 287/512 (56.05%) | 287/512 (56.05%) |  |  |  |
| Sequential AG–Carbon | 290/512 (56.64%) | 290/512 (56.64%) |  |  |  |
| GENECraft (Ours) | 334/512 (65.23%) | 334/512 (65.23%) |  |  |  |

## Footnotes

1 The CAGE ceiling prevents focal transcription initiation without abolishing eRNA transcription, while H3K4me3 suppression with H3K4me1 enrichment enforces canonical enhancer signatures [2, 19].

